# Tissue-specific mutagenesis from endogenous guanine damage is suppressed by Polκ and DNA repair

**DOI:** 10.1101/2025.03.12.642750

**Authors:** Yang Jiang, Moritz Przybilla, Linda Bakker, Foster C. Jacobs, Dylan Mckeon, Roxanne van der Sluijs, Juliëtte Wezenbeek, Joeri van Strien, Jeroen Willems, Alexander E. E. Verkennis, Jamie Barnett, Adrian Baez Ortega, Federico Abascal, Peter W. Villalta, Puck Knipscheer, Inigo Martincorena, Silvia Balbo, Juan Garaycoechea

## Abstract

Knowledge of mutational patterns has expanded significantly, but a persistent challenge is to link these complex patterns to the underlying molecular mechanism or source of DNA damage, especially when that damage is endogenous and not driven by environmental mutagens. Technological advances now allow us to catalogue mutation across tissues or even closely related cell types, but the results are largely descriptive until we identify the endogenous sources of mutation and how they differ across tissues. Here, we combine mouse genetics, advanced sequencing, biochemistry and mass spectrometry to provide a detailed mechanistic understanding of endogenous mutagenesis. We reveal that endogenous guanine adducts are significant drivers of tissue-specific mutagenesis, while the interwoven actions of DNA polymerase Polκ and DNA repair mechanisms are pivotal in mitigating mutagenesis. For the first time, we use untargeted DNA adductomics to characterize new sources of endogenous DNA damage. Our novel approach to understanding endogenous mutational landscapes reveals previously unseen mutational processes and points to vast potential for new discoveries.

## Introduction

The integrity of the genome is constantly under threat. This threat comes from external agents that damage DNA (e.g. UV radiation), and from internal sources that are even more insidious. Normal cellular processes generate a range of reactive by-products that attack DNA and cause endogenous DNA damage. Known sources include spontaneous hydrolysis, reactive oxygen species (ROS) and simple aldehydes which produce chemically distinct DNA lesions^1–3^. Despite the prevalence of endogenous DNA damage, we have only a crude understanding of different DNA lesions and damage sources. This is because direct detection of endogenous DNA lesions is extremely challenging, due to their low abundance, rapid repair, and similarity to unmodified bases. The intrinsic instability of DNA led to the discovery that cells must actively counteract this damage for DNA to fulfill its function as the genetic material of the cell^1^.

Cells have two major pathways to deal with DNA damage, and they differ in fidelity. The first route is DNA repair, which encompasses a toolkit of efficient repair mechanisms that excise or correct DNA lesions, each tailored to specific types of damage. However, some lesions may evade repair or be encountered during DNA replication. The second pathway, known as DNA damage tolerance, allows cells to bypass DNA lesions, mostly during DNA replication to maintain fork progression^4^. A major route of DNA damage tolerance is translesion synthesis (TLS), which relies on diverse specialized DNA polymerases that can synthesize DNA over damaged bases^5^. These TLS polymerases can accommodate various types of distorted DNA because they have bigger active sites. TLS polymerases can be relatively error-free depending on the lesion, but they are more often error-prone, leading to mutations. Therefore, DNA damage tolerance comes at the cost of increased mutation but protects cells from more severe genomic instability, such as DNA breaks arising from failure to replicate past DNA lesions.

This interplay between DNA damage, DNA repair and DNA damage tolerance is essential for maintaining genomic integrity and is a key determinant of mutagenesis. A classic example of this interplay is DNA damage induced by UV radiation, largely repaired by nucleotide excision repair (NER) which removes bulky DNA adducts and helix-distorting lesions^6^. In parallel, the TLS polymerase Polη performs efficient and error-free bypass (or tolerance) of UV damage^7^. Mutations in either NER (*XPA-XPG*) or Polη (*XPV*) cause increased mutagenesis in response to UV, leading to skin cancer and the human syndrome Xeroderma pigmentosum^8^. Therefore, mutagenic outcomes depend on the nature of the DNA lesion, but more importantly on how the cell deals with the damage.

The recent explosion of genome sequencing data has made it possible to characterize mutational patterns or ‘signatures’ across thousands of genomes^9,10^. Certain mutational signatures can be linked to exogenous exposures like UV radiation (single base substitution signature 7, or SBS7^8^) or known endogenous DNA damage (e.g. deamination, SBS1^11^, SBS2, SBS13^12^; or oxidation, SBS18^13^, SBS36^14^). Interestingly, some signatures only occur in a subset of tissues, presumably due to organ-specific cellular physiology. Many mutational signatures are suspected to be caused by endogenous sources, but the identity of the lesions are unknown. For example, NER deficiency leads to SBS8 mutations in tissues not exposed to UV radiation, but the endogenous damage driving these mutations is a mystery^15,16^. Similarly, SBS19 mutations in blood stem cells are driven by persistent endogenous lesions of unknown origin^17^.

Mutational signatures are complex patterns arising from poorly understood interactions among DNA damage, repair, and damage tolerance. Here we untangle this complex interplay in mammalian tissues, with the ultimate aim of uncovering the chemical nature of novel sources of endogenous DNA damage. To tackle this fundamental question, we characterize somatic mutations in mice lacking the TLS polymerase Polκ. We combine organoid culture and recent advances in genome sequencing to determine the contribution of Polκ to the genome-wide somatic mutational landscape of mouse tissues. We discover that Polκ suppresses a novel tissue-specific mutational signature, driven by endogenous guanine adducts. We then combine mouse genetics, adductomics, and biochemistry to demonstrate how both bulky and non-bulky guanine lesions contribute to mutagenesis. Finally, we use a recently developed untargeted DNA adductomics method to uncover novel endogenous lesions. Together, our findings show that Polκ and DNA repair cooperate to limit mutagenesis in tissues, and shed light on the elusive nature of endogenous DNA damage.

## Results

### Characterizing somatic mutations in mouse tissues

Polκ is the most conserved TLS polymerase, with orthologs in bacteria and archaea, and it is able to bypass *in vitro* various bulky and non-bulky lesions^18^. To determine the role of Polκ in shaping mutational landscapes across different organs, we comprehensively assessed somatic mutations across a panel of mammalian tissues. We analyzed tissues from aged (18-month-old) wild type and *Polk-/-* mice using either *in vitro* expansion of single cells or NanoSeq^19^ (**Fig. 1a**). We expanded single-cell clones from bone marrow progenitors, and used organoid culture to expand single stem cells from the small intestine, stomach, lung airways, and liver cholangiocytes, as done previously^20,21^. Clones were expanded for 3-4 weeks and subjected to whole-genome sequencing; in total 32 genomes were sequenced at around 25x depth together with a germline reference (**Fig. 1a**). We used a combination of Strelka2 and Mutect2 to call high-confidence somatic mutations, defined as clonal mutations present in the original cell that are shared by its progeny and have a variant allele frequency (VAF) centered around 0.5. Subclonal mutations that occur during *in vitro* culture have low allele frequencies and are removed from the analysis. Samples without a distinct peak around VAF 0.5 were considered non-clonal and excluded from downstream analysis (**Supplementary Fig. 1a**). Sequencing of single-cell derived clones provides whole-genome coverage data but is limited to cell types that can be expanded *in vitro*. Therefore, in parallel, we used NanoSeq to interrogate wild type and *Polk-/-* tissues without the need for *in vitro* culture. NanoSeq is a highly sensitive single-molecule technique, based on duplex sequencing, that allows detection of rare somatic mutations in a mixed population of cells^19^. We applied NanoSeq to bulk DNA from the kidney, adrenal gland (among the tissues with highest Polκ expression), as well as lung and liver (to draw comparisons between the two methods). Overall, we found good correlation in mutation burden estimates for single-base substitutions (SBSs), doublet base substitutions (DBSs), and insertions/deletions (indels) between clonal expansion of single cells and NanoSeq (**Supplementary Fig. 1b**), with NanoSeq having a higher burden in line with previous work^19^.

**Figure 1.**
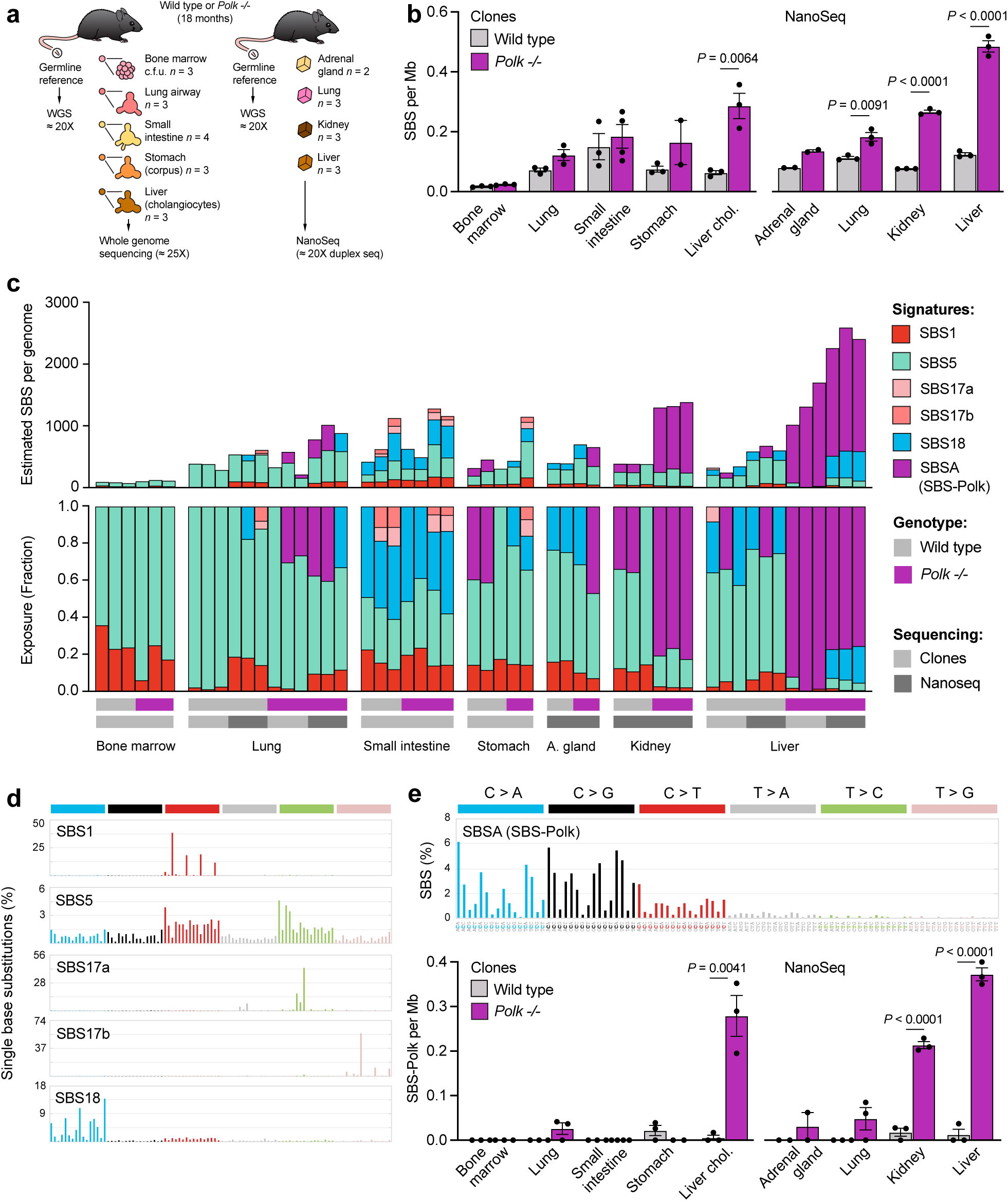
Polk suppresses a novel, tissue-specific mutational signature. **a)** Experimental layout to uncover the somatic mutational landscape of mouse tissues: single cells from aged mice were clonally amplified and subjected to whole genome sequencing, or bulk tissue was sequenced with NanoSeq to identify rare somatic mutations in bulk DNA samples. **b)** Burden of single-base substitutions (SBSs) per genome (*P* calculated by an unpaired *t* test). **c)** Assignment of mutational signatures using SigProfiler toolkit. Stacked bar plots showing estimated number (top) and proportion (bottom) of each mutational signature in individual clones and Nanoseq samples. **d)** Pattern of known COSMIC mutational signatures, 96-classes of SBSs considering the six mutation types but also the bases immediately 5’ and 3’ of the mutated base. **e)** Top, pattern of a novel mutational signature (SBSA, or SBS-Polk), which explains most mutations in *Polk-/-* kidney and liver samples. Bottom, quantification of SBS-Polk mutations in wild type and *Polk-/-* samples (*P* calculated by unpaired *t* tests).

### Polκ suppresses a novel tissue-specific mutational signature

We first assessed the SBS burden in different tissues. In wild type mice, we found that the SBS burden varies across tissues, being highest in the small intestine and lowest in bone marrow progenitors (**Fig. 1b**). Our trends are consistent with previous observations^19–22^. However, *Polk-/-* mice show a strikingly different pattern (**Fig. 1b**). We observe a 4-fold increase in the burden of SBSs in the kidney and liver, while the SBS burden is essentially unchanged in the small intestine and bone marrow progenitors. This shows that Polκ normally suppresses point mutations and that the increased mutagenesis is tissue-specific. We find no obvious correlation between mutation burden and the expression level of Polκ or other TLS polymerases (**Supplementary Fig. 1c**). These results indicate that differences in TLS usage cannot explain the tissue-specific mutagenesis we observe, and that it is likely due to different damage burden across tissues.

To look further into the mutation patterns, we sorted SBSs into six classes, referring to the pyrimidine of the Watson-Crick base pair as the mutated base (**Supplementary Fig. 2a**). This analysis reveals that increased mutagenesis in *Polk-/-* liver cholangiocytes, bulk liver, and kidney is largely driven by C>A and C>G changes, and to a lesser extent by C>T and T>A mutations (**Supplementary Fig. 2a**). These six SBS classes can be further expanded by considering the nucleotide context (i.e., the bases immediately 5’ and 3’ of the mutated base). This 96-context classification allowed us to further refine mutation types and reveals considerable heterogeneity in the somatic mutational landscape of wild type mouse tissues, mirroring observations in human cancer^9,10^ and normal human tissues^23^ (**Supplementary Fig. 2b**).

To systematically explore the differences in mutational landscapes, we performed *de novo* mutational signature extraction using non-negative matrix factorization (NMF) with SigProfilerExtractor, yielding three *de novo* signatures (SBSA-C, **Supplementary Fig. 3a**). We obtained comparable signatures using mSigHdp, a non-NMF approach using a hierarchical Dirichlet process model (HDP)^24,25^ (**Supplementary Fig. 3b**). SigProfiler signatures SBSB and SBSC could be reconstructed (i.e. Cosine similarity > 0.92) by a combination of known signatures from the Catalogue of Somatic Mutations in Cancer (COSMIC). These signatures included: SBS1, caused by deamination of 5-methylcytosine at CpG sites^11^; SBS5, a ubiquitous signature of unknown origin which accumulates over time independently of cell division^19,26,27^; SBS17, induced by 5-fluorouracil and an unknown endogenous driver^28^, and SBS18, caused by the mutagenic bypass of 8oxo-guanine, a lesion linked to reactive oxygen species^13^ (**Fig. 1c, d**).

The SBSA signature, on the other hand, is characterized by C>A, C>G and C>T changes and bears little similarity to known COSMIC signatures (**Fig. 1c,e, Supplementary Fig. 3c**). SBSA was reconstructed by SigProfiler with a combination of COSMIC signatures SBS4 (C>A mutations induced by tobacco smoking) and SBS39 (C>G mutations, unknown cause), albeit with limited confidence (Cosine sim. 0.845) (**Supplementary Fig. 3d**). SBS39 is one of the few C>G rich signatures in the COSMIC database, is mostly found in medulloblastoma and breast cancer, and its driver is unknown. Upon closer inspection, we noted that SBS39 completely lacks transcriptional-strand bias, a prominent feature of both SBS4 and SBSA (**Supplementary Fig. 3e**), which we discuss in more detail below. Due to these differences in mutational strand-asymmetries, and the fact that the mice were not exposed to tobacco, we determined SBSA should not be decomposed further into SBS4 and SBS39 but defined SBSA as a novel signature. Importantly, the SBSA signature was largely responsible for the increase of mutation in *Polk-/-* bulk kidney, liver and liver cholangiocytes (**Fig. 1c,e**). The mutational pattern was almost identical between *Polk-/-* kidney and liver cholangiocytes (Cosine sim. 0.93), but mutations in bulk liver had a larger C>A component compared to liver cholangiocytes (**Supplementary Fig. 2a, b**). As mentioned previously, NanoSeq provides somatic mutations in single DNA molecules from all cells in the tissue of interest. In liver, around 60% of cells are hepatocytes whereas cholangiocytes, which were grown into clones sequenced with WGS, only make up around 3%. In line with this, lung samples had even more distinct patterns, reflecting differences between clones from bronchial epithelium and bulk lung tissue (also containing alveolar epithelium and immune cells amongst other cells)^29^.

In summary, we characterized somatic mutations in mouse tissues using two complementary approaches. We find that the TLS polymerase Polκ suppresses a novel, tissue-specific SBS mutational signature; hereafter, we refer to SBSA as SBS-Polk. The signature is driven by endogenous DNA damage, and our results imply that Polκ predominantly performs error-free bypass of this damage, particularly in the liver and kidney.

### The landscape of doublet base substitutions and indels in mouse tissues

Having uncovered a novel genome-wide SBS mutational signature, we next turned our attention to other types of mutations. DBSs are exceedingly rare across mouse tissues, in the range of 0-20 DBSs/genome, but we find a higher burden of DBSs in *Polk-/-* bulk kidney and liver (**Fig. 2a**). Although we can visualize DBS mutation types using the COSMIC DBS78 classification, we were unable to extract mutational signatures due to the low numbers of mutations detected. In wild type livers, DBSs are dominated by CC>AA (or GG>TT) changes but *Polk-/-* livers display a wider spectrum of changes (**Fig. 2b, Supplementary Fig. 4**). Because a likely cause of DBSs is the mutagenic bypass of tandem base damage (e.g. intrastrand crosslinks), our results suggest that Polκ suppresses endogenous DBSs by contributing to error-free bypass of endogenous tandem lesions.

**Figure 2.**
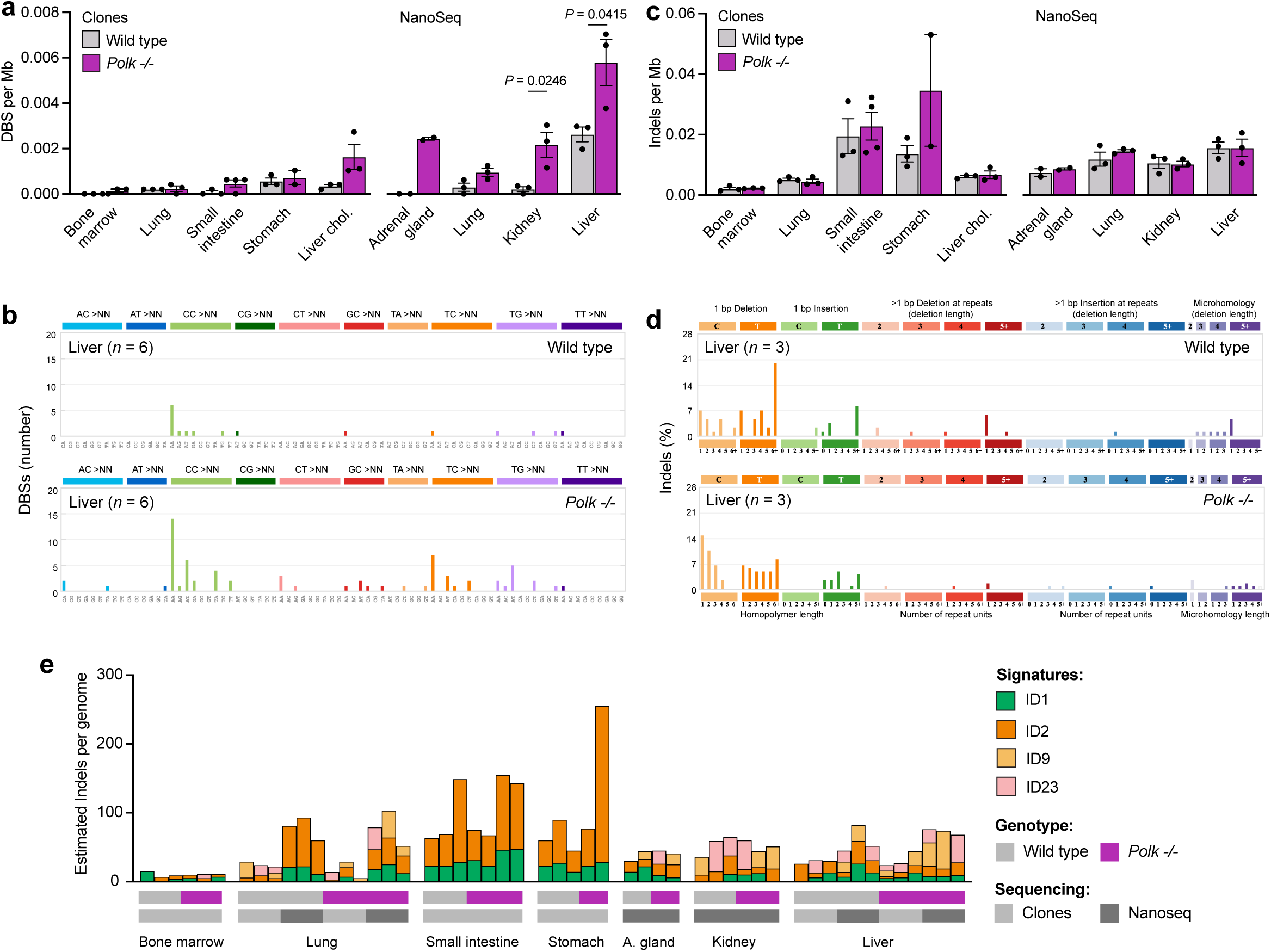
Polk protects tissues from doublet base substitutions (DBSs). **a)** Burden of DBSs per megabase (Mb) (*P* calculated by unpaired *t* tests). **d)** Pattern of DBSs, following the 78-type classification from the COSMIC database. Due to the low number of DBSs per sample, the data from liver clones (n = 3) and NanoSeq (n = 3) was combined. **c)** Burden of insertions/deletions (indels) per genome (no significant changes detected). **d)** Pattern of insertions and deletions in bulk liver DNA, following the 83-type classification from the COSMIC database. **e)** Extraction and assignment of indel mutational signatures using SigProfiler Extractor. Stacked bar plots showing estimated number of each mutational signature in individual clones and NanoSeq samples.

Focusing on the burden of indels, we observe similar burdens in wild type and *Polk-/-* mice across tissues, with the highest indel burden in the small intestine and stomach (**Fig. 2c**). We assessed the pattern of indels using the COSMIC ID83 classification, which considers size, nucleotides affected, and presence on repetitive and/or microhomology regions, and did not find obvious differences (**Fig. 2d, Supplementary Fig. 5**). Despite low numbers of indels, SigProfiler extracted two *de novo* indel signatures which were reconstructed by a combination of known COSMIC signatures ID1, ID2 (both polymerase slippage during replication), ID9 (unknown cause) and ID23 (aristolochic acid exposure) (**Fig. 2e**). The indel mutational landscape in the small intestine and stomach, both actively dividing epithelia, is dominated by ID1 and ID2, consistent with DNA replication driving these indels^9^. Importantly, the liver of *Polk-/-* mice carries a low number of indels indistinguishable from wild type controls (**Fig. 2c**) and with a similar pattern to wild type livers (**Fig. 2d, e**), indicating the mutagenesis in *Polk-/-* livers is restricted to substitutions.

### SBS-Polk mutations are characterized by transcriptional-strand bias (TSB)

The most striking observation in our mutation analysis is the presence of a novel mutational signature in *Polk-/-* kidney and liver. To elucidate the topographical characteristics of SBS-Polk mutations across the mouse genome^30,31^, we characterized the SBS-Polk signature using SigProfilerTopography^31^. First, we considered the relationship with DNA replication and detected an enrichment of SBS-Polk mutations in late vs early-replicating regions (**Fig. 3a**). This is also observed for several other mutational signatures^30–33^ and may be explained in part by the preferential use of error-prone TLS over error-free template switching during late S phase^34–36^. Mutational processes that are coupled to replication (e.g. mismatch repair, Polδ or Polε mutations) show replication-strand bias^31,37,38^. In contrast, SBS-Polk mutations have no replication strand asymmetry (**Fig. 3b**), suggesting that the TLS factors responsible for SBS-Polk mutations are not preferentially associated with either the lagging or leading strands. As it is the case with most mutational signatures, SBS-Polk mutations are depleted in genic regions (**Fig. 3b**).

**Figure 3.**
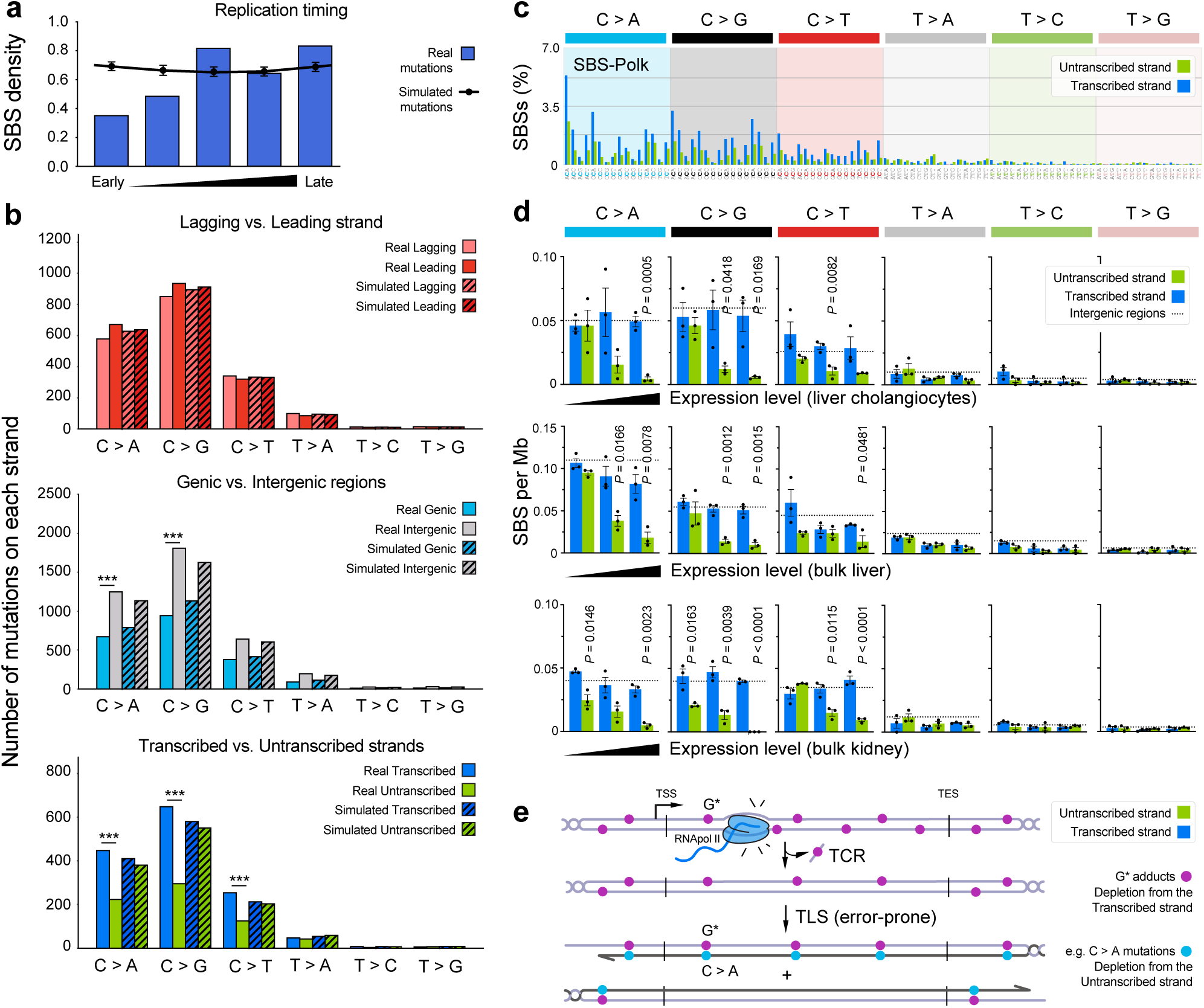
The topography of SBS-Polk mutations is characterized by transcriptional-strand bias. **a)** Normalised mutational densities from early to late replicating regions in the mouse genome are shown with respect to real somatic mutations and simulated mutations. The line reflects the behaviour of simulated mutations, whereas the bars represent the behaviour for real somatic SBS-Polk mutations. **b)** Strand asymmetries for the novel SBS-Polk signature: replication strand asymmetry, genic and intergenic regions, and transcription strand asymmetry. Bar plots display the number of mutations accumulated on each strand for six substitution subtypes based on the mutated pyrimidine base C>A, C>G, C>T, T>A, T>C, and T>G. Simulated mutations on the same strands are displayed in shaded bar plots. Statistically significant strand asymmetries are shown with stars: *** P < 0.001 (Fisher’s exact test corrected for multiple testing using Benjamini-Hochberg). **c)** Pattern of the SBS-Polk mutational signature showing the percentage of mutations in transcribed and untranscribed strands in the 96-classes mutational context. **d)** Relationship between transcriptional strand bias and expression level for mutations in *Polk-/-* liver cholangiocytes clones, bulk liver and bulk kidney (*P* calculated by unpaired *t* tests). Genes are grouped into three quantiles based on expression level. The dashed line represents per-strand mutation rate in intergenic regions. **e)** Simplified diagram depicting how the activity of transcription-coupled repair (TCR) and translesion synthesis (TLS) acting on adducts of guanine would lead to the observed pattern of transcriptional-strand bias. For simplicity, only C>A (G>T) changes are shown but the same applies to C>G (G>C) and C>T (G>A) mutations.

Finally, the most prominent topographical feature of the SBS-Polk signature is its strong transcriptional-strand bias (TSB). Within genes, C>A, C>G and C>T mutations are clearly depleted from the untranscribed strand compared to the transcribed strand (**Fig. 3b, c**). The two main sources of TSB are transcription-coupled repair (TCR) or poorly understood transcription-coupled damage (TCD). The hallmark of TCD is a higher mutation rate in genic vs intergenic regions, which increases with expression level, best illustrated by SBS16 mutations in liver cancer^37,39^. As we find that SBS-Polk mutations are depleted in genic regions (**Fig. 3b**), we infer that TCR is likely the main source of TSB. Indeed, when we further subdivide genic regions into expression bins, we see depletion of mutations in highly expressed genes (**Fig. 3d**), a known consequence of more active TCR^40^. These results lead to several conclusions. First, the endogenous lesions driving SBS-Polk mutagenesis are also subject to repair by TCR, and are possibly bulky lesions that block transcription. Second, these TCR substrate lesions are most likely adducted guanines, because we observe a depletion of cytosine mutations (C>N) from the untranscribed strand (**Fig. 3e**). Third, in the absence of Polκ, the guanine adducts mispair with either A, G or T leading to SBS-Polk mutations (C>A, C>G and C>T). This is compatible with the biochemical function of Polκ, that can incorporate C opposite *N*^2^-dG adducts *in vitro*^41,42^. Together, these results reveal mechanistic details of the SBS-Polk mutations and underscore the fact that mutational signatures are complex patterns jointly shaped by the nature of the DNA lesion and the interplay of DNA repair and lesion bypass processes.

### Two DNA repair pathways protect the liver from SBS-Pol**κ** mutagenesis

Next, to more deeply understand the origin of SBS-Polk mutations, we sought to identify an *in vitro* experimental system where SBS-Polk mutations accumulate spontaneously. For this purpose, we took liver cholangiocytes (where we detected SBS-Polk mutations *in vivo,* **Fig. 1e**), cultured organoid clones for 4 months, and characterized the mutations that accrued specifically during *in vitro* culture (**Supplementary Fig. 6**). Interestingly, despite having a much higher mutation rate compared to mouse tissues, cultured *Polk-/-* cholangiocytes did not accumulate SBS-Polk mutations *in vitro*. Therefore, we conclude that the SBS-Polk mutational signature does not arise simply due to Polκ deficiency but depends on interactions with specific forms of endogenous DNA damage, likely driven by tissue-specific metabolism.

Therefore, we focused on mouse liver *in vivo* for subsequent experiments. The considerable TSB of SBS-Polk mutations suggests that the endogenous lesions driving mutation in the absence of Polκ are also repaired by a transcription-coupled mechanism. This led us to hypothesize that inactivating the relevant repair pathway would lead to an increase in mutation and potentially allow us to better characterize the endogenous DNA lesions. The best characterized TCR pathway is TC-NER, where stalling of RNA polymerase II by transcription-blocking DNA lesions triggers the excision of the damage from the transcribed strand mediated by XPA and the nucleases XPF-ERCC1 and XPG. A related pathway, not coupled to transcription, is global-genome (GG)-NER, where DNA excision of bulky adducts throughout the genome is instead triggered by topological distortion of the DNA helix, detected by XPC. To test if the TSB of SBS-Polk mutations is caused by TC-NER, we generated mice lacking GG-NER (*Xpc-/-*) or both GG-NER and TC-NER (*Xpa-/-*) in addition to loss of Polκ. Double mutants displayed no obvious phenotypes up to the age of 7 months, after which we used clonal expansion of liver cholangiocytes to characterize mutational landscapes *in vivo*.

First, we examined the mutational consequence of NER deficiency alone (**Supplementary Fig. 7a, b**). We find that *Xpa-/-* and *Xpc-/-* mouse livers accumulate mutations which we term SBS-NER, a mutational signature resembling SBS8, which has an unknown mechanism but is likely driven by endogenous NER substrates. Our results are consistent with reports from *Ercc1-/Δ* mouse livers^15^ and *XPC*-/-human leukaemia^16^, with *Xpc-/-* mouse liver carrying a mutational signature that closely resembles that observed in *XPC*-/-human leukaemia (Cosine similarity 0.93) (**Supplementary Fig. 7d**). *Xpc-/-* liver also displays a C>T component resembling SBS32 that is lacking from SBS8 and reduced in *Xpa-/-* livers. Interestingly, a side-by-side comparison reveals a significantly higher mutation burden for *Xpc-/-* compared to *Xpa-/-* mouse livers, for each class of mutation—SBS, DBS, and indels (**Supplementary Fig. 7b**). This is surprising given the canonical view that genetic inactivation of *Xpc* or *Xpa* should equally impair GG-NER. Importantly, a distinctive feature of mutations in *Xpc-/-* livers is strong TSB. Genetic disruption of *Xpc* selectively inactivates GG-NER but TC-NER remains active driving TSB in transcribed regions of the genome. As expected, we find strong TSB in *Xpc-/-* livers, which was absent in *Xpa-/-* and *Ercc1-/Δ* livers (**Supplementary Fig. 7e**). Together, these results show the mutagenic consequences of NER deficiency in the liver and its distinctive features.

Next, we examined the effect of NER deficiency on the mutational landscape of *Polk-/-* livers (**Fig. 4a**). We detected significant increases in the burden of SBS, DBS and indels in the livers of *Xpc-/-Polk-/-* and *Xpa-/-Polk-/-* mutants compared to *Polk-/-* controls (**Fig. 4b**). However, the numbers alone may just reflect the additive effect of two independent mutational processes. We used two different approaches to dissect this. First, we performed *de novo* signature extraction as above to study the consequence of NER deficiency on the SBS-Polk mutational signature (**Fig. 4c-d**). The burden of SBS-Polk mutations was slightly higher in NER-deficient *Polk-/-* mice compared to *Polk-/-* controls, suggesting that NER is a repair pathway that suppresses SBS-Polk mutagenesis, likely through excision of bulky guanine adducts (**Fig. 4e**). Conversely, we also looked at the effect of Polκ loss on the mutational signatures of NER deficiency (SBS-NER, also split into SBS8 and SBS32-like components). We observed no difference on the burden of these mutations between Polκ-proficient and deficient samples, suggesting that the NER substrates driving these signatures require mutagenic lesion bypass by DNA polymerases other than Polκ (**Fig. 4e, Supplementary Fig. 8a**).

**Figure 4.**
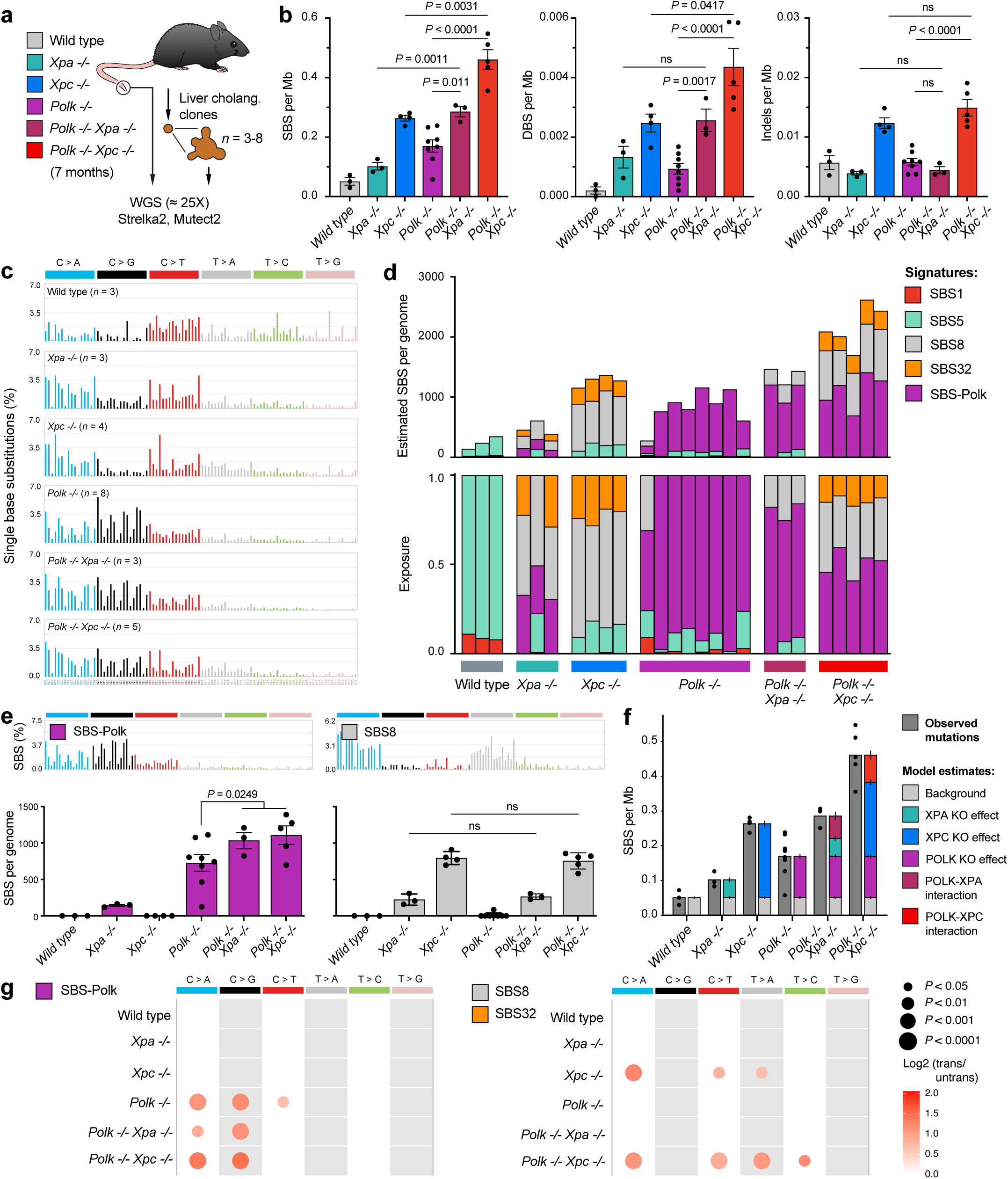
Nucleotide Excision Repair limits SBS-Polk mutagenesis. **a)** Single cholangiocytes were isolated from 7-month-old *Polk-/-Xpc-/-, Polk-/-Xpa-/-* and control mice, and expanded into clonal organoid lines, which were subjected to whole-genome sequencing and variant calling. **b)** Number of single-base substitutions (SBSs), doublet base substitutions (DBSs) and insertions/deletions (indels) per megabase (Mb) (*P* calculated by an unpaired *t* test). **c)** 96-classes of SBSs considering the six mutation types but also the bases immediately 5’ and 3’ of the mutated base. Each graph represents the average mutation pattern for *n* genomes, where *n* is indicated in each panel. **d)** Extraction and assignment of mutational signatures using SigProfiler, two main signatures are present: SBS-Polk and SBS-NER, which can be further decomposed into known COSMIC signatures SBS8, SBS32 and SBS5. Stacked bar plots showing estimated number (top) and proportion (bottom) of each mutational signature in individual cholangiocyte genomes. **e)** Quantification of SBS-Polk and SBS8 mutational signatures in *Xpc-/-Polk-/-* and control samples (*P* calculated by unpaired *t* tests). **f)** A reconstruction of the mutational burdens in each genotype using the effects of gene losses, estimated by binomial regression. The reconstructed mutational burdens are compared to the observed mutational burdens in each genotype (dark grey). The remaining bars represent the estimated effect of individual or combined gene loss on the mutational burden, with vertical lines displaying the 95% confidence interval. **g)** Quantification of transcriptional strand bias for the SBS-Polk and SBS8/SBS32 signatures. The size of the dots represents significance (*P* calculated by Fisher’s exact test corrected for multiple testing using Benjamini-Hochberg) obtained using SigProfilerTopography, while the colour represents log2 of the enrichment.

Second, we used a non-NMF approach. We hypothesized that combined loss of Polκ and NER results in an increased mutational burden beyond the additive effect of both individual gene losses, and that these mutations overlap with, but do not fully reflect the mutational spectrum of SBS-Polk. To test this, we took advantage of our genetically controlled experimental set up, directly estimating the effects of each gene deficiency on the total mutational burden, using binomial regression on the mutational burdens of wild type, *Polk-/-*, *Xpa-/-* and *Xpc-/-* livers. To determine whether a combined gene loss effect is required to explain the observed mutational burdens, we evaluated two models. The first model contains only the additive effects of the individual deficiencies, while the second model includes additional interaction effects representing the combined loss of Polκ and NER (**Fig. 4f**). The second model better fits the data, based on a likelihood ratio test (*p* value = 1.5^-49^) and a reduced Akaike information criterion (AIC) (ΔAIC = 220.7), and shows that the Polκ-NER interaction significantly contributes to the total mutational burden (*p* value < 1^-25^ for both interactions) (**Fig. 4f**, **Supplementary Table 1**). We then apply the interaction model on the 6- and 96-class SBS types and reveal patterns nearly identical to the NMF signatures (cosine similarities 0.98) and, as we hypothesized, the mutation pattern induced by the Polκ-NER interaction does not fully match the SBS-Polk signature (**Supplementary Fig. 8b, c, Supplementary Table 2**). Thus, NER loss results in a partial increase of the *Polk-/-* induced mutations rather than a uniform increase of the full SBS-Polk signature, in line with the limited increase in the SBS-Polk burden (**Fig 4e**). These findings suggest that Polκ bypasses additional lesions other than those processed by NER, or that lack of NER increases the presence of only a subset of the lesions bypassed by Polκ.

Next, we looked at transcriptional-strand asymmetry in liver genomes lacking both NER and Polκ. We find that the TSB of SBS-Polk mutations is increased in *Xpc-/-Polk-/-* and decreased in *Xpa-/-Polk-/-* mice compared to *Polk-/-* controls (**Fig. 4g**), again supporting the model where a subset of the endogenous guanine lesions driving mutation are processed by NER. Surprisingly, and in complete contrast to SBS8 mutations (**Fig. 4g, Supplementary 8d**), we find that SBS-Polk mutations retain considerable TSB in *Xpa-/-Polk-/-* samples, which completely lack NER. This effect is not due to transcription-coupled damage as the mutation rate is lower in highly expressed genes. While Xpa is considered essential for TC-NER to deal with UV damage, our data imply that residual TC-NER occurs in the absence of Xpa in response to endogenous DNA lesions. Alternatively, our findings suggest the presence of another source of TSB other than NER, potentially an alternative transcription-coupled repair pathway of endogenous guanine lesions, such as TC-BER or TC-DPC repair ^43–45^.

Finally, we analyzed DBS and indels in double mutant and control mice. While both Polκ and NER suppress DBSs, the low number limited the extraction of DBS signatures (**Fig. 4b, Supplementary Fig. 9a**). We find that indels in *Xpc-/-* samples are dominated by ID9 (which has an unknown mechanism) and not affected by loss of Polκ (**Supplementary Fig. 9b-c**). Interestingly, we detect ID6 (repair of DNA breaks) which is unique to *Ercc1-/Δ* livers, consistent with the function of Xpf-Ercc1 in crosslink repair. The lack of ID6 in other genotypes suggests that joint inactivation of Polκ and NER does not result in the formation of DNA breaks in the liver.

In summary, these data add to our mechanistic understanding of SBS-Polk mutations by showing that NER slightly suppresses SBS-Polk mutagenesis, implying that a subset of the endogenous guanine lesions are substrates of NER but that Polκ bypasses additional lesions other than those processed by NER. In addition, our results provide evidence that NER is not the sole repair pathway involved, with a second TCR route of guanine adducts limiting SBS-Polk mutations.

### Polk promotes replication-coupled bypass of endogenous dG adducts

Our results so far indicate that endogenous adducts of guanine drive tissue-specific mutation in the absence of Polκ. Uncovering the chemical nature of the DNA damage could reveal its true origins. However, we have little *a priori* knowledge of what the damage is, and analyzing DNA modifications directly is challenging. Endogenous DNA adducts are exceedingly rare (on the order of 10^−6^−10^−8^), undergo rapid repair, and have similar chromatographic properties to unmodified nucleosides. It is also likely that multiple lesions contribute to the SBS-Polk mutational signature, as our data suggests some guanine lesions are substrates of NER while others are repaired by a transcription-coupled pathway other than NER.

To tackle this fundamental question, we undertook mass spectrometric quantification of DNA adducts that might drive mutation, we first targeted known dG modifications followed by an untargeted screen aimed at identifying unknown dG lesions. DNA was extracted from mouse liver or kidney (where SBS-Polk mutations are highest), hydrolyzed, and purified as previously described^46^(**Fig. 5a**). First, we took a candidate approach. *In vitro*, Polκ bypasses DNA adducts at the *N*^2^ position of guanine in a predominantly error-free manner^18,47–49^. Therefore, we quantified the abundance of three *N*^2^-dG lesions using isotopically labelled internal standards: 1,*N*^2^-ProdG (*N*^2^-propano-dG), γ-OH-Acr-dG (*N*^2^-acrolein-dG) and εdG (*N*^2^-etheno-dG). We also included 8-oxo-dG, a small and abundant oxidative lesion, as a control. We analyzed wild type and NER-deficient tissues, to test if these lesions are excised by NER *in vivo*. We found *N*^2^-propano-dG is significantly increased in *Xpa-/-* and *Xpc-/-* kidneys, while *N*^2^-etheno-dG is higher in NER-deficient kidney and liver (**Fig. 5b**). Interestingly, *N*^2^-etheno-dG is more abundant in *Xpc-/-* than *Xpa-/-* samples, suggesting differences in the excision of this lesion *in vivo*, which correlates with differences in mutagenic burden between these genotypes (**Supplementary Fig. 7b**). Therefore, we can detect endogenous *N*^2^-dG lesions, some of which are also substrates of NER *in vivo*.

**Figure 5.**
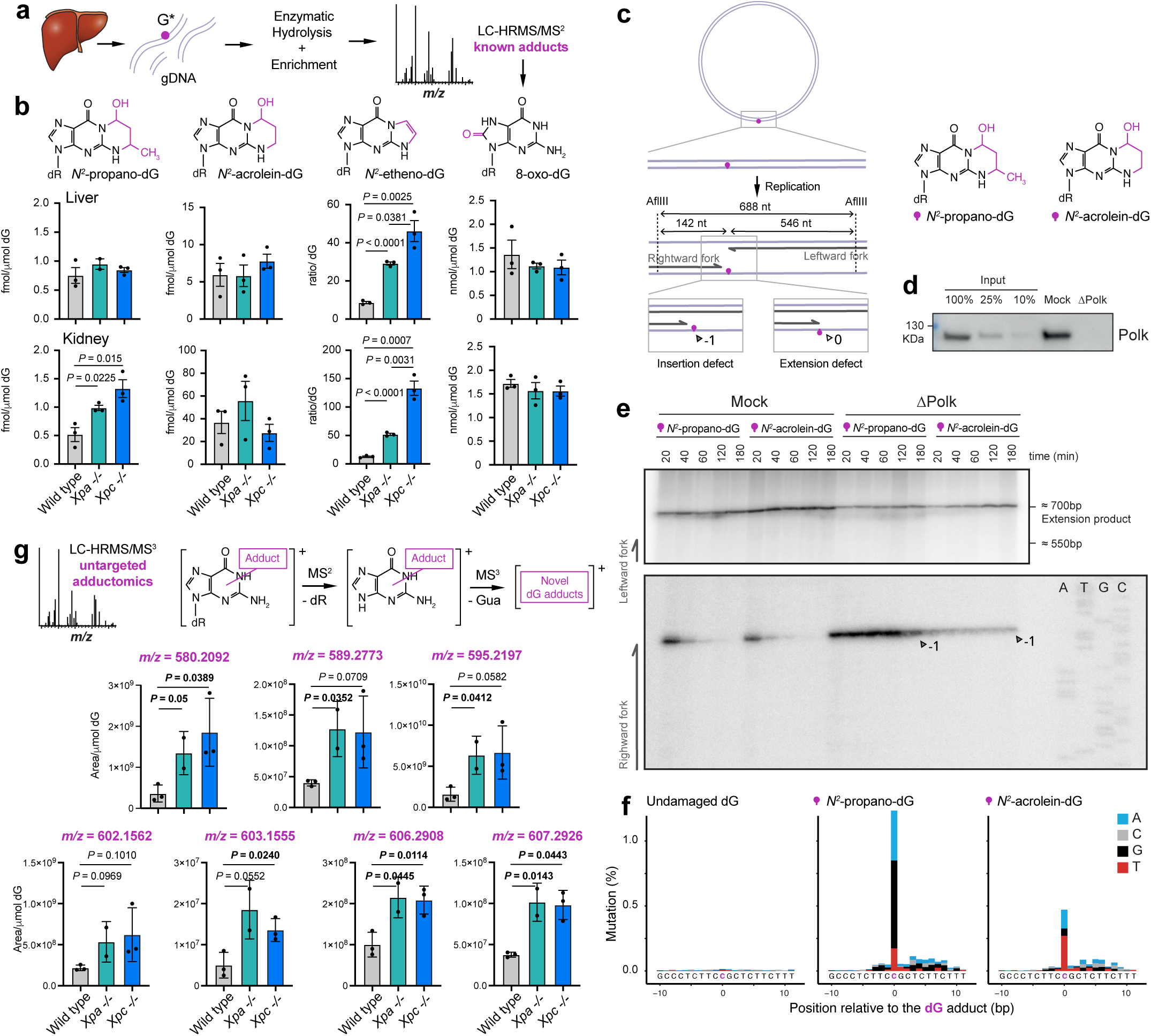
Polk is essential for the replication-coupled bypass of endogenous dG adducts. **a)** Scheme for the mass spectrometric quantification of dG adducts in mouse tissues: genomic DNA was isolated, hydrolysed, purified and analysed by mass spectrometry to quantify known dG lesions. **b)** Quantification of known dG adducts in the liver and kidney of wild type and NER-deficient mice (*P* calculated by an unpaired *t* test). Etheno-dG was measured using a relative quantitation method based on the ratio of the intensity of the signal of the analyte vs the internal standard and not on the absolute concentration. **c)** Use of *Xenopus* of egg extracts to study the replication-coupled bypass of site-specific *N*^2^-propano-dG and *N*^2^-acrolein-dG. Scheme showing the products detected in f). **d)** Western blot showing successful depletion of Polk from the extracts. **e)** Plasmids were replicated in mock or Polk-depleted (ΔPolk) extracts, repair intermediates were digested with AflIII and separated on a sequencing gel alongside a sequencing ladder. Dark grey arrows: –1 products. **f)** Distribution and frequency of nucleotide mis-incorporation in a 20 bp region flanking the lesions. **g)** Relative quantification of unknown dG adducts in the liver of wild type and NER-deficient mice (*P* calculated by an unpaired *t* test). The exact mass of the ions is shown in bold. Refer to Supplementary Figure 9 for details on workflow used to generate the list of putative adducts.

To understand the role of Polκ in bypassing these lesions, we set out to recapitulate this process. Previous studies using primer extension assays have shown that Polκ can insert a C opposite certain *N*^2^-dG lesions in a minimal system^18,47–49^. To study bypass of these adducts in a more physiological setting with *bona fide* replication forks and other TLS polymerases, we used the *Xenopus* egg extract system. We introduced site-specific *N*^2^-propano-dG and *N*^2^-acrolein-dG adducts into plasmids and replicated them in mock and Polκ-depleted *Xenopus* egg extracts (**Fig. 5c, d**). Replication intermediates were digested and separated on a sequencing gel (**Fig. 5e**). In mock depleted extracts, we find that the rightward fork (or bottom strand) that encounters the adduct shows transient stalling at 1 nucleotide (nt) before the lesion (−1 position), which is resolved after 20-40 min concomitant with accumulation of extension products. Replication products from the leftward fork, that does not encounter the lesion directly, do not show replication stalling. Depletion of Polκ causes severe stalling at the −1 position, until at least 180 minutes, indicative of a persistent insertion defect. This insertion defect in the absence of Polκ is stronger on *N*^2^-propano-dG compared to *N*^2^-acrolein-dG containing plasmids, potentially indicating that backup polymerases act more readily on acrolein adducts. Finally, we investigated the miscoding properties of these adducts after complete replication in mock extracts (**Fig. 5f**). We find that lesion bypass is largely error-free in the presence of Polκ (mutation rate 0.5-1%). The mutation was highest for bases inserted across the adducts and was followed by ≈10bp of higher mutation burden, likely reflecting collateral mutagenesis by the extension TLS polymerase^50^. Interestingly, we observed differences in the miscoding properties of the two structurally related lesions, with *N*^2^-propano-dG causing C>G and C>A mutations, and *N*^2^-acrolein-dG mostly miscoding C>T (**Fig. 5f**). In summary, we used mass spectrometry to quantify candidate endogenous *N*^2^-dG lesions *in vivo*, and show that Polκ is critical for bypass of two of these *N*^2^-dG adducts in *Xenopus* egg extracts, by accomplishing TLS insertion opposite the adducts, largely in an error-free manner.

### Characterization of novel endogenous guanine lesions by untargeted DNA adductomics

It is likely that several endogenous dG lesions contribute to the SBS-Polk signature. To further explore additional and potentially novel DNA lesions beyond the well-known adducts described above, we used our high-resolution LC/MS^3^ adductomics method for the characterization of endogenous DNA damage. The method monitors ions characterized by the neutral loss of the exact mass of deoxyribose (dR = 116.0474 ± 0.0006 *m/z*), or one of the four DNA bases (e.g., guanine+H^+^ = 152.0567 *m/z*). Importantly, this exploratory discovery experiment focused specifically on any dG adduct showing a significant increase in *Xpa-/-* and *Xpc-/-* livers, which lack damage excision and are expected to accumulate endogenous DNA lesions. Excitingly, we see seven unknown adducts which are consistently increased in NER tissues compared to wild type, with masses in the range of 580.2092 – 607.2926 *m/z* (**Fig. 5g, Supplementary Fig. 10-17**). The size and fragmentation spectra of these novel endogenous adducts are consistent with larger adducts of guanine, and possibly with some crosslinks, as seen for the adduct with *m/z* 603.1555 (**Supplementary Fig. 15**) where the loss of cytosine in the MS^2^ event triggers the appearance of guanine in the MS^3^ spectra. These findings are in line with the role of NER in repairing bulky lesions. While this suggests the exciting possibility that crosslinks are a prevalent endogenous DNA modification, we currently face the challenge of determining the precise molecular structure of these lesions and confirm their nature. This will require additional targeted investigations, including analysis with different collision energies and chromatographic conditions to collect more detailed structural information and support the synthesis of chemical standards that will allow absolute identification and quantitation. In summary, we have applied, for the first time, an untargeted adductomics approach in a setting of DNA repair deficiency, uncovering novel endogenous DNA adducts that are likely to contribute substantially to mutagenesis.

## Discussion

Here, we explore the interplay between endogenous DNA damage, DNA repair, and damage tolerance in mammalian tissues. By characterizing patterns of somatic mutation across mouse tissues, we reveal a new tissue-specific mutagenic process driven by loss of the TLS polymerase Polκ. Investigating how the novel SBS-Polk mutations are modulated by DNA repair, we obtain new mechanistic insights into how NER limits mutagenesis caused by endogenous DNA damage. Finally, we analyze the mutagenic bypass of endogenous DNA damage *in vitro* and, for the first time, exploit recent advances in the field of DNA adductomics to uncover novel sources of DNA damage.

### Mechanism of SBS-Polk mutations

We comprehensively characterize somatic mutations across mouse tissues using two complementary approaches. This allows us to identify a novel mutational signature that is suppressed by the TLS polymerase Polκ and is mainly observed in liver and kidney (**Fig. 6a**). We propose that Polκ suppresses mutation by performing error-free bypass at endogenous guanine lesions, analogous to the role of Polη in suppressing mutation by error-free bypass of UV-induced damage. Polη−deficient cells are hypermutable in response to UV, which is driven by the error-prone bypass of cyclobutene pyrimidine dimers by Polκ, Polι and Polζ. Identifying the TLS polymerases responsible for promoting endogenous SBS-Polk mutations will require the generation of additional mouse models and will be an area of future work. However, unlike Polη (*POLH*/*XPV*), no germline mutations have so far been described for human *POLK*.

**Figure 6.**
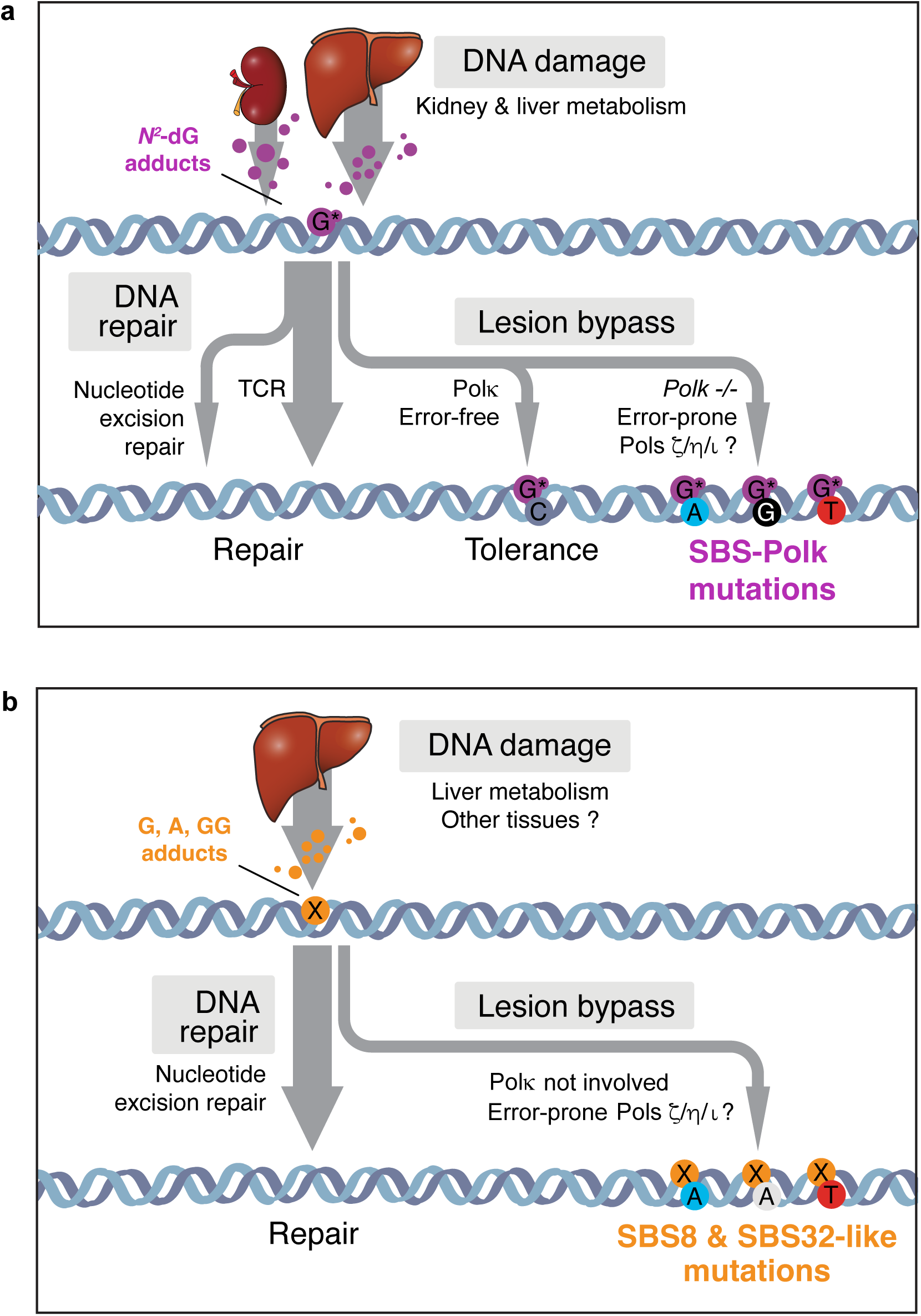
Mechanisms of mutagenesis driven by Polk and NER deficiency. **a)** Mechanism for the tissue-specific mutations that arise in the absence of Polk. Polk and at least two pathways of DNA repair cooperate to suppress C>N mutagenesis caused by endogenous guanine adducts. **b)** Building a model for spontaneous mutations caused by NER deficiency. NER suppresses SBS8 and SBS32-like mutations signatures found in cancer. These mutations are driven by adenine and guanine adducts and Polk is not involved in the genesis of these mutations.

Our finding that Polκ suppresses mutation is consistent with older work using the short untranscribed lacZ reporter sequence^51^. Here, we characterize the genome-wide pattern of mutation which allows us to study the effect of DNA repair, transcription and other topographical features on mutagenesis. From the strong transcriptional strand bias of SBS-Polk mutations, we infer that the endogenous lesions driving mutation are adducts of guanine and substrates of TCR. We test this genetically by investigating the role of NER on SBS-Polk mutations *in vivo*. NER is the best-characterized TCR pathway and indeed we find it contributes to suppression of SBS-Polk mutations, with NER-deficient *Polk-/-* mice having a modest increase in SBS-Polk mutations compared to *Polk-/-* controls. Notably, we find that SBS-Polk mutations retain considerable transcriptional strand bias in *Xpa-/-Polk-/-* mice. This is surprising, as Xpa is considered essential for TC-NER, particularly for the repair of UV-induced damage^52,53^. However, a recent report suggests some excision repair of UV-induced DNA damage is detectable in XPA-deficient humans cell lines, flies, and worms^54^. Therefore, some form of TC-NER may take place in the absence of Xpa for the repair of endogenous damage. Or, alternatively, some of the lesions underlying SBS-Polk mutations are repaired by other TCR pathways. Indeed, recent work uncovered a novel transcription-coupled pathway of formaldehyde-induced DNA-protein crosslinks (DPCs), which depends on CSB but not downstream NER factors^43–45^. Many aspects of this pathway are still unclear, namely whether the DNA-protein monoadduct is excised or bypassed but, interestingly, Polκ can perform efficient error-free bypass of dG-acrolein-peptide crosslinks^55^. In summary, our genetic dissection of SBS-Polk mutations reveals that TCR pathways other than NER may exist and suppress mutagenesis by endogenous DNA damage.

Finally, we use DNA adductomics and replication of adducted DNA in *Xenopus* egg extracts to shed light on the nature of the endogenous guanine adducts driving mutations. We identify two endogenous lesions - *N*^2^-propano-dG and *N*^2^-acrolein-dG – that are bypassed by Polκ in a largely error-free manner and are potent blocks to replication in the absence of Polκ. However, whether these are the main adducts responsible for the bulk of SBS-Polk mutations is unclear. Indeed, the untargeted adductomics approach already identified novel NER guanine substrates in the liver which could potentially be mutagenic, so it is plausible that the SBS-Polk signature is the cumulative effect of several guanine adducts. Further insights into the main source of damage could be gained from genetic experiments; for example, modulation of candidate sources of damage (e.g. formaldehyde-induced DPCs) or repair (e.g. TCR) could help refine our understanding. Alternatively, DNA adductomics applied to a panel of tissues could identify guanine lesions which are specifically increased in the kidney and liver, correlating with the burden of SBS-Polk mutations in these tissues.

### Mechanistic insights into spontaneous mutagenesis driven by NER deficiency

We explore the role of NER in shaping the landscape of the novel SBS-Polk mutations and also uncover interesting mechanistic insights into cancer mutational signatures linked to NER deficiency alone (**Fig. 6b**). Mouse livers lacking *Ercc1* or human cancers lacking *XPC* accumulate a mutational signature that resembles SBS8, with an additional C>T component that is similar to SBS32^15,16^. SBS8 is found in lung, brain, breast, and prostate cancers and has an unknown etiology. SBS32 is caused by azathioprine treatment^56^, but an endogenous SBS32-like mutational signature of unknown origin is found in astrocytes and blood stem cells^22,57^.

We address gaps in our knowledge of these signatures by contrasting the mutational signatures and burden of mouse liver lacking *Xpa* and *Xpc* in an isogenic system via side-by-side comparison. As expected, mutations in *Xpc-/-* livers have strong TSB, which is absent from either *Xpa-/-* or *Ercc1-/Δ* livers. But surprisingly, we find a significantly higher mutation burden of SBS, DBS, and indels in *Xpc-/-* livers compared to *Xpa-/-* livers. These results challenge the canonical view that *XPC* and *XPA* mutations should equally inactivate GG-NER; if the current model were correct, mutagenesis should be equivalent, if not higher, in *Xpa-/-* when compared to *Xpc-/-* livers. A similar observation was made recently with the sequencing of skin cancers from Xeroderma pigmentosum patients, when it was found that *XPC* genomes have a higher burden of UV-induced mutagenesis compared to *XPA* or *XPD*-deficient cancers^8^. The difference was attributed to differences in disease severity and UV exposure, but this cannot explain our results in endogenous liver mutagenesis. When controlling for age, the mutation burden in *Xpc-/-* mice was comparable to *Ercc1-/Δ* mice. Therefore, background residual NER excision is a potential explanation for lower mutation burden in *Xpa-/-* mice. In line with this, we find more *N*^2^-etheno-dG adducts in *Xpc-/-* livers compared to *Xpa-/-* livers. An alternative explanation is a unique and non-canonical function of Xpc in limiting mutagenesis. Expanding our analysis to other NER-deficient mice (e.g. *Xpe*, *Xpd*) will help distinguish between these possibilities. Much of our knowledge of NER comes from exposing cells to UV radiation, while our results highlight key differences in the repair of endogenous DNA damage.

We show that loss of Polκ had no effect on the burden of SBS8, SBS32, and ID9-like mutations, indicating that Polκ is not involved in the generation of mutations from NER deficiency. Interestingly, *Xpc-/-Rev1-/-* mice succumb to bone marrow failure demonstrating a strong interaction between NER and TLS^58^. The consequences for mutagenesis have not been explored, so it will be interesting to characterize mutagenesis in this and other NER/TLS double mutants to identify TLS factors that mediate the bypass of endogenous NER substrates. Finally, with regard to the endogenous damage driving mutation, we infer from the TSB in *Xpc-/-* livers that endogenous lesions are adducts of guanine and adenine; the increased DBS also point to tandem lesions or intrastrand crosslinks. We have shown that formaldehyde is one endogenous factor that drives phenotypes associated with loss of NER^59^. The untargeted adductomics analysis presented here, even though unable to currently provide the precise structural identity of the adducts detected, will pave the way to identifying novel sources of damage.

### Unbiased DNA adductomics to uncover endogenous DNA damage

The identification of DNA adducts is important for understanding mechanisms of mutagenesis and cancer initiation. Unlike targeted methods that focus on known adducts, untargeted DNA adductomics seeks to detect all possible DNA adducts in a sample without prior knowledge of what those adducts might be. Until recently, this powerful approach was limited by technology, but recent advances in mass spectrometry led to the development of DNA adductomics that uses high-resolution data-dependent scanning and neutral loss MS^3^ triggering to profile all DNA modifications. This approach has allowed us to identify novel DNA adducts, including cross-links in several experimental settings including tobacco-specific nitrosamine NNK^60^, the gut bacterial-derived genotoxin Colibactin^61^ and the chemotherapy agents Busulfan and Cyclophosphamide^62,63^. For the first time, we applied this methodology to characterize adducts that accumulate spontaneously in a setting of DNA repair deficiency. We took the first steps in this direction, focusing on adducts of guanine repaired by NER, and found seven novel large adducts which were consistently increased in NER-deficient livers. Further work is needed to identify the structure of these lesions, confirm the identity of any crosslink, to determine which of them are functionally relevant and mutagenic, and to identify their possible sources. However, our proof-of-concept experiment gives us confidence that the combination of genetics with untargeted DNA adductomics will become a powerful tool to dramatically expand our ability to explore the complex interactions between metabolism and DNA damage, and their roles in disease.

Taken together, our work explores the complex interplay between endogenous DNA damage, DNA repair, and damage tolerance. Our work uncovered a new mutagenic process, and provides key mechanistic insights into cancer-associated mutational signatures.

## Acknowledgements

The authors would like to thank members of the Hubrecht Institute Flow Cytometry and Animal facilities for essential support. We thank Gerry Crossan, Francesca Mattiroli, Ina Sonnen, Jacques Bothma and members of the Garaycoechea lab for critical reading of the manuscript. The research was supported by Dutch Cancer Society KWF Young Investigator Grant (project 12260) and Dutch Research Council NWO VIDI Grant (project VI.Vidi.213.046).

## Author contributions

Organoid isolation and culture, Y.J., J.W., J.I.G.; bioinformatic analysis Y.J., L.B., J.v.S.; Nanoseq sequencing and data analysis, M.P., A.B.O., F.A., I.M.; mouse husbandry and experimentation J.W., DNA adductomics, F.C.J., D.M., P.W.V., S.B.; *Xenopus* egg extracts R.v.d.S, A.E.E.V., J.B., P.K.; figure preparation, Y.J., L.B., J.I.G.; study concept and design, Y.J., J.I.G.; manuscript Y.J., J.I.G. with contributions from all authors.

## Methods

### Mice

All animal experiments were performed after institutional review by the Animal Ethics Committee of the Royal Netherlands Academy of Arts and Sciences (KNAW) with project license of AVD8010020198847. The *Polk^tm^*^1^*^.1Rsky^* (MGI 2445458, C57BL/6J) mice were described previously and acquired from JAX^64^. *Xpa^tm1Hvs^*(MGI 1857939, C57BL/6) and *Xpc^tm1Ecf^* (MGI 1859840, C57BL/6) mice were described previously and a kind gift from G.T. van der Horst, Errol Friedberg and Jan Hoeijmakers^65,66^. We used 18-month old mice for organoid isolation and NanoSeq sequencing of wild type and *Polk-/-* mice, and 7-month-old mice for cholangiocyte organoid isolation of *Xpa-/-Polk-/-, Xpc-/-Polk-/-* and age-matched controls.

### Organoid culture

Liver cholangiocytes were harvested and enzymatically digested as previously reported^67^. Briefly, minced liver was digested with 125 µg/mL collagenase (Sigma-Aldrich), 125 µg/mL dispase II (ThermoFisher) and 0.1 mg/mL of DNAase (Sigma-Aldrich) in wash buffer. The wash buffer was based on DMEM (ThermoFisher) supplemented with penicillin/streptomycin (ThermoFisher) and 2% fetal bovine serum (FBS) (Sigma-Aldrich) as described by Broutier *et al*.^68^. Biliary tree fragments and associated stroma were dissociated into single cells using 7x TrypLE (Gibco) and subjected to fluorescence-activated cell sorting (FACS) based on size and granularity and singlets using a BD Influx™ Cell Sorter. Cholangiocytes were selected based on EpCAM positivity and negative exclusion of the hematopoietic/endothelial markers CD31, CD45 and TER-119. The isolated cholangiocytes were subjected to clonal expansion as previously reported^69^. Cells were cultured in basal medium (advanced DMEM/F12 with 10 mM HEPES, 1x Glutamax, and Penicillin/Streptomycin of 100 U/mL) supplemented with B27 (Invitrogen), 1 μM N-acetylcysteine (Sigma-Aldrich), 10 nM gastrin (Sigma-Aldrich), 50 ng/mL mEGF (PeproTech), 50 ng/mL rhHGF (Bio-Techne R&D), 100 ng/mL FGF-10 (Peprotech), 1 % R-spondin 3 conditioned medium, 10 mM nicotinamide (Sigma-Aldrich) and 10 µM Rho-kinase inhibitors (AbMole). Additionally, the cells were supplemented with 150 ng/mL Noggin (IPA) and 27ng/mL Wnt surrogate (IPA) for the initial four days following seeding.

The preparation of mouse lung cell suspensions was conducted via collagenase digestion of the lungs, with the upper airways removed as previously described^70^. The dissociated cells were resuspended and incubated in 2% FCS with antibodies, including CD31, CD45, EpCAM, MHCII and CD24. The labeled cells were then washed and sorted using a BD Influx™ Cell Sorter. Club cells were defined as CD31/CD45-EpCAM+ MHCII-CD24low. The sorted cells were then subjected to single-cell expansion in Matrigel (BD Biosciences) and subsequently cultured in basal medium supplemented with 50 ng/mL EGF, 100 ng/mL FGF-7, 100 ng/mL FGF-10, and 2 µM Rho-kinase inhibitors.

Gastric epithelial stem cells were harvested, and cultured as reported^71^. In brief, the glands were extracted from 1 cm^2^ of mouse stomach using EDTA in cold PBS. The gastric glands were filtered through a 100 mm strainer and cultured in Matrigel, followed by limiting dilution for single-cell expansion. The culture medium for gastric epithelial stem cells is based on basal medium supplemented with 150 ng/mL noggin, 27ng/mL Wnt surrogate, 2% R-spondin3 conditioned medium, 50 ng/mL EGF, 100 ng/mL FGF-10, 10 nM gastrin, 0.5 mM TGF-β inhibitor and 10 µM Rho-kinase inhibitor.

Small intestinal crypts were isolated according to the methodology previously described^72^. Briefly, 1 cm small intestines were incised lengthwise, chopped into pieces, and thoroughly washed with cold PBS. The tissue fragments were vigorously shaken in PBS containing 2.5 mM EDTA and incubated on ice for 30 minutes. This process was repeated once more, after which the supernatant was passed through a 70 µm cell strainer in order to remove residual villous material. Subsequently, the isolated crypts were subjected to centrifugation and re-suspension for bulk culture, followed by limiting dilution for single-cell expansion. The cells were cultured in basal medium supplemented with basal medium supplemented with B27, 1 mM N-acetylcysteine, 10 nM gastrin, 50 ng/mL EGF, 1% R-spondin3 conditioned medium, 10 mM nicotinamide, 10 µM Rho-kinase inhibitors, and 27ng/mL Wnt surrogate. Bone marrow cells were harvested and cultured as reported previously^73^. Briefly, the bone marrow cells were collected from both the femur and tibia of the mouse using wash media (DMEM + 2% FCS), and filtered through a 40 µm strainer. The total bone marrow was suspended and cultured in MethoCult GF M3434 (Stem Cell Technologies) semi-solid medium, followed by limiting dilution for single cell expansion.

### Whole genome sequencing of cultured organoids and data processing

Genomic DNA was extracted from the respective organoid samples using the QIAamp DNA Mini Kit (Qiagen). The library preparation and sequencing were conducted by Novogene. The sequencing was performed using the Illumina 2×150 bp paired-end sequencing on a Novaseq6000 platform, with a minimum coverage of 20X. The quality of sequencing reads was evaluated using FastQC (v0.11.9), and the adapters sequences, primers, poly-A tails, and other undesirable sequences were removed using cutadapt (v4.2). The sequencing reads were mapped to the mouse genome GRCm38 (mm10) using BWA-MEM with the default settings. Further mapping cleanup was conducted with Samtools (v.1.6), Picard CleanSam, Picard FixMateInformation and Picard MarkDuplicates (v.2.18.29).

### Variant calling and filtering

Somatic single and doublet base substitutions were identified using Strelka (version 2.9.10), with the corresponding tail sample serving as the normal. The quality of single nucleotide variant (SNV) calls was evaluated using FINGs (version 1.7.2) with the default settings, with the exception of a maximum depth threshold of ≤60 and a maximum variant allele frequency (maxvafnormal) of 0.01 in the normal sample (tail). A combination of Strelka and GATK Mutect2 (version 4.5.0) was employed to identify high-quality indels. Only the indels identified by both tools were subjected to further evaluation, requiring a mapping quality (MQ) > 50, a read depth >10 and <60 in both tumor (organoid) and normal (tail) samples. To minimize sequencing strand bias, valid indels were required to be present on at least two forward and two reverse read strands. Small indels (<4 bp) located within tandem repeat regions of at least 9 repeats were excluded. All SNV and indel variants with an allele frequency <0.3 or >0.7 were excluded to ensure clonality and minimize sequencing artifacts. For both SNVs and indels, only positions flagged as “PASS” by Mutect2 and Strelka were considered. Furthermore, variants that were present in samples from the same parental clone, as well as in normal (tail) samples or other mice, were discarded to eliminate potential germline variants.

### NanoSeq library preparation and sequencing

Restriction-enzyme NanoSeq libraries were prepared from 20 ng genomic DNA of a respective sample as input. In the case of matched normal samples, 40ng of genomic DNA extracted from the same mouse tails was used to prepare an undiluted NanoSeq library. The preparation of all libraries was in equivalence with the protocol described in the original publication of the NanoSeq method^19^. The NanoSeq libraries were sequenced using 2×150 bp paired-end reads on a NovaSeq 6000 platform at the Wellcome Sanger Institute. The data pre-processing was implemented in alignment with descriptions by Abascal and colleagues ^19^. Briefly, reads were aligned to the mouse genome GRCm38 using BWA-MEM. Alignments were then sorted using biobambam2 as previously reported^74^.

### NanoSeq variant calling

Variant calling was performed as described previously ^19^. Briefly, the method requires a matched normal sample from the same individual to filter out germline SNPs. For this purpose, matched normal samples were generated from undiluted NanoSeq libraries of the tail of each mouse. For a mutation to be called as a variant, several criteria had to be fulfilled: (1) each read bundle had to contain a minimum of two reads from each of the two original DNA strands; (2) the consensus base quality scores needed to be ≥ 60; (3) the minimum difference between the primary (AS) and secondary alignment score (XS) was > 50 to keep only read pairs with unambiguous mapping; (4) the average number of mismatches in a group of reads should not be > 2, either in the matched normal or sample itself; (5) the maximum number of 5’ clips needed to be 0; (6) the minimum number of improper read pairs needed to be 0; (7) base calls in read ends, referring to the last 8 bp from the 5’ or 3’ ends, are discarded; (8) for SNV calling, reads in the RB are not allowed to contain indels; (9) the number of reads per strand in the matched at a given site was required to be ≥ 15; (10) for a given mutation, the respective base will not be seen at a frequency > 0.01 in the matched normal; (11) a site should not overlap a common SNP and noise mask. We created this mask by calling variants across all matched normal samples, including variants supported by ≥ 2 reads and VAF ≥ 0.01. The resulting mask had a size of 79MB in total.

### Mutation burden and trinucleotide substitution profiles

Given the biases in the creation of restriction-enzyme NanoSeq libraries, resulting from trinucleotides overlapping the restriction enzyme site, a correction for sequence composition is applied to each of the 96 possible trinucleotide substitutions as detailed in the methods for the original publication^19^. We use corrected substitution counts to calculate the corrected mutation burden, as well as the extrapolated burden per genome/cell by multiplying the burdens with the size of a diploid mouse genome (2 x 2.6 Gb). Indel burdens were calculated by dividing the number of detected indels by the total number of base pairs sequenced. Confidence intervals were determined by performing an exact Poisson test in R (poisson.test).

### Mutational Signature Analysis

Mutations were analyzed using SigProfilerMatrixGenerator^75^ (v. 1.2.25) to classify mutations into specific categories and generate mutational spectra plots. The resulting matrix was then utilized as the input for signature extraction. Two algorithms were employed for this purpose: SigProfilerExtractor^76^ (v. 1.1.23), which is based on non-negative matrix factorization, and mSigHdp^25^ (v. 2.1.2), which utilizes a Bayesian hierarchical Dirichlet process mixture model. In addition to the standard SBS96 catalogue, we also extracted de novo signatures in the context of SBS288, which divides the genome into transcriptional, non-transcriptional, and intergenic regions. SBS288 considers the influence of DNA repair on mutation in gene bodies and provides further insight into the disparate mutagenic pathways that assist in differentiating de novo signatures from those that have already been identified. Where applicable, the extracted de novo signatures were decomposed using SigProfilerAssignment^77^ (v. 0.1.3) to the COSMIC signature database (v. 3.4), with potential artifact signatures excluded. The candidate signatures considered for decomposition were restricted to those suggested by SigProfilerExtraction. For SBS288 signatures, the aforementioned decomposition was performed subsequent to the collapsing of the 288 channels into the standard 96 catalogue. A comparison of the results obtained from the two methods, as well as the decomposition of de novo signatures into COSMIC signatures, is presented in Figure S3. The specific extraction parameters for both methods are available upon request.

### Topography of mutational signatures

The analysis of mutational signatures was conducted using SigProfilerTopography (v. 1.0.85) to examine strand asymmetries and distributions of mutations related to replication time. The detailed workflow for SigProfilerTopography has been previously described^31^. In summary, the workflow randomly generated SBS while preserving the original somatic mutation patterns in each sample at a predetermined resolution. Mutations were retrieved from each strand/region across six mutational channels (C>A, C>G, C>T, T>A, T>C, and T>G), from real and simulated results. P values were calculated for the odds ratio between the ratio of real mutations and the ratio of simulated mutations. Only those strand asymmetries with a corrected p-value <0.05 and odds ratios >1.10 were considered to be significant.

### Transcription Strand Asymmetries Analysis

A comparative analysis of tissue-specific transcription strand bias between genes expressed at low and high levels was conducted using R (v. 4.3.3) and RStudio. The clonal data was derived from cholangiocyte-specific RNA-seq from Aloia *et al*.^78^, and the mice liver-specific RNA-seq from Li *et al*.^79^. The mutations were classified as either transcribed or untranscribed within gene bodies, and the genes were divided into three quantiles based on their expression level. The statistical significance of each quantile was determined using unpaired t-tests. In the case of clonal samples, only genomic regions with a coverage greater than 20x were included in the analysis to calculate the mutational burden. Due to the restriction-enzyme used in the NanoSeq experiment, approximately 30% of the genome was sequenced; consequently, the analysis was confined to these sequenced genic regions.

### Binomial regression of mutational burdens

The effects of gene losses were estimated using two binomial general linear models, with the total number of mutated bases per covered base in mouse liver cholangiocytes as the response variable. As we expect independent mutational effects to act additively, an identity link was used for the binomial models. Each individual gene loss, as well as combined gene loss interaction effects are modeled as binary variables, where the intercept represents the background mutational burden. The simple additive model includes only the individual gene losses, while the full model includes both individual gene loss variables as well as interaction effects for combined gene losses. The binomial general linear model was fitted using the ‘GLM’ function from the python *statsmodels* package^80^. To evaluate the fit of the models to the data, the difference in Akaike information criterion (AIC) was determined and a likelihood ratio test was performed. To investigate the mutational profile of the mutations resulting from each gene deficiency, regression with the full model was repeated on each of the six substitution types as well as on the 96 trinucleotide mutation classes, using the total burden of each mutation type as the response variable. The reported p-values and confidence intervals for the estimated effects have been Bonferroni-corrected to account for multiple testing.

### Replication-coupled lesion bypass in *Xenopus* egg extract

All *Xenopus laevis* procedures were performed in accordance with national animal welfare laws, reviewed by the Animal Ethics Committee of the Royal Netherlands Academy of Arts and Sciences (KNAW) (license number AVD80100202216633). Preparation of *Xenopus* egg extracts and DNA replication were performed as previously described^81,82^. For DNA replication, plasmids were incubated in a high-speed supernatant extract (HSS) at a final concentration of 7.5 ng/µl for 20 min at room temperature to license the DNA. Two volumes of nucleoplasmic extract (NPE) were added to start DNA replication. To label the nascent strands, HSS was supplemented with ^32^P-α-dCTPs. At the indicated time, the reactions were stopped with 10 volumes of replication stop solution II (Stop II: 50 mM Tris pH 7.5, 0.5% SDS, 10 mM EDTA pH 8)., followed by Proteinase K (0.5 μg/μl) treatment for 1 h at 37 °C or overnight at room temperature. DNA was phenol/chloroform extracted; ethanol precipitated with glycogen (0.3 μg/μl), and resuspended in 10 mM Tris pH 7.5 in a volume equal to the reaction sample taken.

Polk was depleted from *Xenopus* egg extract using an antibody raised against a C-terminal peptide (KSKPNSSKNTIDRFFK) of *Xenopus laevis* Polk (Biosynth). Affinity-purified Polk antibody was incubated with Dynabeads Protein A beads (Thermo Fisher Scientific) to their maximum binding capacity. Two volumes of the antibody-coated beads were then mixed with one volume of HSS or NPE and incubated for 30 min at 4 °C. Depleted extracts were collected and immediately used for replication assays. Nascent strand analysis was performed as previously described^83^. In brief, DNA replication products were digested with AflIII, one volume of denaturing PAGE Gel Loading Buffer II (Invitrogen™) was added, the samples were separated on a 7% polyacrylamide sequencing gel and visualized by autoradiography. The sequencing gel ladder was produced using the Thermo Sequenase Cycle Sequencing Kit (USB) and primer S (5’-CATGTTTTACTAGCCAGATTTTTCCTCCTC-3’).

### Generation of 1,*N*^2^-ProdG (*N*^2^-propano-dG), γ-OH-Acr-dG (*N*^2^-acrolein-dG) containing plasmids

Generation of the plasmid containing a site-specific 1,*N*^2^-ProdG (*N*^2^-propano-dG) was described previously^84^. The plasmid containing a site-specific γ-OH-Acr-dG (*N*^2^-acrolein-dG) was generated using a similar method. Specifically, a custom oligonucleotide (5’-[phos]-GCA CGA AAG AAG AGC 2FdI-GA AG-3’, Eurogentec) was synthesized, using a 5’-Dimethoxytrityl-2-fluoro-O6-p-nitrophenylethyl-2’-deoxyInosine,3’-[(2-cyanoethyl)-(N,N-diisopropyl)]-phosphoramidite, and shipped on its support. The support (ca 0.25 μmol oligo) was incubated overnight with 7 mg 4-amino-1,2-butanediol (FluoroChem) in 220 μl DMSO and 110 μl TEA with agitation at RT. The support was washed three times with 200 μl DMSO and 400 μl CH3CN, followed by deprotection of the O6-p-nitrophenylethyl group with 300 μl of 1 M DBU in CH_3_CN at RT for 1 h. The support was washed three times with 250 μl CH_3_CN and treated with 500 μl aq. 28% NH_4_OH at 55 °°C for 6 h to remove the remaining protecting groups and elute the N^2^-(3,4-dihydroxybutyl)-guanine-modified oligo from the support. The oligo was dried using a SpeedVac, resuspended in water, applied to a MonoQ 5/50 GL column in buffer A (10 mM TRIS-HCL pH 7.5, 100 mM NaCl) and eluted in a gradient of 3% buffer B (10 mM TRIS-HCL pH 7.5, 800 mM NaCl) per CV at 4 °°C. The N^2^-(3,4-dihydroxybutyl)-guanine-modified oligo eluted at 46.1 Ms/cm. Peak fractions were pooled and re-injected to increase purity followed by desalting using a NAP-5 column (Cytiva). The modified oligo was reacted with 50 mM NaIO4 for 1 h at RT and the reaction was quenched by desalting over a NAP-5 column in MQ water. The resulting γ-hydroxy-1,N^2^-propanoguanine-modified oligodeoxynucleotide was mixed at a 1:1 molar ratio with the complementary oligo: 5’-[phos]-CCC TCT TCC GCT CTT CTT TC-3’ in PBS and annealed at 85 °C for 5 min, ramped to 25°C at - 0.1°C s-1 and flash frozen immediately to prevent crosslink formation.

### Mutational analysis of replicated *N*^2^-propano-dG and *N*^2^-acrolein-dG containing plasmids

To analyze mutations generated upon replication of lesion containing plasmids in *Xenopus* egg extract, 5 ng of RNaseA treated extracted DNA from a replication reaction was used. Using an equimolar amount of primer A (5’-GTT CAG ACG TGT GCT CTT CCG ATC TNN NNN NNN NNN NNN NNT AGG TGT TGG GGC GGG ACT ATG GTT GCT GAC T-3’), that anneals to the lesion-containing strand (123 nt downstream of the lesion) and contains a sample specific barcode and 16 nt UMI sequence, a linear amplification was performed with Herculase II Fusion polymerase (1 unit, Agilent Technologies). Reaction products were further amplified using a nested PCR with Primer B (5’-GA CTG GAG TTC AGA CGT GTG CTC TTC CGA TCT-3’), and primer C (5’-ACA CTC TTT CCC TAC ACG ACG CTC TTC CGA TCT CTC CTG ACT ACT CCC AGT CAT AGC TGT CCC-3’), annealing 114 nt upstream of the lesion. Illumina-compatible adapters were incorporated by subsequent PCR amplification using NEBNext Dual Index Primers and the libraries were sequenced by Novogene. The sequencing reads were deduplicated using umi tools (v1.1.6), mapped to the reference plasmid using BWA-MEM, and further cleaned using fgbio (v2.4.0) to remove the soft-clip sequences. The mapped paired-end reads were converted to fastq files using bedtools(v.2.31.1), which was subsequently used for analysis using SIQ (v1.8) with default settings except a max base error of 0.001.

### DNA Extraction for Mass Spectrometry

Genomic DNA from liver and kidney tissues of wild-type, *Xpc -/-*, and *Xpa -/-* 10-month-old mice was extracted using Puregene Kit (Qiagen) with modifications. Briefly, tissues were minced and lysed using the Cell Lysis Solution supplemented with 1 mM glutathione, 200 mM pentostatin, 100 mM deferoxamine, 100 mM butylated hydroxytoluene, and 25 mL proteinase K. Tissue samples were lysed overnight on a rotator at 25°C, followed by RNase treatment and protein precipitation. DNA was then precipitated with isopropanol, washed, and dissolved in TE buffer containing antioxidants. The DNA solution was further purified using chloroform/isoamyl alcohol extraction to ensure removal of contaminants, followed by a final DNA precipitation, washing, and drying. Throughout the process, care is taken to degas the buffers and minimize oxidative stress on the samples, ensuring the integrity of the isolated DNA.

### DNA Enzymatic Digestion, sample purification and enrichment

The hydrolysis and purification of the isolated DNA was performed similarly to what has been described previously^85^. Isolated DNA (60 µg) from the livers and kidneys of mice was dissolved in 800 µL buffer of 10 mM sodium succinate, 5 mM CaCl_2_, and 5mM GSH (pH 7.0). A buffer blank (800 µL of buffer) and calf thymus DNA (60 µg in 800 µL of buffer) were prepared as negative and positive controls, respectively. The DNA was then enzymatically hydrolyzed with 30 units of micrococcal nuclease and 0.18 units of phosphodiesterase II incubated at 37 °C for 5 hours. Then, 60 units of alkaline phosphatase (from calf intestine) was added, and the mixture was incubated at 37 °C overnight. The following day [^13^C_5_]1,*N*^2^-ε-dG (*N*^2^-etheno-dG), [^13^C ^15^N_5_]ɣ-OH-Acr-dG (*N*^2^-acrolein-dG), [^15^N_5_](6S,8S;6R,8R)ɣ-OH-Cro-dG (*N*^2^-propano-dG), and [^13^C^15^N_2_]-8-oxo-dG were spiked into the hydrolysate as internal standards. The samples were then added to Amicon 10K filters (Ultracel^®^ 10K, Millipore) with centrifugal filtration performed at 14000 x *g* for 20 min. After filtration, 10 μL aliquots were removed for dG quantitation by HPLC. The rest of the hydrolysate was purified using solid-phase extraction (SPE) with reversed-phase separation (Strata-X, 33 µm, 30 mg/1 mL (Phenomenex)). The SPE cartridges were activated with 3 mL of CH_3_OH and 3 mL of H_2_O/0.1 mM GSH. After the samples were added, the cartridges were washed with 6 mL of H_2_O/0.1 mM GSH, 1 mL of 3% CH_3_OH in H_2_O/0.1 mM GSH. The analytes were eluted with 1 mL 70% CH_3_OH in H_2_O/0.1 mM GSH into 1.2 mL silanized glass vials (Chrom Tech) containing 0.65 µL of 100 mM GSH. The eluted samples were then evaporated to dryness via SpeedVac and stored at −20 °C until analysis.

### Quantitation of dG

Quantitation of dG was conducted using a Dionex UltiMate 3000 RSLCnano HPLC system (ThermoFisher) with a UV detector set to 254 nm and equipped with a Luna C18 column (25 cm x 0.5 μm ID, 5 μm, 100 Å) (ThermoFisher). The mobile phases were H_2_O (A) and CH_3_OH (B), the flow rate was 15 µL/min, and 2 µL were injected into the system. The gradient started at 5% B for 1 min, followed by an increase to 25% in 10 min. The gradient then increased to 95% in 3 min and was maintained at those conditions for 5 min before returning to 5% B in 2 min. The instrument was re-equilibrated at 5% B for 9 min for a total run time of 30 min. A calibration curve for dG (ranging from 2 ng/µL to 32 ng/µL in H_2_O) was run in duplicate and used to calculate the dG content in each sample.

### Quantitative Parallel Reaction Monitoring (PRM) of Endogenous Adducts

Samples from wild type, *Xpa -/-* and *Xpc -/-* liver and kidney DNA were reconstituted in 20 μL of H_2_O for LC-MS^2^ analysis targeting known endogenous DNA adducts. The analysis was done using an Orbitrap Exploris 480 instrument (ThermoFisher) coupled to a Vanquish™ Neo UHPLC system, (ThermoFisher) using positive nanoelectrospray ionization (NSI) with a source temperature of 300°C and a spray voltage of 1900V. The reversed phase chromatographic separation was performed using a nanoflow column (50 cm x 75 μm, CoAnn Technologies, Richland, WA) self-packed with Luna C18 (5 μm,100 Å, Phenomenex) stationary phase. The mobile phases consisted of 5 mM NH_4_OAc (A) and 95% CH_3_CN in H_2_O (B) and the injection volume was 4 μL. The gradient started with an increase from 1% to 5% B over 5 min at a flow rate of 0.3 μL/min, followed by an increase to 22% B over 30 min. The gradient was then increased to 95% B over 1 min and the flow rate was increased to 0.9 μL/min. Finally, the gradient was maintained at 95% B for 2 min and the flow rate was increased to 1.0 µL/min to wash the system for a total run time of 43 min. The column was re-equilibrated at the starting conditions with 5 column volumes to prepare for the next injection. This targeted method included the following precursor ions and corresponding extracted product ions used for quantitation: 338.1459 *m/z* → 222.0986 *m/z* for *N*^2^-propano-dG; 343.1311 *m/z* → 227.0837 *m/z* for [^15^N_5_]*N*^2^-propano-dG; 324.1302 *m/z* → 208.0829 *m/z* for *N*^2^-acrolein-dG; 339.1490 *m/z* → 218.0848 *m/z* for [^13^C ^15^N]*N*^2^-acrolein-dG; 284.0989 *m/z* → 168.0516 *m/z* for 8-oxo-dG; 287.0964 *m/z* → 171.0490 *m/z* for [^13^C^15^N_2_]-8-oxo-dG; 292.1040 *m/z* → 176.0567 *m/z* for *N*^2^-etheno-dG and 297.1208 *m/z* → 176.0567 *m/z* for [^13^C_5_]*N*^2^-etheno-dG. MS^2^ fragmentation was performed with a quadrupole isolation width of 1.5 m/z, HCD collision energy of 30%, AGC value of 1000%, maximum injection time of 200 ms, and a resolution setting of 60,000. A 100-650 *m/z* full scan event with a resolution setting of 15,000 was included to monitor for any anomalies in sample composition or irregularities in the analysis. Calibration curves were prepared using standard solutions of the *N*^2^-acrolein-dG and *N*^2^-propano-dG ranging from 2.5 to 250 amol/μL with 100 amol/μL of the internal standards [^13^C ^15^N]*N*^2^-acrolein-dG and [^15^N]*N*^2^-propano-dG. In a separate calibration curve, a constant amount of the internal standard [^13^C^15^N_2_]-8-oxo-dG (1 fmol/μL) was mixed with different amounts of 8-oxo-dG (10–400 fmol/μL). Utilizing these calibration curves, we were able to absolutely quantify each of our adducts except for *N*^2^-etheno-dG which was not included in the calibration curve standard mix. Semi-quantitation of *N*^2^-etheno-dG was performed by assuming linear and equal response for *N*^2^-etheno-dG and [^13^C_5_]*N*^2^-etheno-dG in the sample data. Quantified adduct levels were all normalized to the measured dG amounts.

### Parallel Reaction Monitoring of Putative Adducts

Samples from wild-type, *Xpa -/-* and *Xpc -/-* liver DNA were prepared for LC-MS^2^ analysis of putative DNA adducts detected in the screening assay using an Orbitrap Lumos instrument (ThermoFisher) coupled to a UHPLC system (Ultimate 3000 RSLCnano UHPLC, ThermoFisher) using positive NSI with the source temperature of 300°C and the spray voltage set to static at 2200V. The UHPLC was equipped with a 5 µL loop and reversed phase chromatographic separation was performed using a nanoflow column (50 cm x 75 μm, CoAnn Technologies, Richland, WA) self-packed with Luna C18 (5 μm,100 Å, Phenomenex). The mobile phases consisted of 5 mM NH_4_OAc (A) and 95% CH_3_CN in H_2_O (B) and the injection volume was 4 μL. The gradient started at 1% B for 20 min at a flow rate of 0.3 μL/min, followed by an increase to 5% over 5 min, then an increase to 22% over 35 min, followed by an increase to 95% over 1 min and held at these conditions for 2 min. The gradient was then returned to 1% B in 1 min and the column was re-equilibrated at this mobile phase composition for 3 min at a flow rate of 0.9 μL/min before the next injection for a full run time of 69 min. This targeted approach MS^2^ fragmentation (quadrupole isolation width of 1.5 m/z, HCD collision energy of 30%, AGC value of 1000%, maximum injection time of 200 ms, and Orbitrap resolution setting of 60,000) was performed on 33 *m/z* values: 249.093 *m/z*, 276.1343 *m/z*, 284.0338 *m/z*, 284.0746 *m/z*, 293.1167 *m/z*, 298.1147 *m/z*, 361.0637 *m/z*, 365.1013 *m/z*, 375.2234 *m/z*, 384.0895 *m/z*, 384.0975 *m/z*, 391.0741 *m/z*, 401.1119 *m/z*, 401.1128 *m/z*, 408.1085 *m/z*, 424.1021 *m/z*, 442.1132 *m/z*, 530.2819 *m/z*, 578.2565 *m/z*, 578.2565 *m/z*, 580.2092 *m/z*, 587.1613 *m/z*, 589.2773 *m/z*, 595.2197 *m/z*, 597.1568 *m/z*, 602.1562 *m/z*, 603.1555 *m/z*, 604.1405 *m/z*, 605.1959 *m/z*, 606.1998 *m/z*, 606.2908 *m/z*, 607.1010 *m/z*, and 607.2926 *m/z*. A 200-650 *m/z* full scan event (with a maximum injection time of 400 ms, an AGC value of 50%, and an Orbitrap resolution setting of 15,000) was included to monitor for any anomalies in sample composition or irregularities in the analysis.

### Untargeted LC-MS^2^/MS^3^ Screening

Samples from wild-type and Xpc -/-liver DNA were prepared in triplicates as described above. The samples were reconstituted in 20 μL of H_2_O, then all three 20 μL aliquots of the wild-type samples were combined into one vial. The same was done for the three 20 μL aliquots of the *Xpc -/-* samples. The analysis was done using an Orbitrap Lumos instrument (ThermoFisher) coupled to a UHPLC system (UltiMate 3000 RSLCnano UHPLC, ThermoFisher) using positive NSI with the source temperature at 300°C and the spray voltage at 2200V. The UHPLC was equipped with a 5 μL loop and reversed phase chromatographic separation was performed using a nanoflow column (50 cm x 75 μm ID, New Objective, Woburn, MA) self-packed with Luna C18 (5 μm,100 Å, Phenomenex). The mobile phases consisted of 5 mM NH_4_OAc (A) and 95% CH_3_CN in H_2_O (B) and the injection volume was 4 μL. The gradient started at 1% B for 20 min at a flow rate of 0.3 μL/min, followed by an increase to 5% over 5 min, then an increase to 22% over 35 min, followed by an increase from to 95% over 1 min and held at these conditions for 2 min, the gradient was then returned to 1% B in 1 min and the column was re-equilibrated at this mobile phase composition for 3 min at a flow rate of 0.9 μL/min before the next injection for a full run time of 69 min. The samples were injected four separate times with each injection using a different mass range (range 1: 145-288 *m/z*, range 2: 283-426 *m/z*, range 3: 421-564 *m/z*, and range 4: 559-702 *m/z*) with a maximum injection time of 250 ms, an AGC value of 1250%, and a resolution setting of 120,000. Data dependent parameters included a mass tolerance of ± 5 ppm, a repeat count of 1, a dynamic exclusion of 5 s, a minimum intensity of 5.0e3, and a cycle time of 2 s. MS^2^ fragmentation involved a quadrupole isolation width of 1.5 m/z, stepped HCD collision energy of 15, 30, and 45%, an AGC value of 1000%, a maximum injection time of 100 ms, and a resolution setting of 15,000. MS^2^ product ions were isolated in the ion trap with a 2 m/z isolation window, and MS^3^ fragmentation was triggered upon observation of the neutral loss of 2ʹ-deoxyribose (-dR; 116.0474 Da), the base moieties (-G; 151.0494 Da, -A; 135.0545 Da, -T; 126.0429 Da; -C; 111.0433 Da), or base moieties plus water (-G+H_2_O; 169.0646 Da, -A+H_2_O; 153.0651 Da, -T+H_2_O; 144.0535 Da, -C+H_2_O; 129.0538 Da).

### LC-MS^2^/MS^3^ Data Analysis using Compound Discoverer (CD)

The data generated from the untargeted analysis on the Orbitrap Lumos instrument was imported into CD (ThermoFisher) which provides analyte identification, characterization and comparative analyses between sample groups. CD generated a list of all the potential compounds present in both the wild-type and *Xpc -/-* liver samples, the list consisted of a total of 65,108 potential compounds. Filters were then implemented based on the following criteria: peak area ratio of *Xpc -/-* over wild-type greater than 1.00, presence of guanine product ion in MS^2^ spectra (provided by the Compound Class node in CD), or neutral loss of either deoxyribose, guanine, cytosine, adenine, thymine, or any of those four base moieties plus water (provided by the Neutral Loss node in CD). With these filters implemented the list of potential compounds decreased to 285. All 285 compounds were then manually confirmed using Xcalibur Freestyle software (ThermoFisher). The manual confirmation resulted in 33 of the 285 showing a peak area ≥ 1.5 times higher in *Xpc -/-* than the wild-type sample, a Gaussian shaped peak, and similar retention times in both sample groups. The parameters used to generate the list of putative adducts are illustrated in **Supplementary Figure 10** and listed below.

**Supplementary Table 3.**
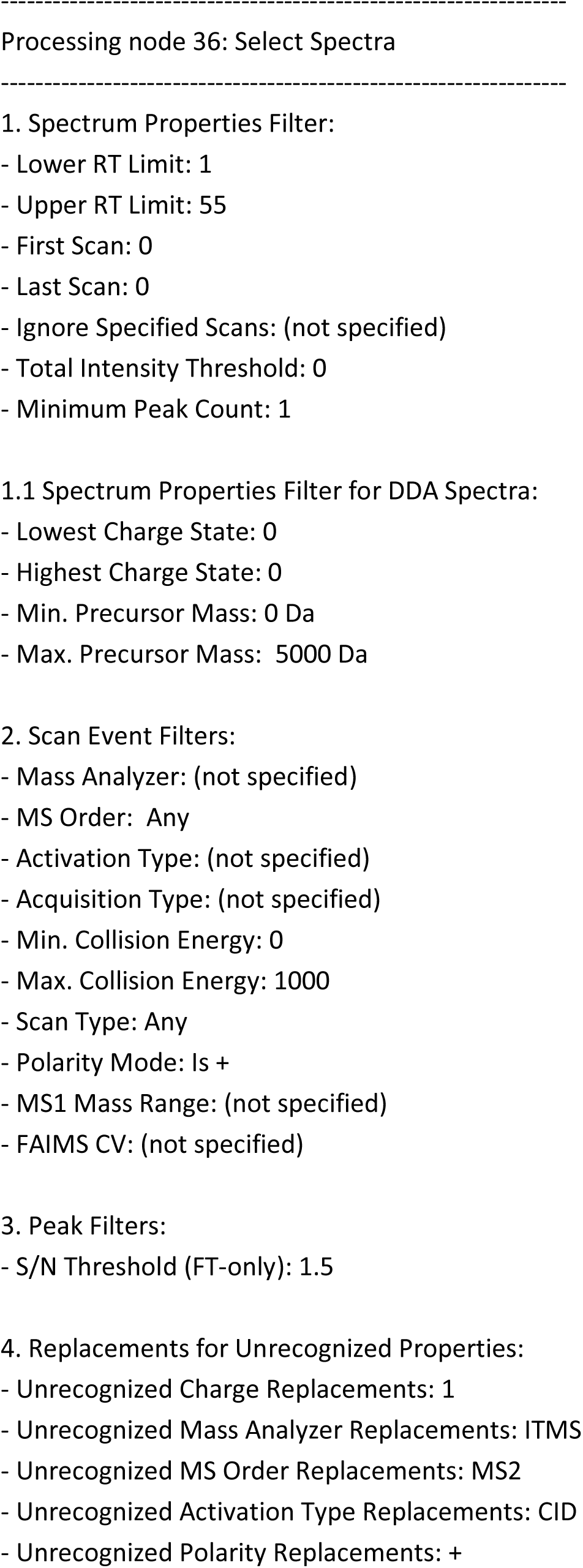
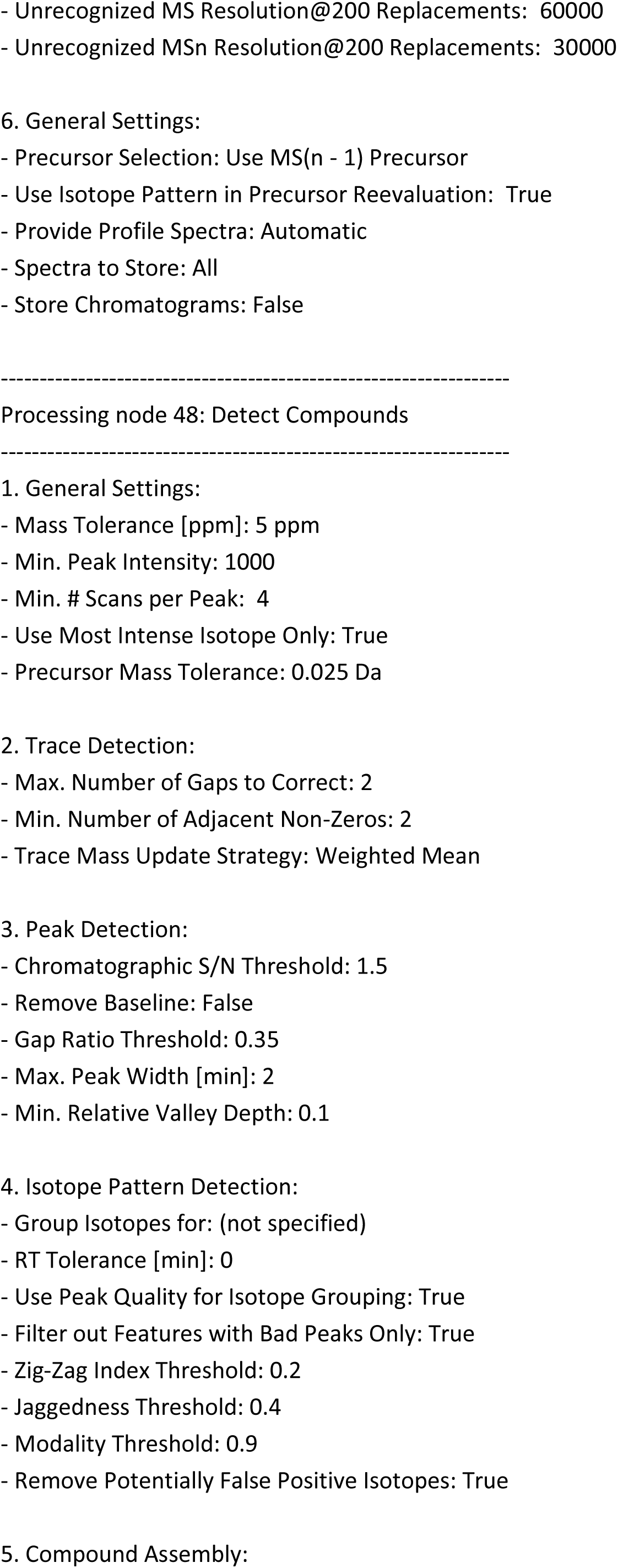
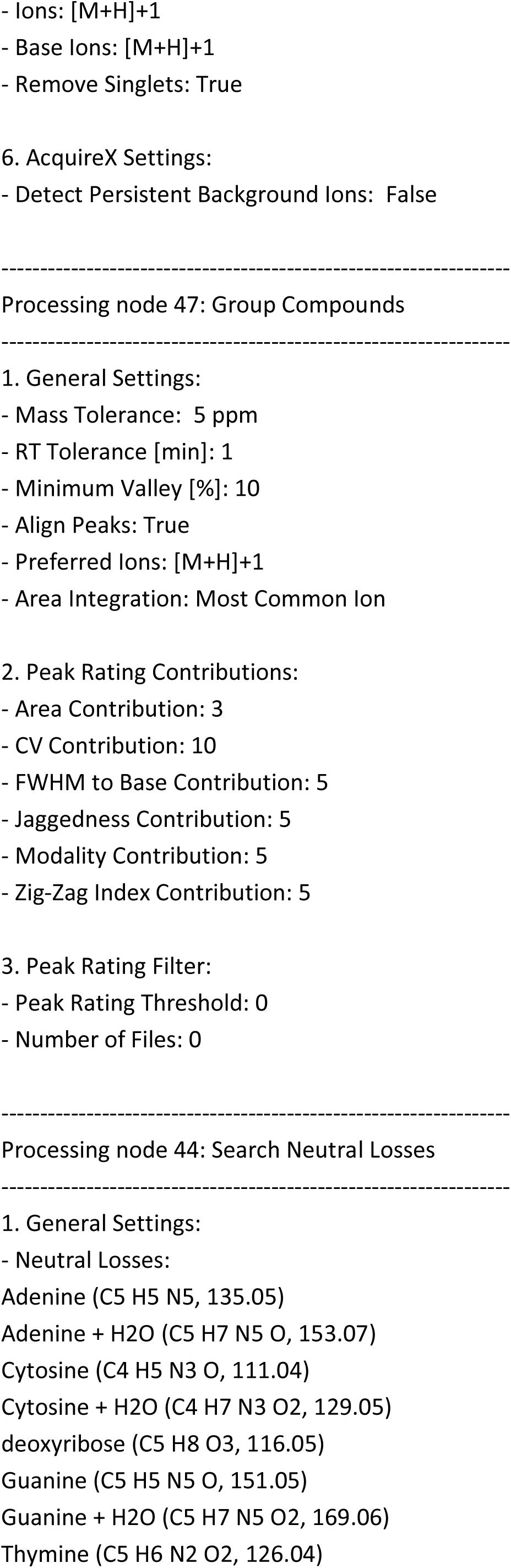
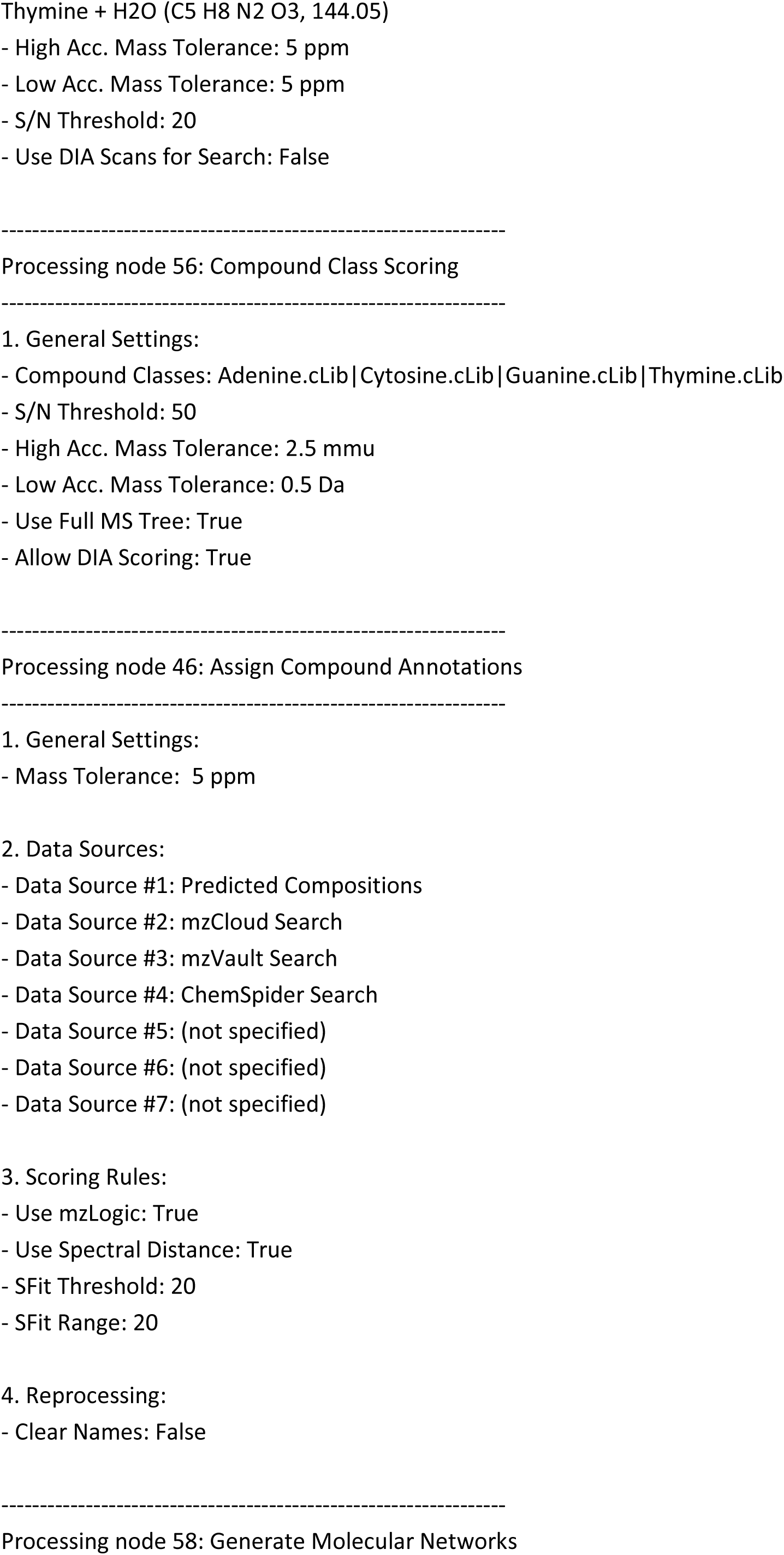
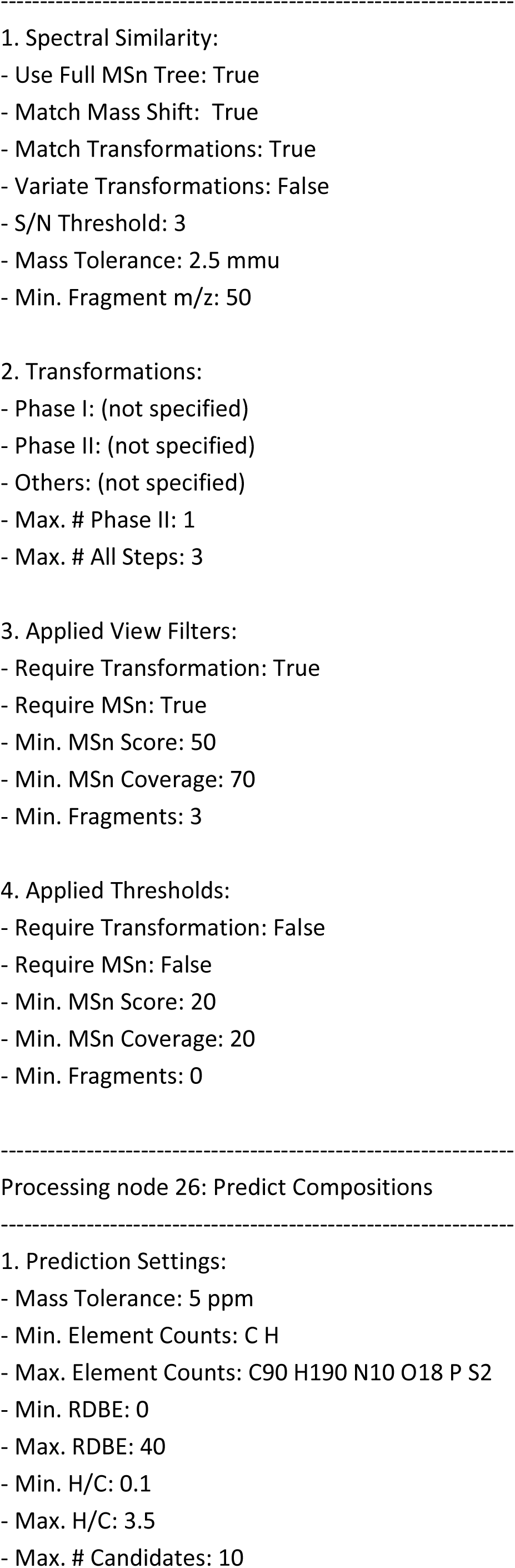
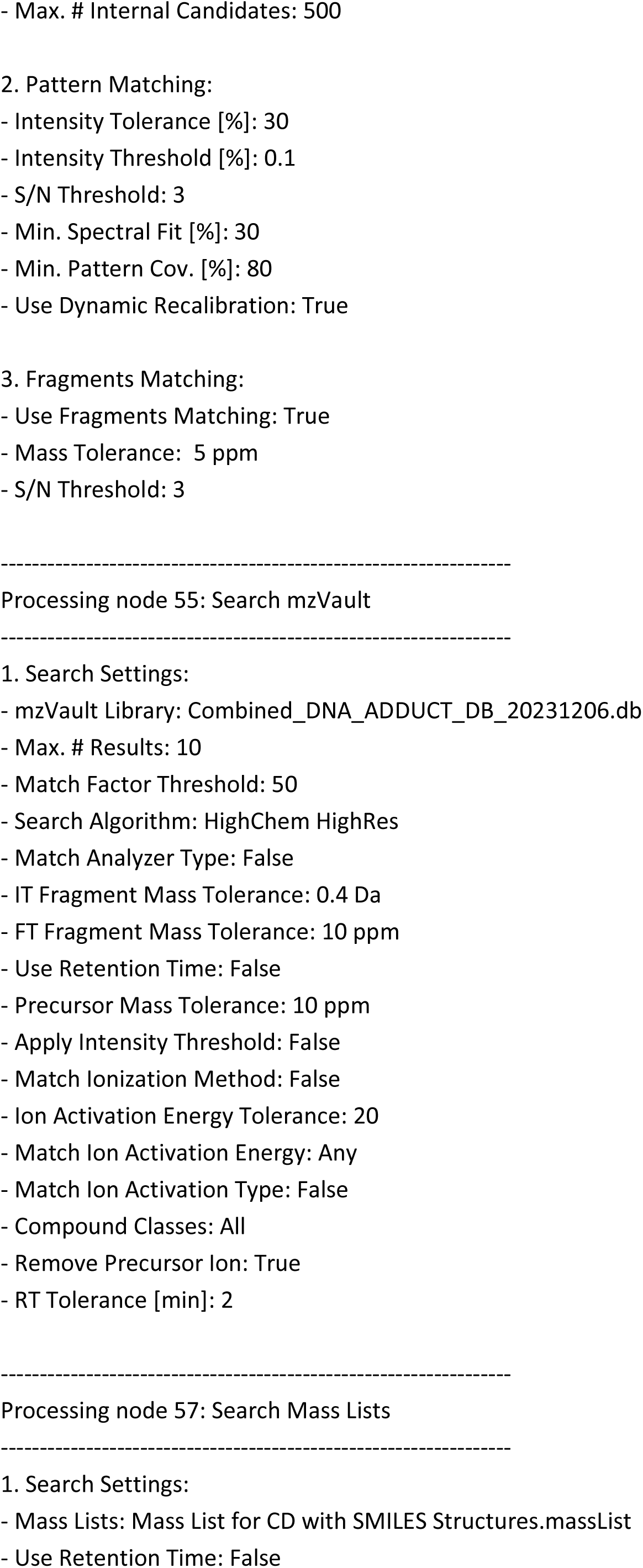
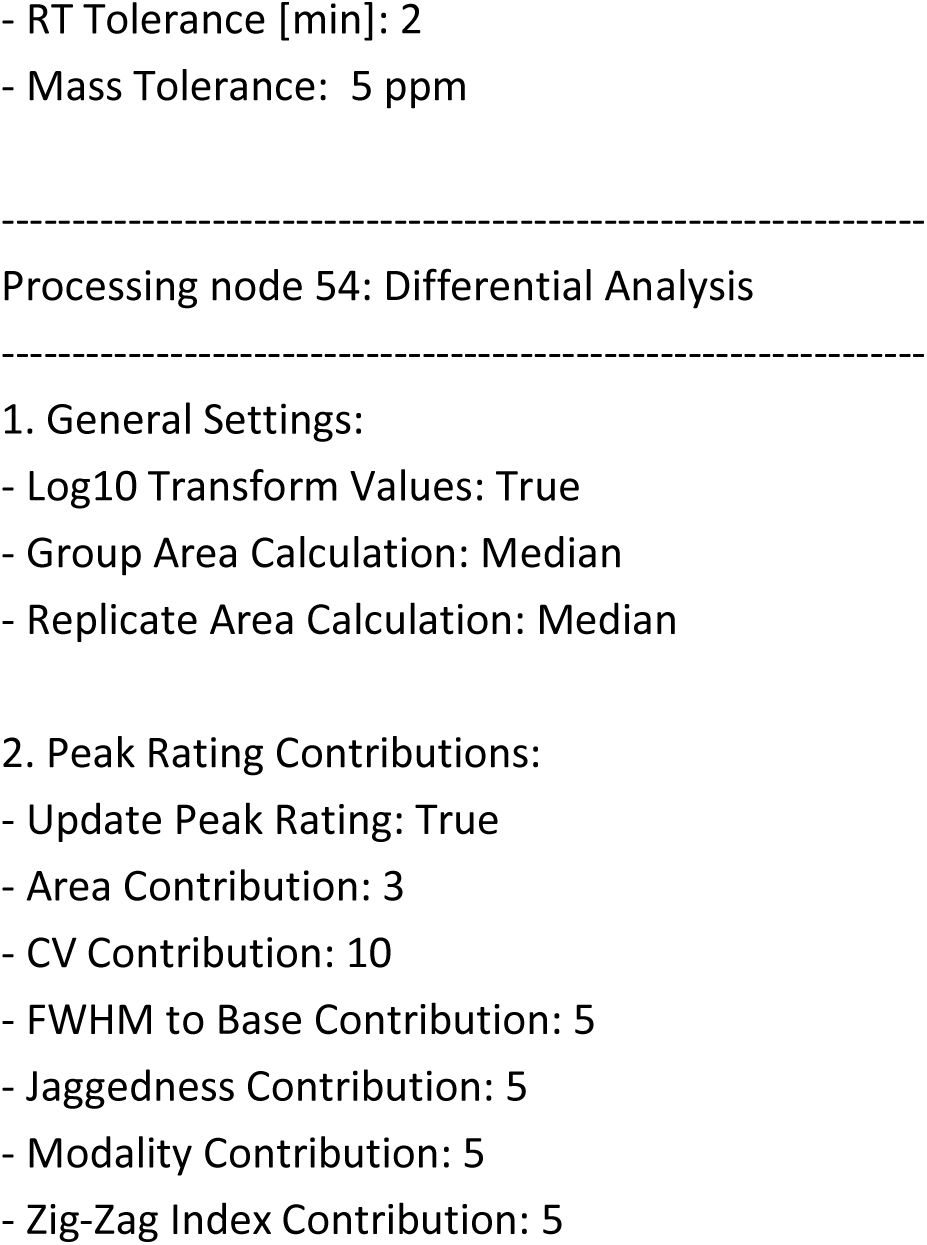

## Figure legends

**Supplementary Figure 1.**
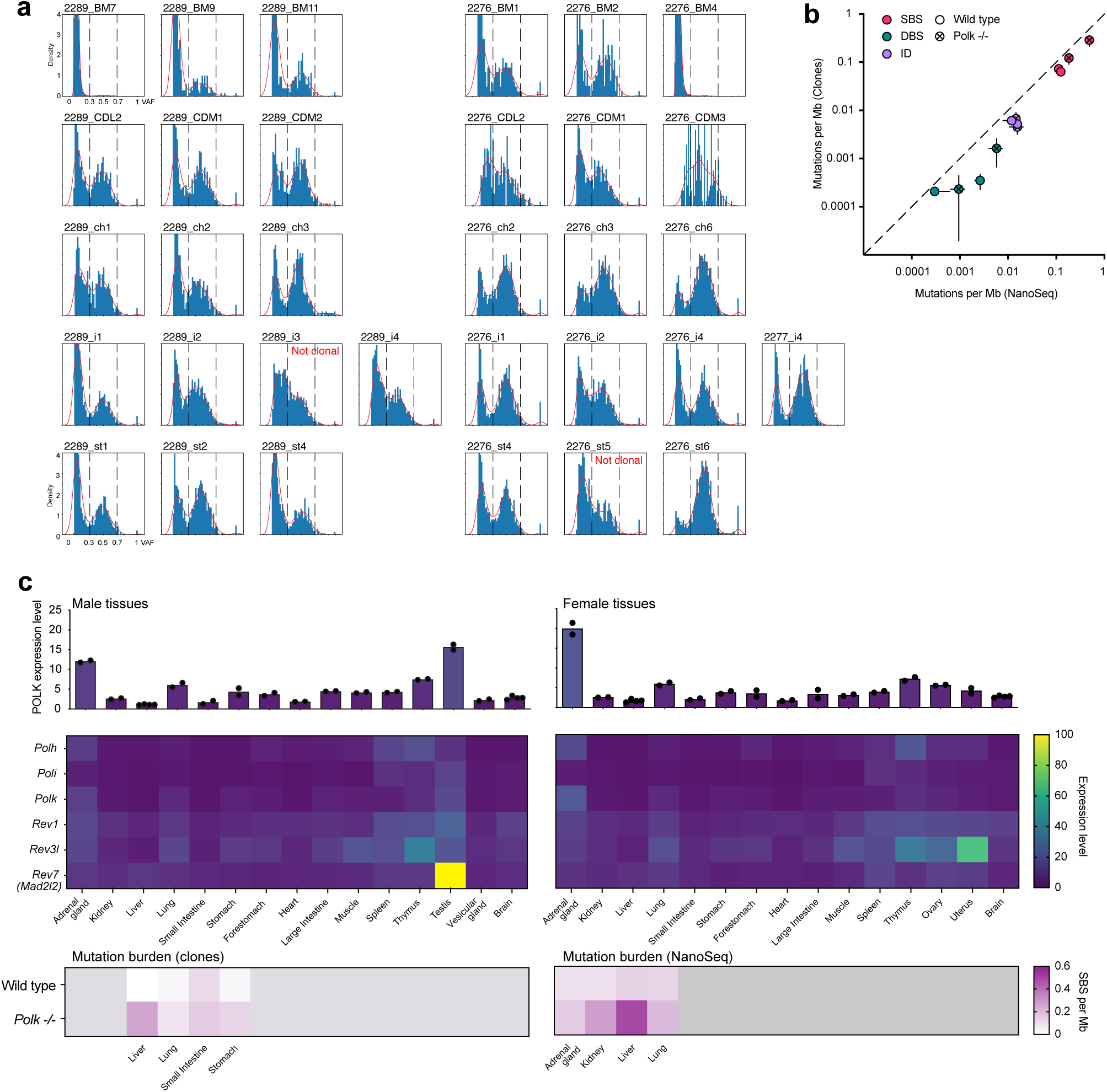
Characterizing mutation in mouse tissues. **a)** Variant allele frequency (VAF) distribution of single cell clones used in Figure 1. Samples with a clear peak centered around VAF = 0.5 were considered clonal and used in subsequent analyses. Samples with no distinct peak or a peak with VAF < 0.5 were considered ‘non-clonal’ and excluded from analysis. **b)** Comparison of mutations burden between clone whole-genome sequencing and bulk NanoSeq data. Burden of single-base substitutions (SBSs), doublet base substitutions (DBSs) and indels (ID), for aged wild type and *Polk-/-*, lung and liver tissue. Dots represent the mean of *n* = 3 genomes or samples, and lines standard deviation. **c)** Comparison between SBS mutation burden and bulk RNA expression of Polk and other TLS polymerases. Expression data (FPKM normalized) is derived from Li *et al*., Sci Rep. 2017.

**Supplementary Figure 2.**
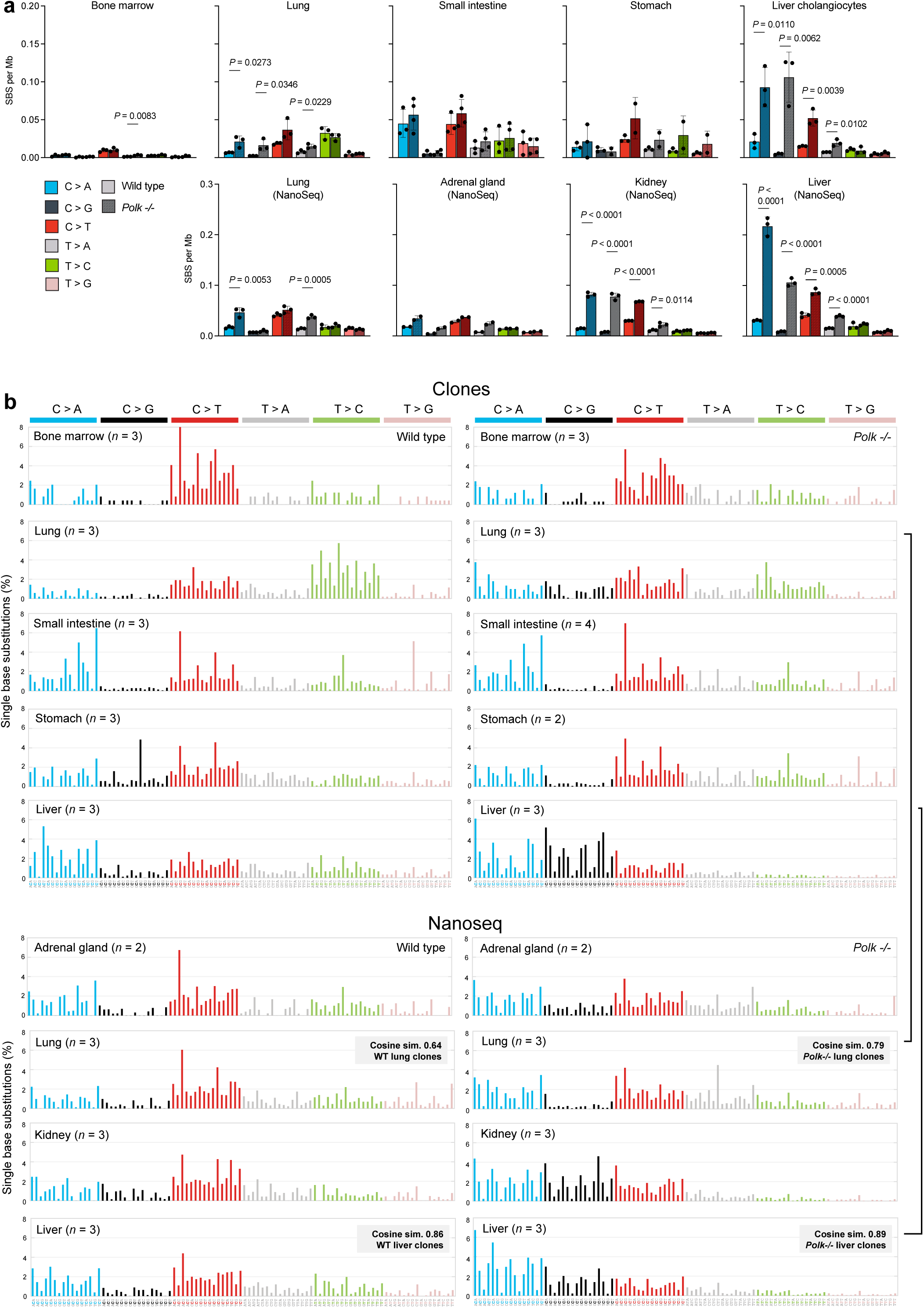
Overview of Single Base Substitutions (SBS) across tissues. **a)** Burden of the six classes of SBSs, in wild type and *Polk-/-* mice (*P* calculated by an unpaired *t* test). **b)** 96-classes of SBSs considering the six mutation types but also the bases immediately 5’ and 3’ of the mutated base. Each graph represents the average mutation pattern for *n* genomes or samples, where *n* is indicated in each panel.

**Supplementary Figure 3.**
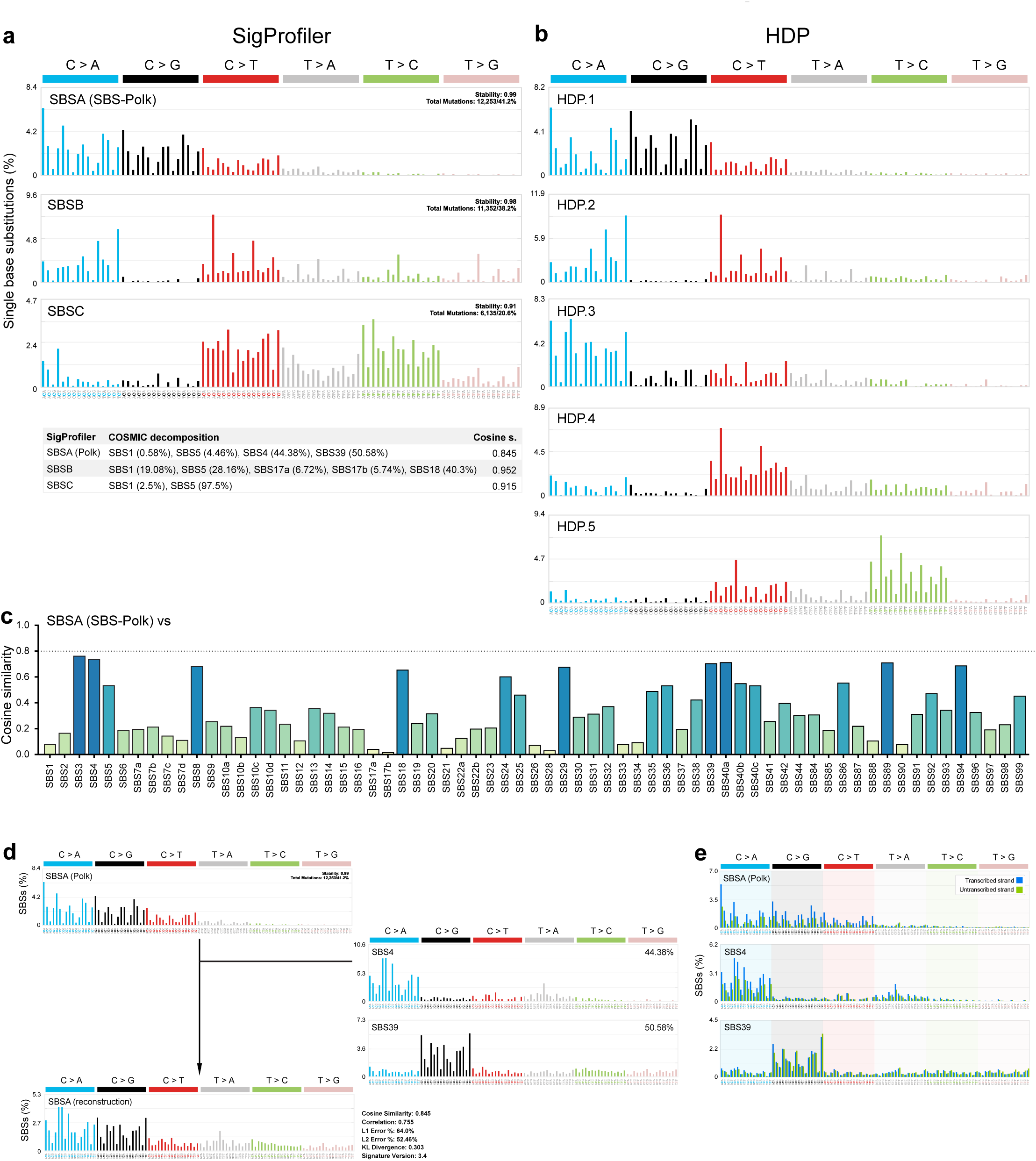
Extraction of Single Base Substitutions (SBS) mutational signatures. **a)** *De novo* mutational signature extraction using non-negative matrix factorization (NMF) with SigProfilerExtractor yielded three signatures (SBSA-C). The bottom table shows the decomposition of these signatures into known signatures of the Catalogue of Somatic Mutations in Cancer (COSMIC). **b)** *De novo* mutational signature extraction using hierarchical Dirichlet process (HDP) with mSigHdp yielded five signatures (HDP.1-5). SBSA and HDP.1 signatures have a Cosine similarity of 0.96 showing an identical signature was extracted by two methods. **c)** Comparison of the Cosine similarities between **SBSA** and known COSMIC signatures v3.4. **d)** Reconstruction of SBSA using a combination of SBS4 and SBS39 by SigProfiler. **e)** Plots showing the percentage of mutations in transcribed and untranscribed strands for each mutational signature in the 96 mutational contexts.

**Supplementary Figure 4.**
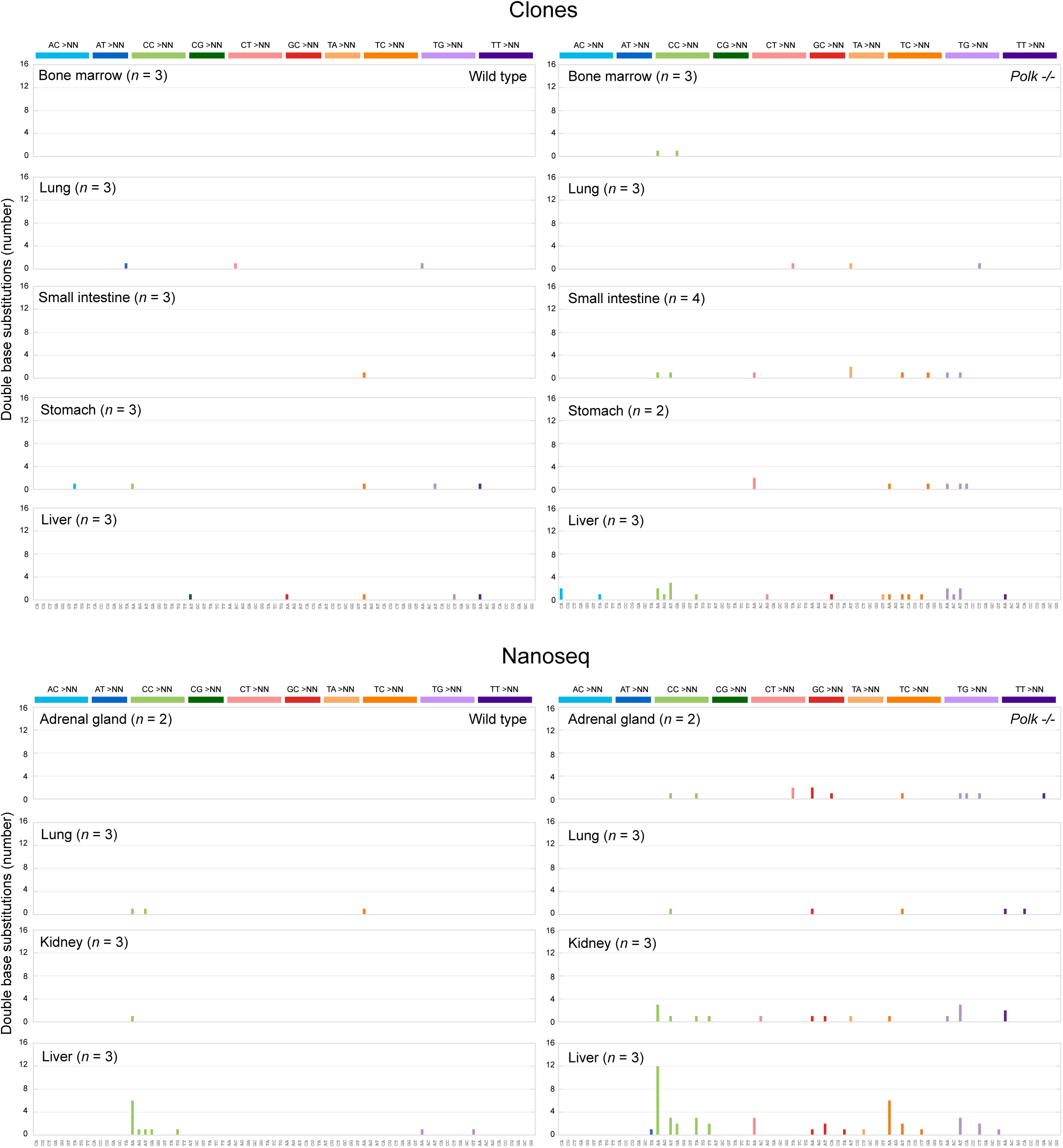
Overview of Doublet Base Substitutions (DBS) across tissues. 78-classes of DBSs according to the COSMIC classification. Each graph represents the average mutation pattern for *n* genomes or samples, where *n* is indicated in each panel.

**Supplementary Figure 5.**
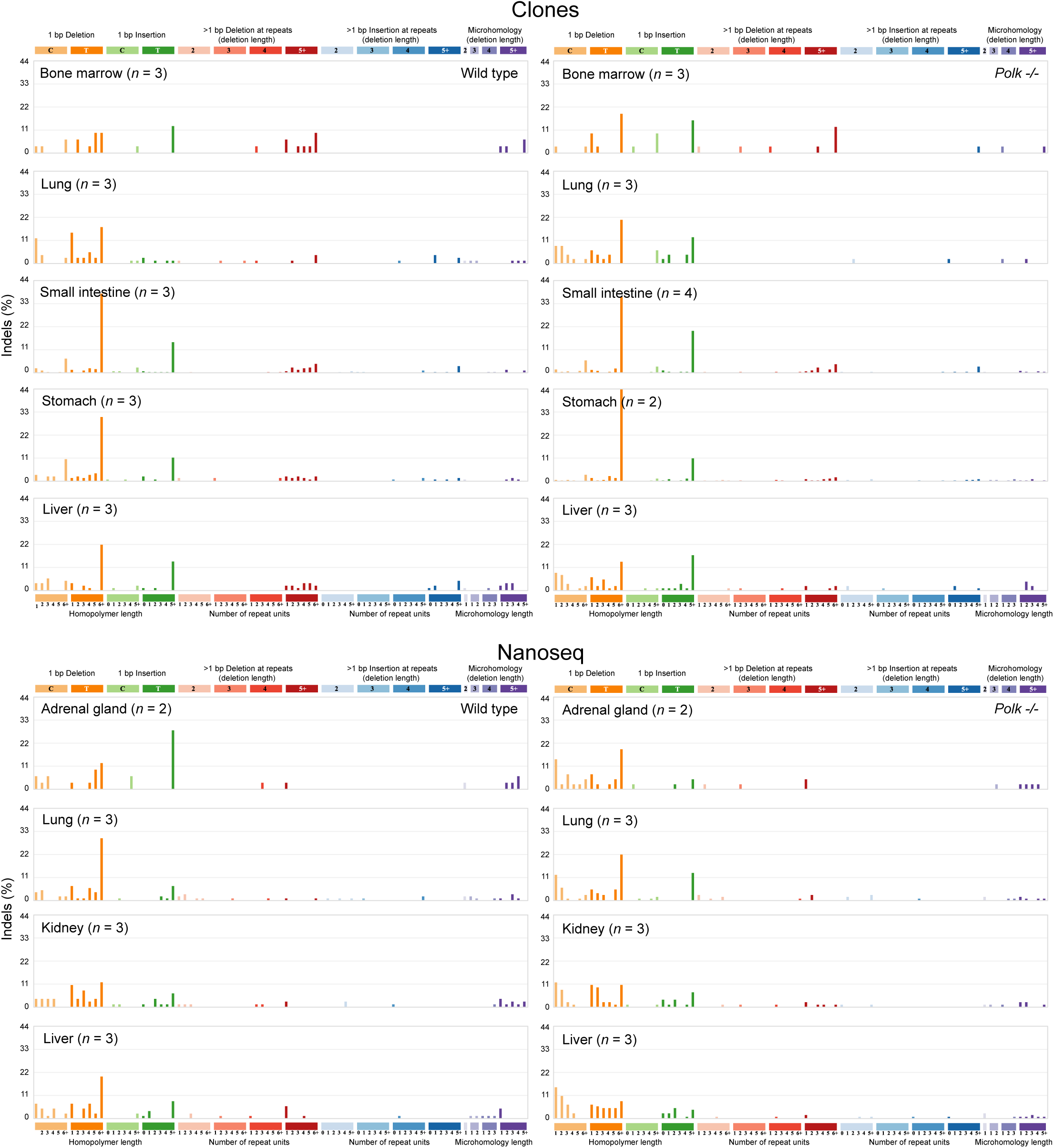
Overview of Indels across tissues. 83-classes of insertions and deletions (indels, ID) according to the COSMIC classification, which considers size, nucleotides affected and presence on repetitive and/or microhomology regions. Each graph represents the average mutation pattern for n genomes or samples, where n is indicated in each panel.

**Supplementary Figure 6.**
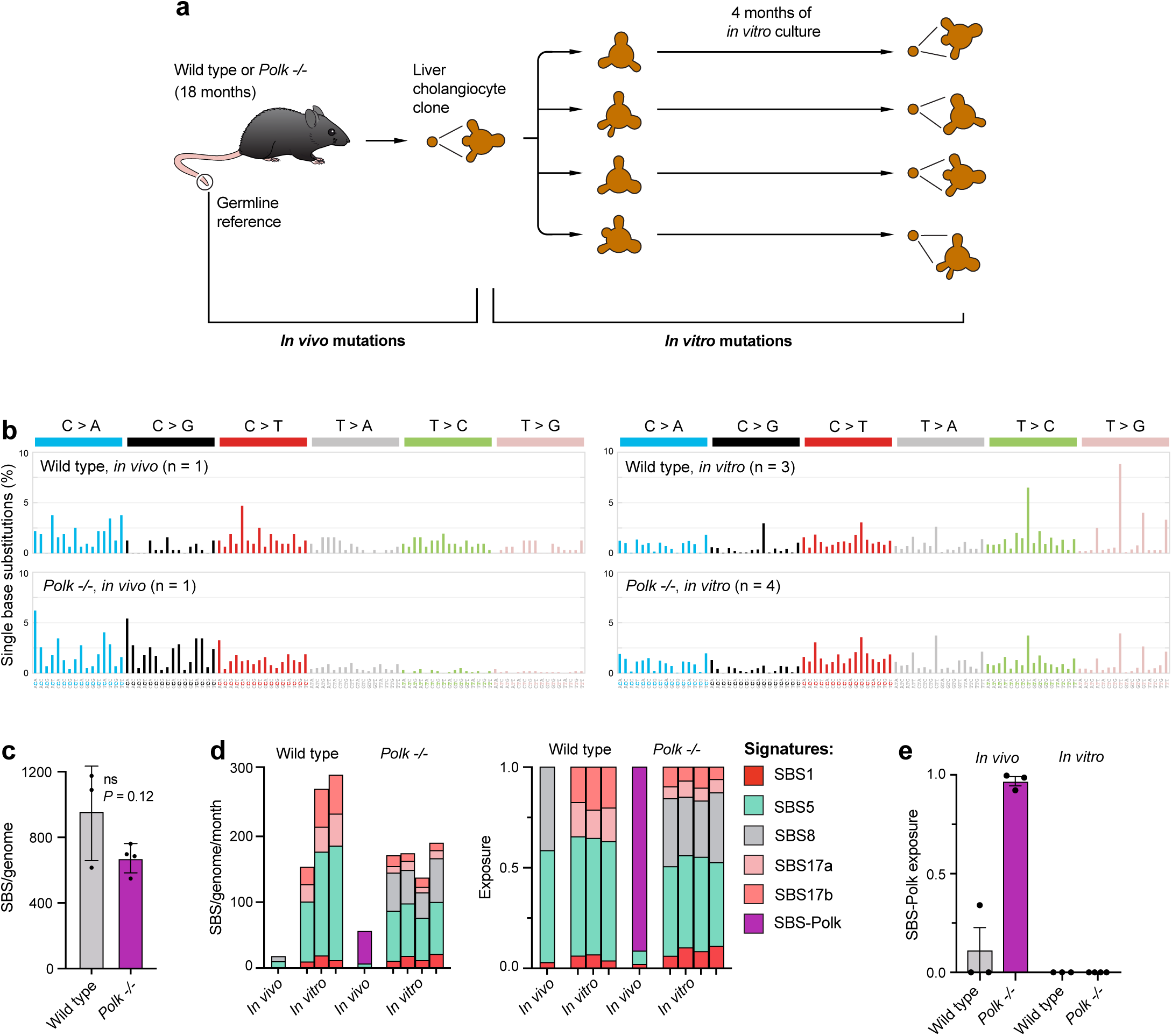
SBS-Polk mutations do not accumulate during *in vitro* culture. **a)** Scheme for the characterization of mutations during in vitro culture of liver cholangiocytes. A wild type or *Polk-/-* clone (from Figure 1) were split into 4 cultures, passaged for 4 months, sub-cloned and subjected to whole genome sequencing. Mutations present at the beginning of the experiment were removed during variant calling. **b)** 96-classes of single-base substitutions (SBSs) for the initial clone and the clones at the end of the *in vitro* culture. Each graph represents the average mutation pattern of *n* = 1-4 genomes. **c)** Quantification of SBSs substitutions at the end of the experiment. **d)** Extraction and assignment of mutational signatures using SigProfiler Extractor. Stacked bar plots showing estimated number (left) and proportion (right) of each mutational signature in individual clones. **e)** Contribution of the SBS-Polk mutational signatures to the mutational landscapes of liver cholangiocytes grown *in vivo* or *in vitro*.

**Supplementary Figure 7.**
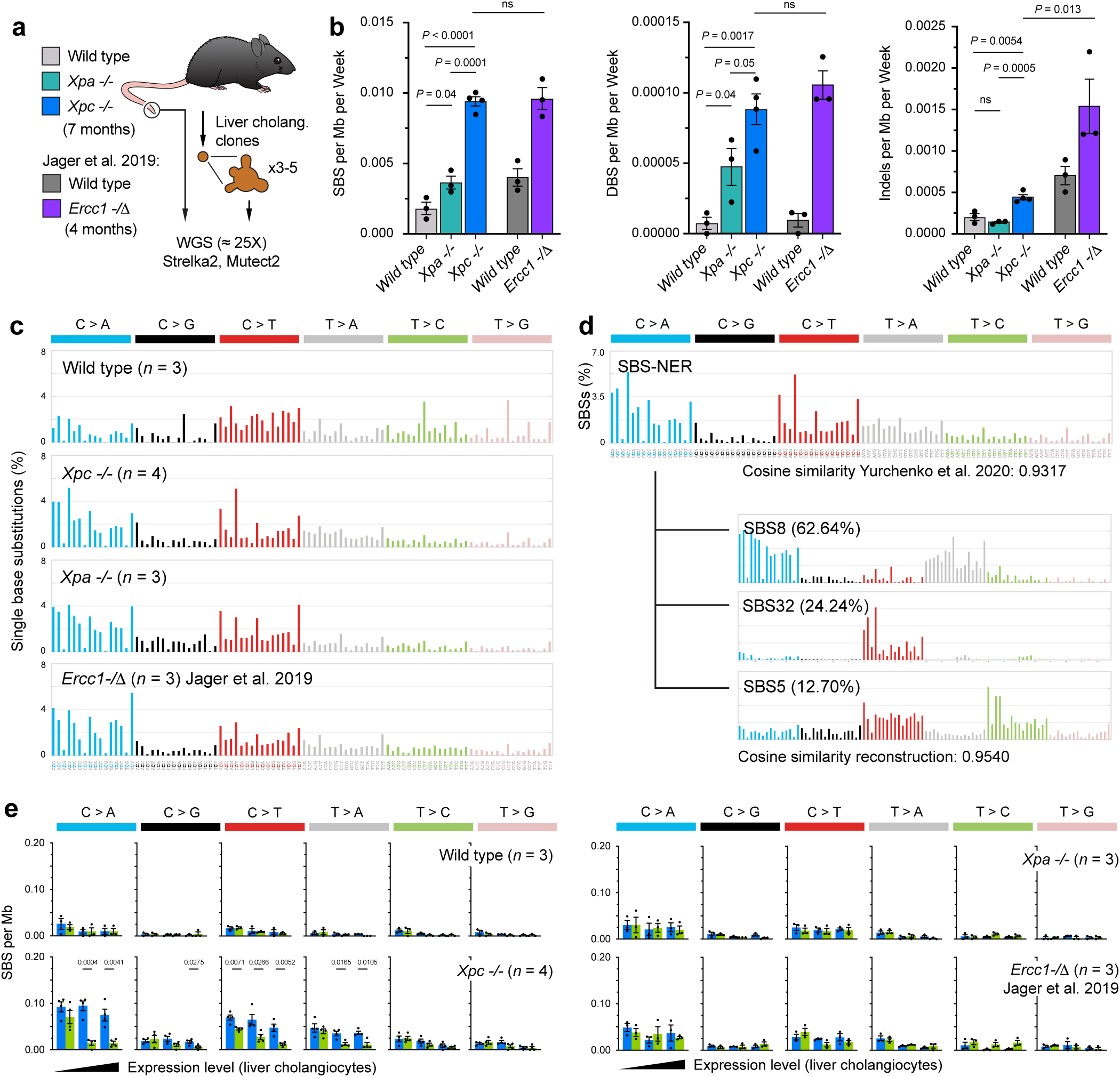
Mutational consequences of NER deficiency in liver cholangiocytes. **a)** Single cholangiocytes were isolated from 7-month-old wild type, *Xpc-/-* and *Xpa-/-* mice, and expanded into clonal organoid lines, which were subjected to whole-genome sequencing and variant calling. For comparison, we reanalysed published data 4-month-old *Ercc1-/Δ* liver cholangiocytes (Jager *et al*. 2019). **b)** Number of single-base substitutions (SBSs), doublet base substitutions (DBSs) and insertions/deletions (indels) per genome (*P* calculated by an unpaired *t* test). **c)** 96-classes of SBSs considering the six mutation types but also the bases immediately 5’ and 3’ of the mutated base. Each graph represents the average mutation pattern for *n* genomes, where *n* is indicated in each panel. **d)** Mutational signature of NER deficiency (SBS-NER), which is almost identical to the mutational signature of XPC leukaemias (Yurchenko *et al*. 2020), and can be further decomposed into known COSMIC signatures SBS8, SBS32 and SBS5. **e)** Relationship between transcriptional strand bias and expression level for mutations in liver cholangiocyte clones in NER-deficient samples (*P* calculated by unpaired *t* tests).

**Supplementary Figure 8.**
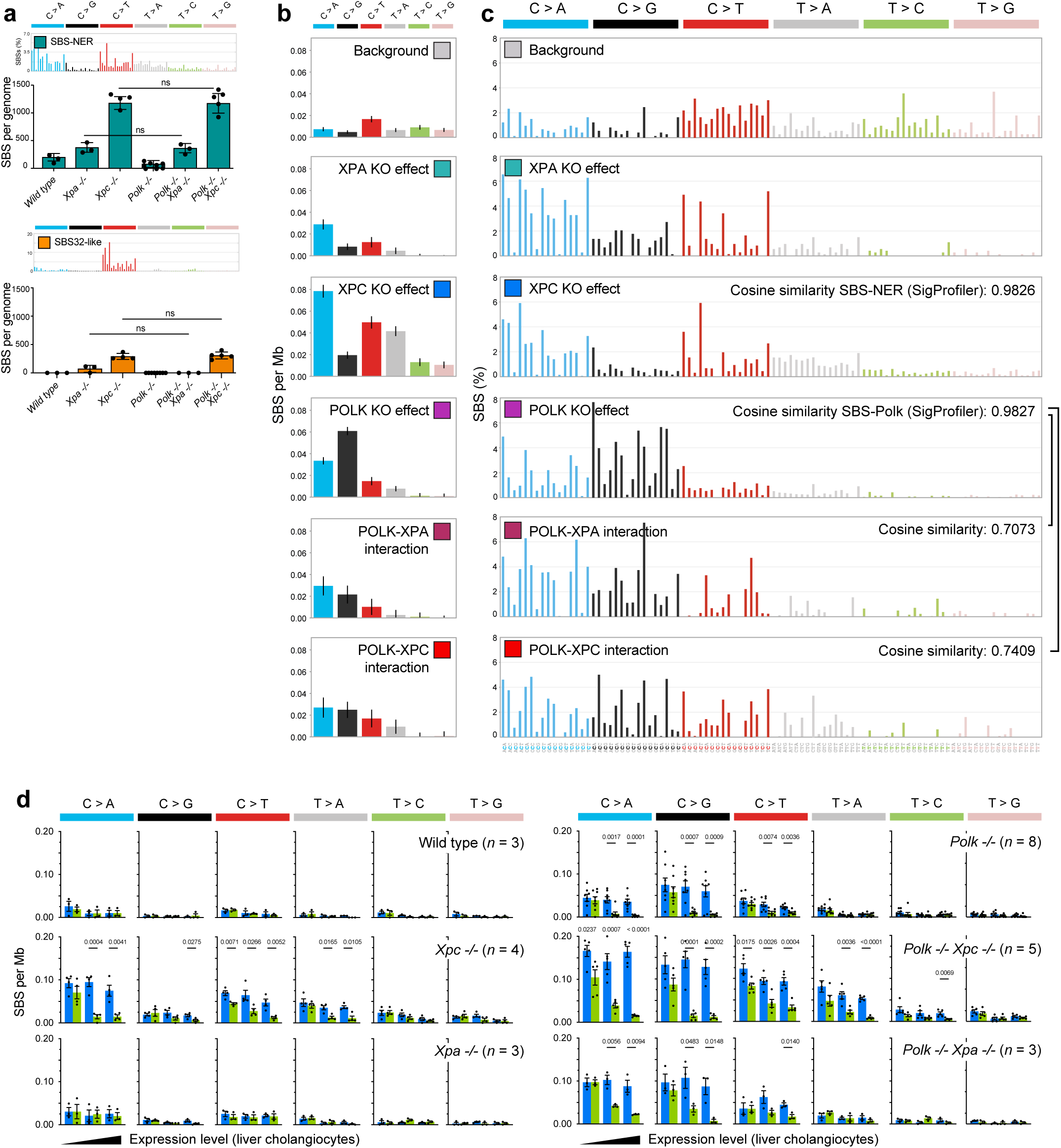
Mutations resulting from the Polk-NER interaction. **a)** Quantification of SigProfiler mutational signatures, SBS-NER and SBS32-like mutational signatures in *Xpc-/-Polk-/-* and control samples, related to Figure 4e (*P* calculated by unpaired *t* tests). **b**) 6-class substitution profiles of the mutations induced by gene loss in mouse livers, estimated with binomial regression, related to Figure 4f. The mutational burdens induced by gene losses are estimated by fitting a model for each substitution type. The interaction effects represent the mutational burden induced by combined gene loss on top of the individual gene loss effects. Bar heights represent the estimated effects, the vertical lines represent the Bonferroni-corrected confidence intervals. **c**) 96-class trinucleotide substitution profiles estimated with binomial regression per trinucleotide mutation class. As in b), the mutational profiles of the interaction effects represent the mutations induced by combined gene loss on top of those induced by individual gene losses. Bar heights represent the estimated percentage of total mutations assigned to each trinucleotide mutation class. **d)** Relationship between transcriptional strand bias and expression level for mutations in liver cholangiocyte clones in NER, Polk and NER/Polk-deficient samples (*P* calculated by unpaired *t* tests). Note the presence of transcriptional-strand bias in *Polk-/-* genomes also lacking Xpa.

**Supplementary Figure 9.**
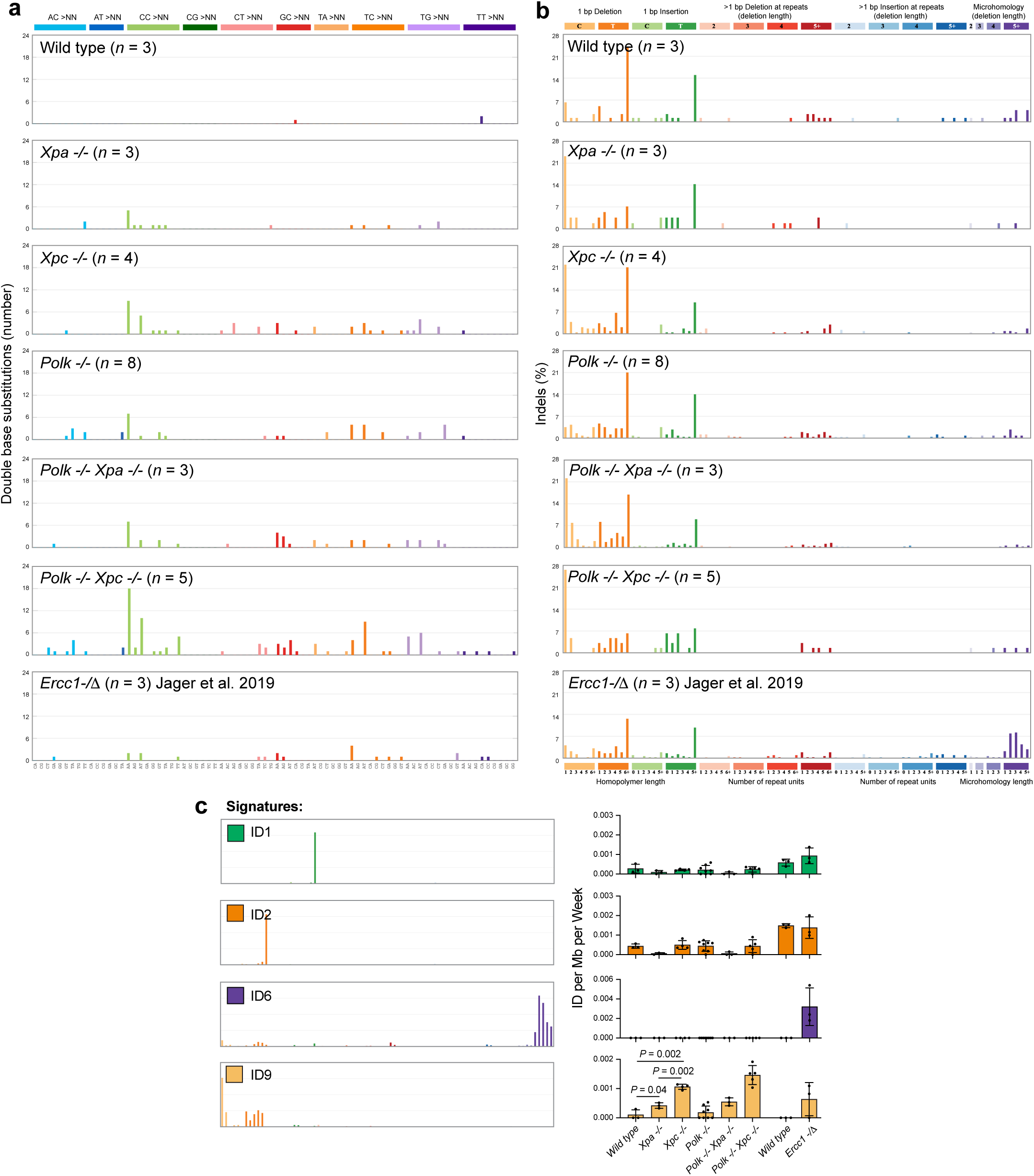
Doublet Base Substitutions (DBS) and Indels in mice lacking NER and Polk. **a)** Pattern of DBSs, following the 78-type classification from the COSMIC database. Each graph represents the average mutation pattern for *n* cholangiocyte genomes, where *n* is indicated in each panel. **b)** Pattern of insertions and deletions in bulk liver DNA, following the 83-type classification from the COSMIC database. Each graph represents the average mutation pattern for *n* cholangiocyte genomes, where *n* is indicated in each panel. **c)** Extraction of indel mutational signatures using SigProfilerExtractor. Stacked bar plots showing estimated number of each mutational signature in individual clones.

**Supplementary Figure 10.**
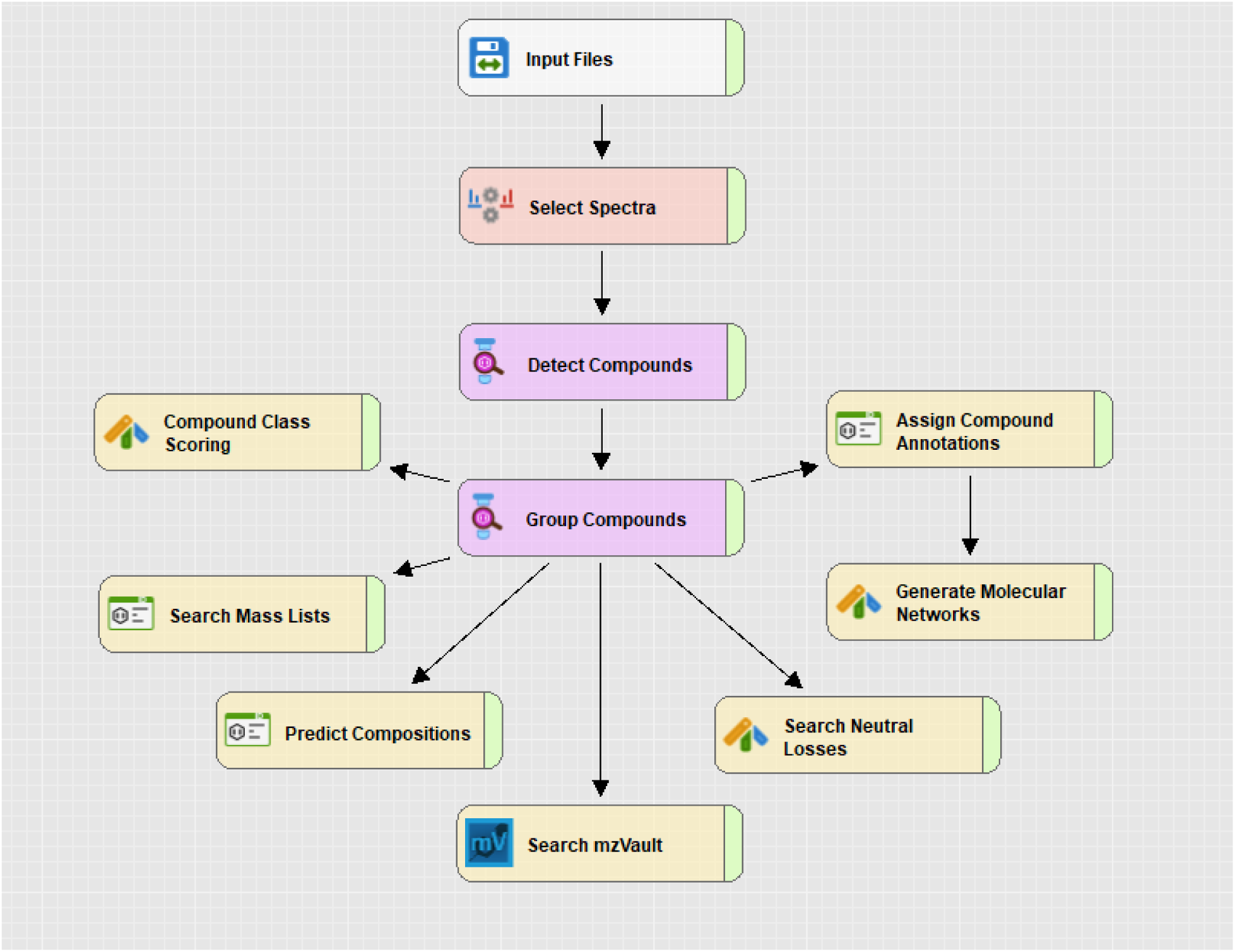
Visual representation of the workflow used to generate the list of putative adducts in Compound Discoverer. Each box represents a different processing node. The parameters for each node are detailed in Supplementary Table 3.

**Supplementary Figure 11.**
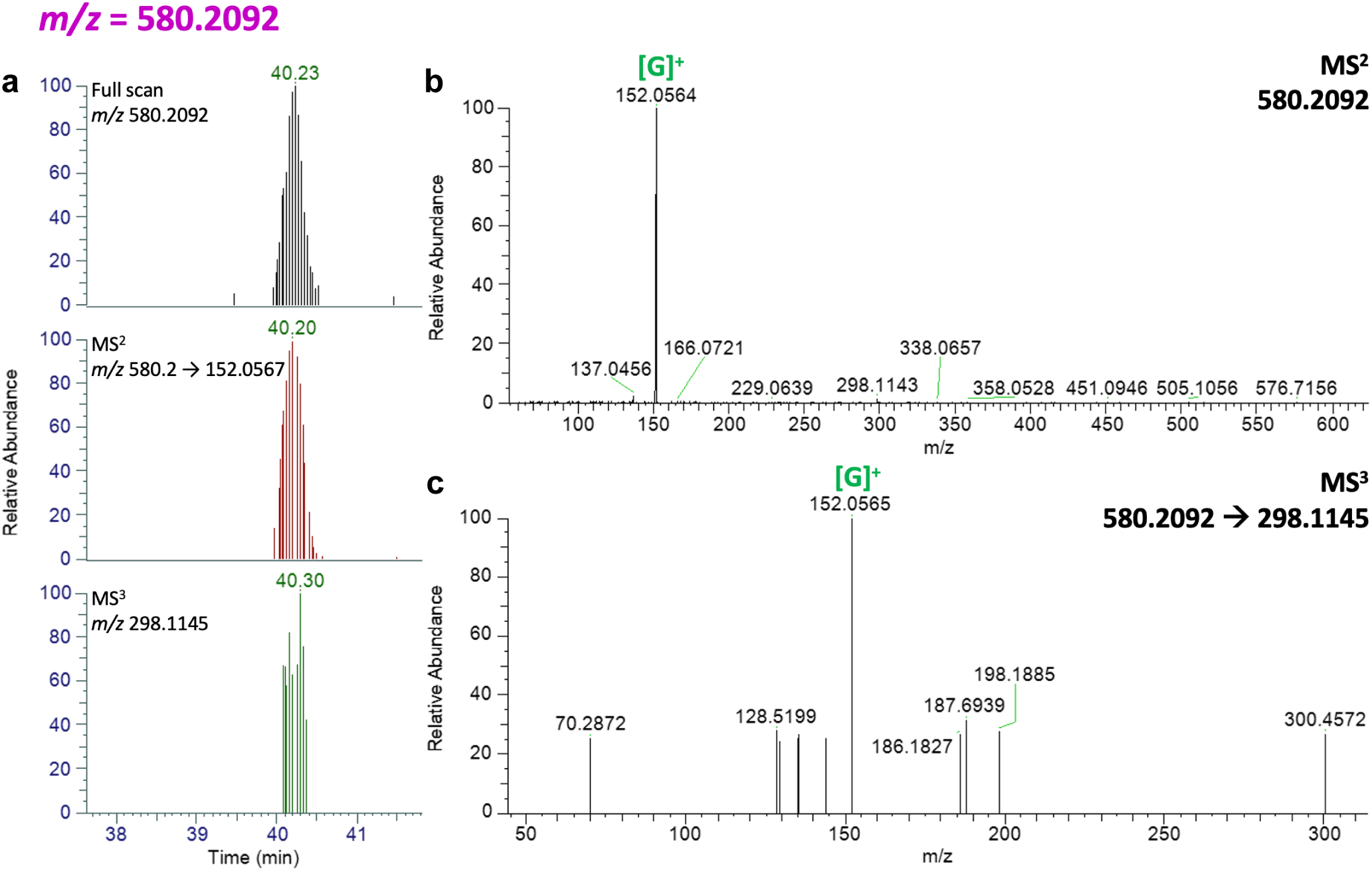
Mass Spectrometry data for the adduct with *m/z* 580.2092. **a)** Extracted chromatogram for the adduct in the full scan and the co-eluting peaks corresponding to the fragments detected in the MS^2^ and the MS^3^ fragmentation events. **b)** Spectrum corresponding to the MS^2^ fragmentation event with the mass of a known fragment highlighted. **c)** Spectrum of the MS^3^ fragmentation event triggered by one of the major fragments appearing in the MS^2^ spectrum, showing the appearance of Guanine.

**Supplementary Figure 12.**
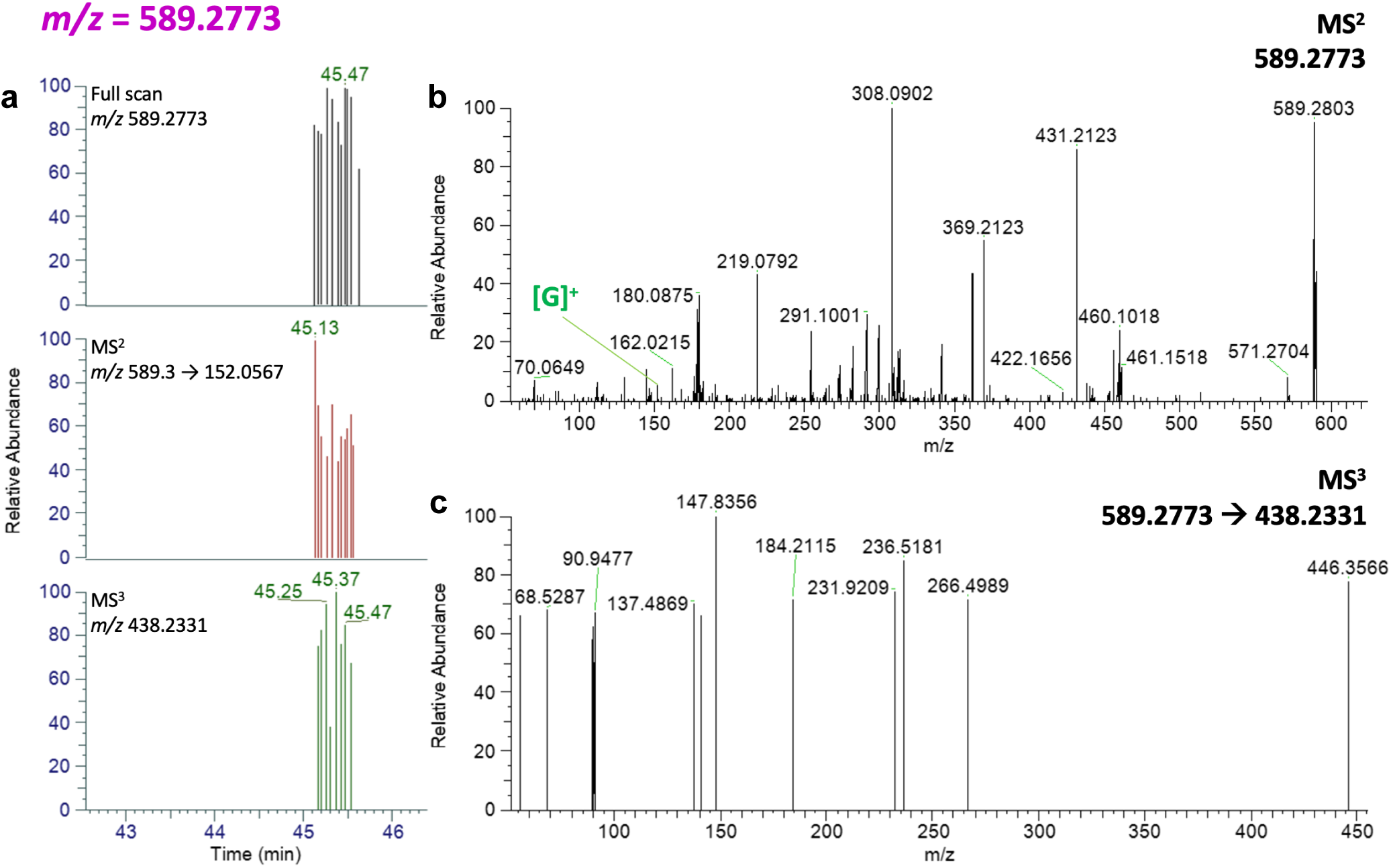
Mass Spectrometry data for the adduct with *m/z* 589.2773. **a)** Extracted chromatogram for the adduct in the full scan and the co-eluting peaks corresponding to the fragments detected in the MS^2^ and the MS^3^ fragmentation events. **b)** Spectrum corresponding to the MS^2^ fragmentation event with the mass of a known fragment highlighted. **c)** Spectrum of the MS^3^ fragmentation event triggered on one of the major fragments appearing in the MS^2^ spectrum.

**Supplementary Figure 13.**
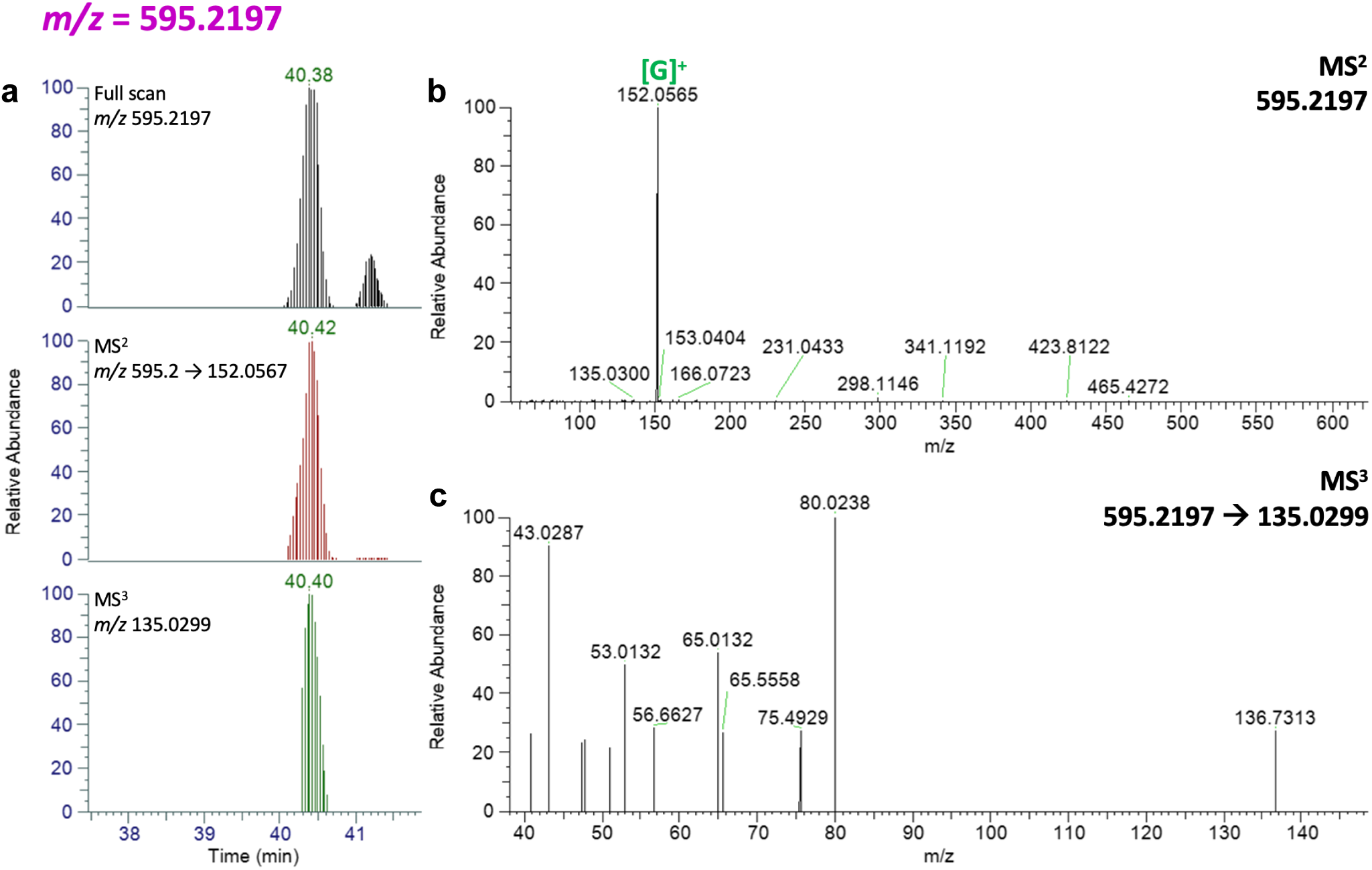
Mass Spectrometry data for the adduct with *m/z* 595.2197. **a)** Extracted chromatogram for the adduct in the full scan and the co-eluting peaks corresponding to the fragments detected in the MS^2^ and the MS^3^ fragmentation events. **b)** Spectrum corresponding to the MS^2^ fragmentation event with the mass of a known fragment highlighted. **c)** Spectrum of the MS^3^ fragmentation event triggered by one of the major fragments appearing in the MS^2^ fragmentation event.

**Supplementary Figure 14.**
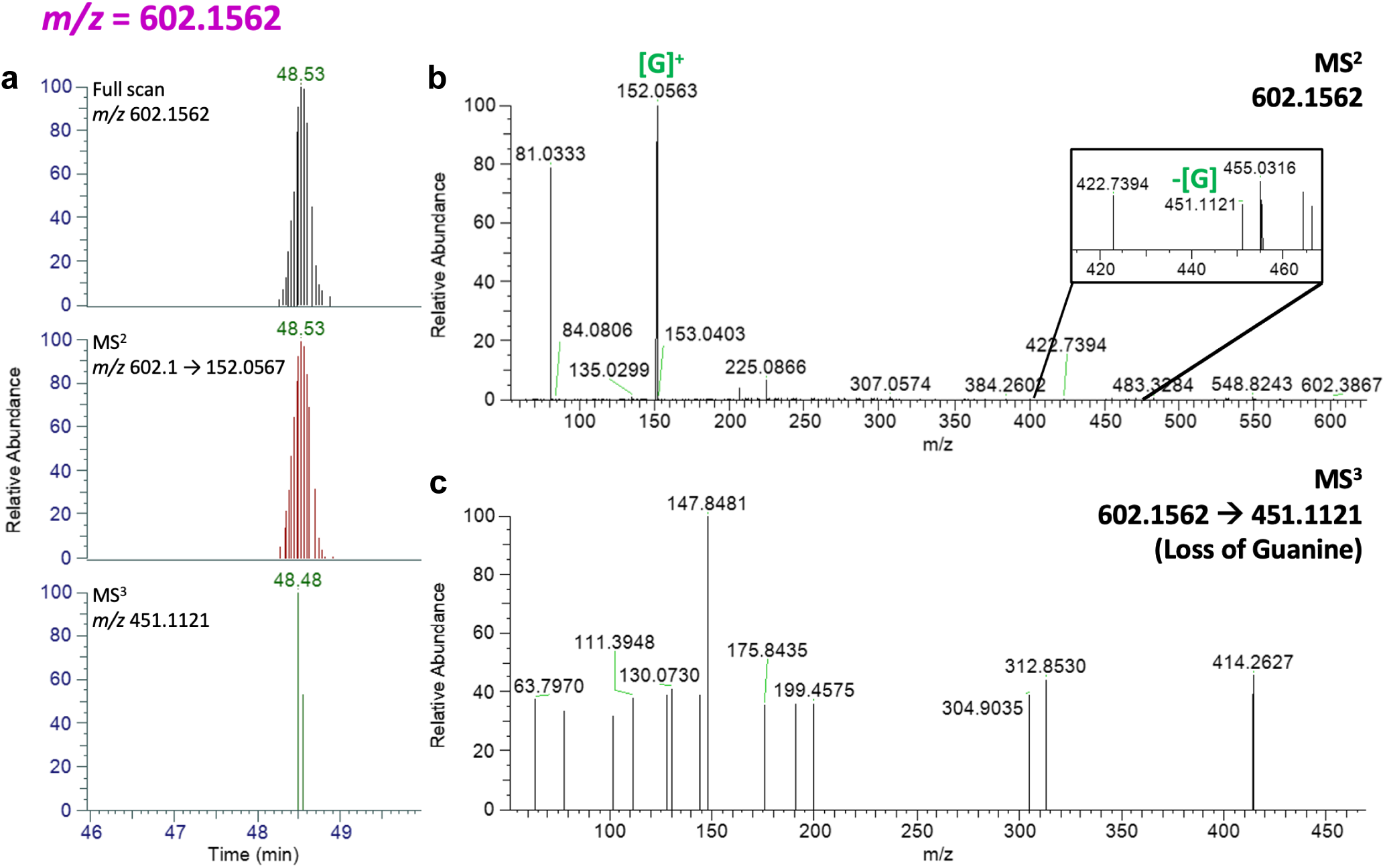
Mass Spectrometry data for the adduct with *m/z* 602.1562. **a)** Extracted chromatogram for the adduct in the full scan and the co-eluting peaks corresponding to the fragments detected in the MS^2^ and the MS^3^ fragmentation events. **b)** Spectrum corresponding to the MS^2^ fragmentation event with masses of known fragments highlighted. **c)** Spectrum of the MS^3^ fragmentation event triggered by the loss of Guanine in the MS^2^ fragmentation event.

**Supplementary Figure 15.**
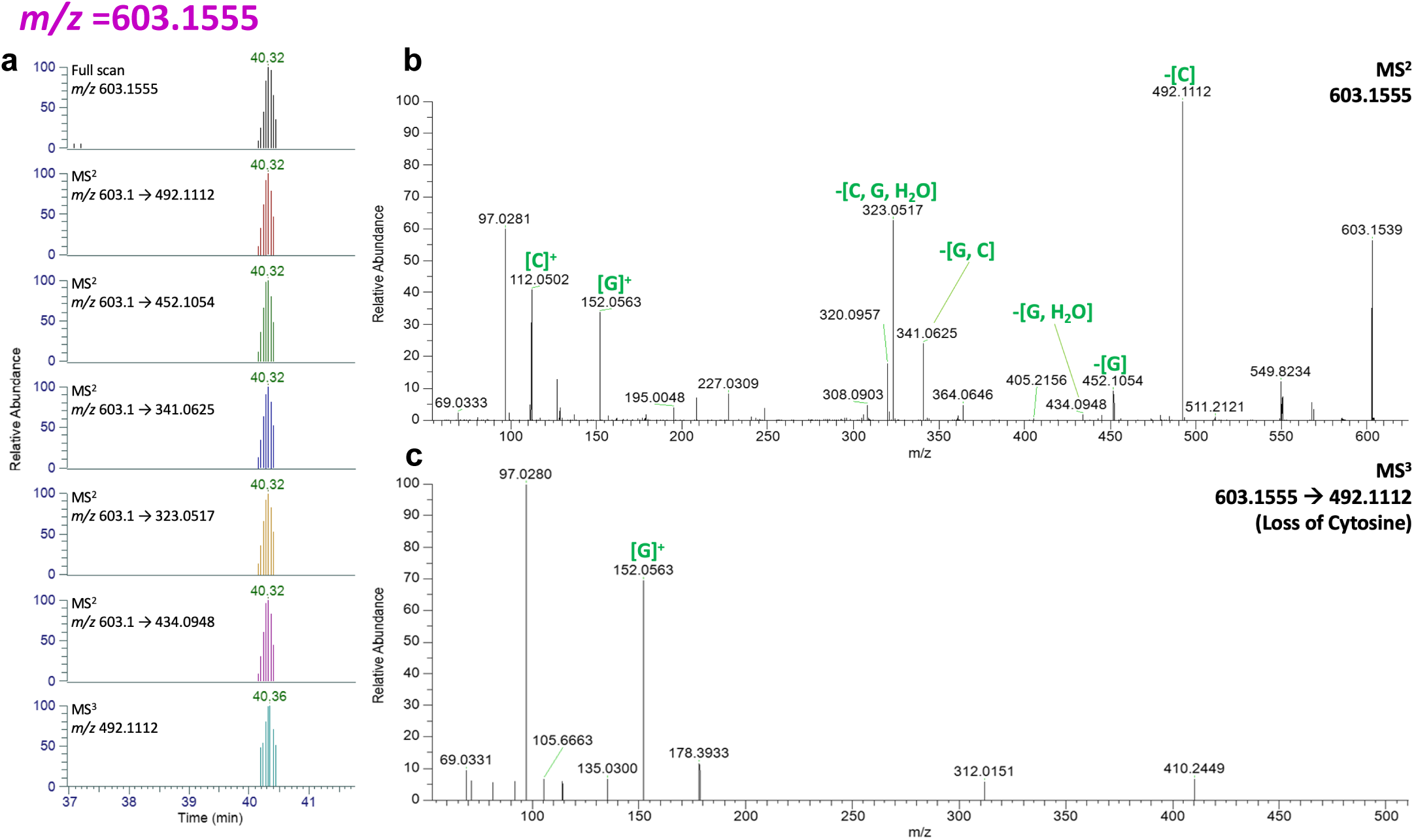
Mass Spectrometry data for the adduct with *m/z* 603.1555. **a)** Extracted chromatogram for the adduct in the full scan and the co-eluting peaks corresponding to the fragments detected in the MS^2^ and the MS^3^ fragmentation events. **b)** Spectrum corresponding to the MS^2^ fragmentation event with the masses of known fragments highlighted. **c)** Spectrum of the MS^3^ fragmentation event, with the appearance of Guanine, triggered by the loss of Cytosine in the MS^2^ fragmentation event.

**Supplementary Figure 16.**
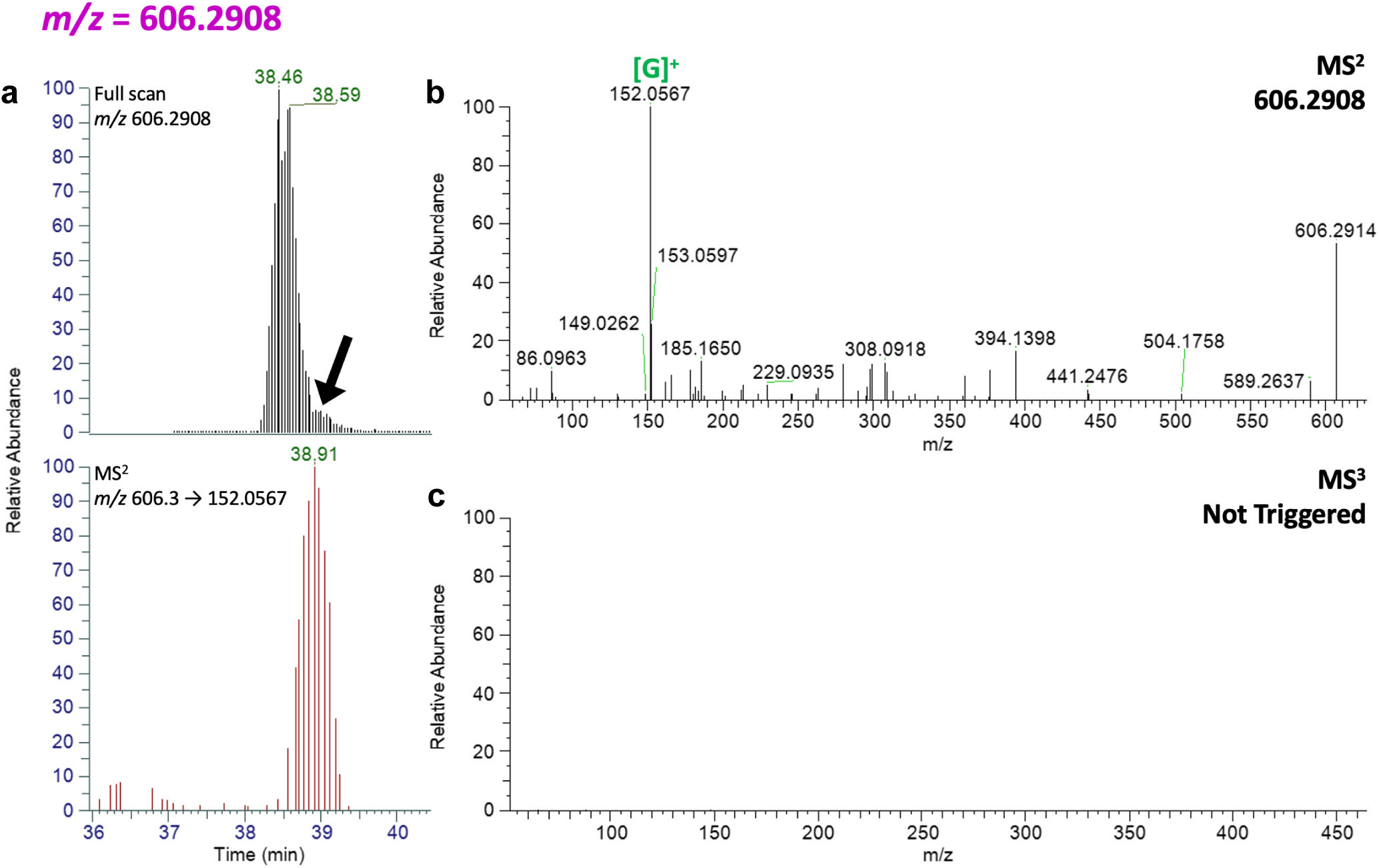
Mass Spectrometry data for the adduct with *m/z* 606.2908. **a)** Extracted chromatogram for the adduct in the full scan (the presence of a closely eluting peak with the same accurate mass reduces the ability to see a clear single peak) and the co-eluting peak corresponding to the fragment detected in the MS^2^ fragmentation event. **b)** Spectra corresponding to the MS^2^ fragmentation event with the mass of a known fragment highlighted. **c)** No MS^3^ fragmentation was triggered for this adduct.

**Supplementary Figure 17.**
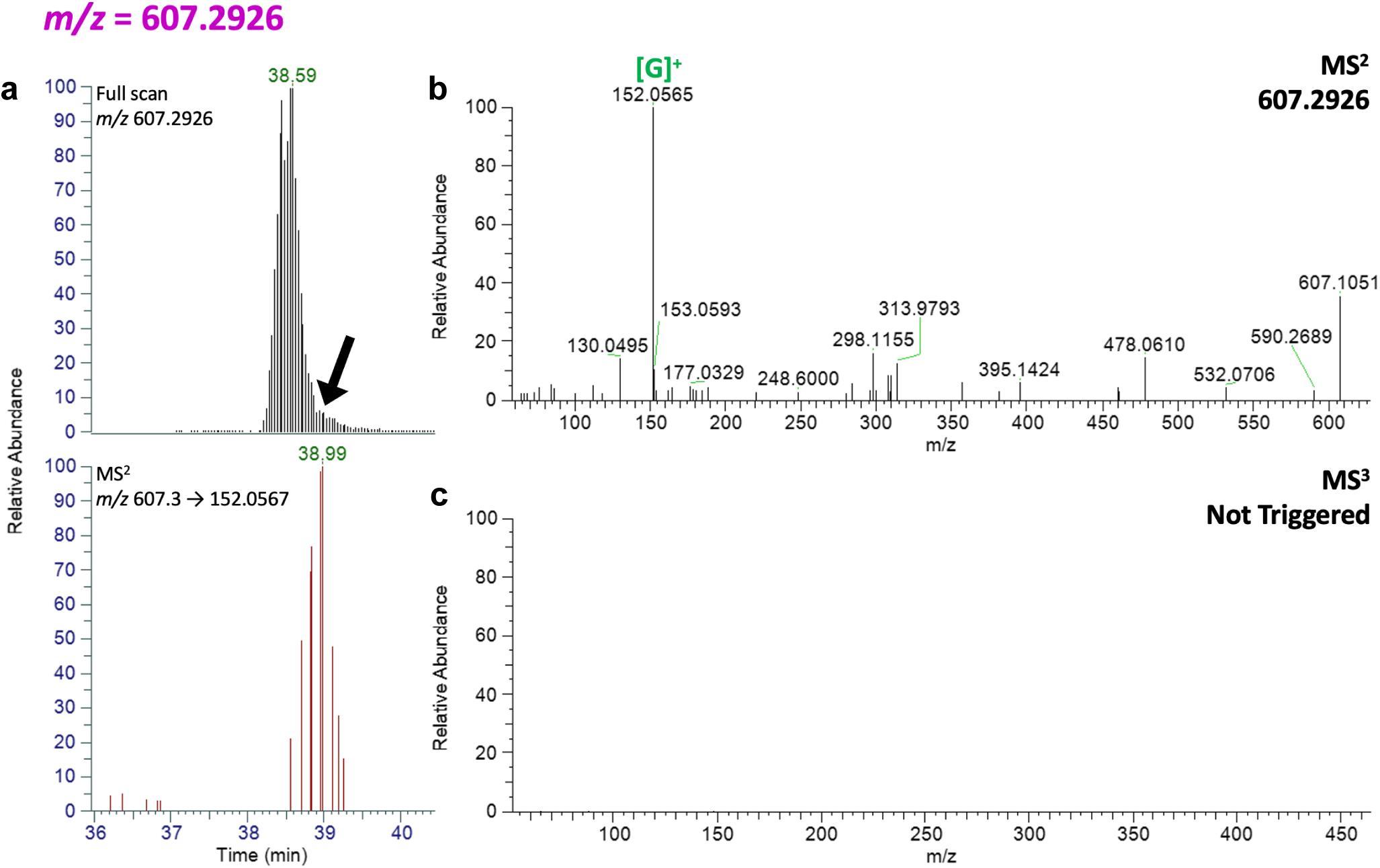
Mass Spectrometry data for the adduct with *m/z* 607.2926. **a)** Extracted chromatogram for the adduct in the full scan (the presence of a closely eluting peak with the same accurate mass reduces the ability to see a clear single peak) and the co-eluting peak corresponding to the fragment detected in the MS^2^ fragmentation event. **b)** Spectra corresponding to the MS^2^ fragmentation event with the mass of a known fragment highlighted. **c)** No MS^3^ fragmentation was triggered for this adduct.

## References

1. Lindahl, T. (1993). Instability and decay of the primary structure of DNA. Nature 362, 709–715. 10.1038/362709a0.

2. Tubbs, A., and Nussenzweig, A. (2017). Endogenous DNA Damage as a Source of Genomic Instability in Cancer. Cell 168, 644–656. 10.1016/j.cell.2017.01.002.

3. Garaycoechea, J.I., Crossan, G.P., Langevin, F., Daly, M., Arends, M.J., and Patel, K.J. (2012). Genotoxic consequences of endogenous aldehydes on mouse haematopoietic stem cell function. Nature 489, 571–575. 10.1038/nature11368.

4. Pilzecker, B., Buoninfante, O.A., and Jacobs, H. (2019). DNA damage tolerance in stem cells, ageing, mutagenesis, disease and cancer therapy. Nucleic Acids Res 47, 7163–7181. 10.1093/nar/gkz531.

5. Sale, J.E. (2013). Translesion DNA synthesis and mutagenesis in eukaryotes. Cold Spring Harb Perspect Biol 5, a012708. 10.1101/cshperspect.a012708.

6. Marteijn, J.A., Lans, H., Vermeulen, W., and Hoeijmakers, J.H.J. (2014). Understanding nucleotide excision repair and its roles in cancer and ageing. Nat Rev Mol Cell Biol 15, 465–481. 10.1038/nrm3822.

7. Gratchev, A., Strein, P., Utikal, J., and Sergij, G. (2003). Molecular genetics of Xeroderma pigmentosum variant. Exp Dermatol 12, 529–536. 10.1034/j.1600-0625.2003.00124.x.

8. Yurchenko, A.A., Rajabi, F., Braz-Petta, T., Fassihi, H., Lehmann, A., Nishigori, C., Wang, J., Padioleau, I., Gunbin, K., Panunzi, L., et al. (2023). Genomic mutation landscape of skin cancers from DNA repair-deficient xeroderma pigmentosum patients. Nat Commun 14, 2561. 10.1038/s41467-023-38311-0.

9. Alexandrov, L.B., Kim, J., Haradhvala, N.J., Huang, M.N., Tian Ng, A.W., Wu, Y., Boot, A., Covington, K.R., Gordenin, D.A., Bergstrom, E.N., et al. (2020). The repertoire of mutational signatures in human cancer. Nature 578, 94–101. 10.1038/s41586-020-1943-3.

10. Alexandrov, L.B., Nik-Zainal, S., Wedge, D.C., Aparicio, S.A.J.R., Behjati, S., Biankin, A. V., Bignell, G.R., Bolli, N., Borg, A., Børresen-Dale, A.L., et al. (2013). Signatures of mutational processes in human cancer. Nature 500, 415–421. 10.1038/nature12477.

11. Alexandrov, L.B., Jones, P.H., Wedge, D.C., Sale, J.E., Campbell, P.J., Nik-Zainal, S., and Stratton, M.R. (2015). Clock-like mutational processes in human somatic cells. Nat Genet 47, 1402–1407. 10.1038/ng.3441.

12. Nik-Zainal, S., Alexandrov, L.B., Wedge, D.C., Van Loo, P., Greenman, C.D., Raine, K., Jones, D., Hinton, J., Marshall, J., Stebbings, L.A., et al. (2012). Mutational processes molding the genomes of 21 breast cancers. Cell 149, 979–993. 10.1016/j.cell.2012.04.024.

13. Kuijk, E., Jager, M., van der Roest, B., Locati, M.D., Van Hoeck, A., Korzelius, J., Janssen, R., Besselink, N., Boymans, S., van Boxtel, R., et al. (2020). The mutational impact of culturing human pluripotent and adult stem cells. Nat Commun 11. 10.1038/s41467-020-16323-4.

14. van den Boogaard, M.L., Oka, R., Hakkert, A., Schild, L., Ebus, M.E., van Gerven, M.R., Zwijnenburg, D.A., Molenaar, P., Hoyng, L.L., Dolman, M.E.M., et al. (2021). Defects in 8-oxo-guanine repair pathway cause high frequency of C > A substitutions in neuroblastoma. Proc Natl Acad Sci U S A 118. 10.1073/pnas.2007898118.

15. Jager, M., Blokzijl, F., Kuijk, E., Bertl, J., Vougioukalaki, M., Janssen, R., Besselink, N., Boymans, S., De Ligt, J., Pedersen, J.S., et al. (2019). Deficiency of nucleotide excision repair is associated with mutational signature observed in cancer. Genome Res 29, 1067– 1077. 10.1101/gr.246223.118.

16. Yurchenko, A.A., Padioleau, I., Matkarimov, B.T., Soulier, J., Sarasin, A., and Nikolaev, S. (2020). XPC deficiency increases risk of hematologic malignancies through mutator phenotype and characteristic mutational signature. Nat Commun 11. 10.1038/s41467-020-19633-9.

17. Campbell, P., Chapman, M.S., Mitchell, E., Yoshida, K., Williams, N., Fabre, M., Ranzoni, A.M., Robinson, P., Wilk, C.M., Boettcher, S., et al. (2023). Prolonged persistence of mutagenic DNA lesions in stem cells. Preprint, 10.21203/rs.3.rs-3610927/v1.

18. Stern, H.R., Sefcikova, J., Chaparro, V.E., and Beuning, P.J. (2019). Mammalian DNA Polymerase Kappa Activity and Specificity. Molecules 24. 10.3390/molecules24152805.

19. Abascal, F., Harvey, L.M.R., Mitchell, E., Lawson, A.R.J., Lensing, S. V., Ellis, P., Russell, A.J.C., Alcantara, R.E., Baez-Ortega, A., Wang, Y., et al. (2021). Somatic mutation landscapes at single-molecule resolution. Nature 593, 405–410. 10.1038/s41586-021-03477-4.

20. Behjati, S., Huch, M., Van Boxtel, R., Karthaus, W., Wedge, D.C., Tamuri, A.U., Martincorena, I., Petljak, M., Alexandrov, L. B., Gundem, G., et al. (2014). Genome sequencing of normal cells reveals developmental lineages and mutational processes. Nature 513, 422–425. 10.1038/nature13448.

21. Blokzijl, F., De Ligt, J., Jager, M., Sasselli, V., Roerink, S., Sasaki, N., Huch, M., Boymans, S., Kuijk, E., Prins, P., et al. (2016). Tissue-specific mutation accumulation in human adult stem cells during life. Nature 538, 260–264. 10.1038/nature19768.

22. Osorio, F.G., Rosendahl Huber, A., Oka, R., Verheul, M., Patel, S.H., Hasaart, K., de la Fonteijne, L., Varela, I., Camargo, F.D., and van Boxtel, R. (2018). Somatic Mutations Reveal Lineage Relationships and Age-Related Mutagenesis in Human Hematopoiesis. Cell Rep 25, 2308–2316.e4. 10.1016/j.celrep.2018.11.014.

23. Moore, L., Cagan, A., Coorens, T.H.H., Neville, M.D.C., Sanghvi, R., Sanders, M.A., Oliver, T.R.W., Leongamornlert, D., Ellis, P., Noorani, A., et al. (2021). The mutational landscape of human somatic and germline cells. Nature. 10.1038/s41586-021-03822-7.

24. Lee-Six, H., Olafsson, S., Ellis, P., Osborne, R.J., Sanders, M.A., Moore, L., Georgakopoulos, N., Torrente, F., Noorani, A., Goddard, M., et al. (2019). The landscape of somatic mutation in normal colorectal epithelial cells. Nature 574, 532–537. 10.1038/s41586-019-1672-7.

25. Liu, M., Wu, Y., Jiang, N., Boot, A., and Rozen, S.G. (2023). mSigHdp: hierarchical Dirichlet process mixture modeling for mutational signature discovery. NAR Genom Bioinform 5, lqad005. 10.1093/nargab/lqad005.

26. Lodato, M. a., Rodin, R.E., Bohrson, C.L., Coulter, M.E., Barton, A.R., Kwon, M., Sherman, M. a., Vitzthum, C.M., Luquette, L.J., Yandava, C.N., et al. (2018). Aging and neurodegeneration are associated with increased mutations in single human neurons. Science 359, 555–559. 10.1126/science.aao4426.

27. Luquette, L.J., Miller, M.B., Zhou, Z., Bohrson, C.L., Zhao, Y., Jin, H., Gulhan, D., Ganz, J., Bizzotto, S., Kirkham, S., et al. (2022). Single-cell genome sequencing of human neurons identifies somatic point mutation and indel enrichment in regulatory elements. Nat Genet 54, 1564–1571. 10.1038/s41588-022-01180-2.

28. Christensen, S., Van der Roest, B., Besselink, N., Janssen, R., Boymans, S., Martens, J.W.M., Yaspo, M., Priestley, P., Kuijk, E., Cuppen, E., et al. (2019). 5-Fluorouracil treatment induces characteristic T to G mutations in human cancer. Nat Commun 10, 4571. 10.1038/s41467-019-12594-8.

29. Sikkema, L., Ramírez-Suástegui, C., Strobl, D.C., Gillett, T.E., Zappia, L., Madissoon, E., Markov, N.S., Zaragosi, L.E., Ji, Y., Ansari, M., et al. (2023). An integrated cell atlas of the lung in health and disease. Nat Med 29, 1563–1577. 10.1038/s41591-023-02327-2.

30. Morganella, S., Alexandrov, L.B., Glodzik, D., Zou, X., Davies, H., Staaf, J., Sieuwerts, A.M., Brinkman, A.B., Martin, S., Ramakrishna, M., et al. (2016). The topography of mutational processes in breast cancer genomes. Nat Commun 7, 1–11. 10.1038/ncomms11383.

31. Otlu, B., Díaz-Gay, M., Vermes, I., Bergstrom, E.N., Zhivagui, M., Barnes, M., and Alexandrov, L.B. (2023). Topography of mutational signatures in human cancer. Cell Rep 42. 10.1016/j.celrep.2023.112930.

32. Koren, A., Polak, P., Nemesh, J., Michaelson, J.J., Sebat, J., Sunyaev, S.R., and McCarroll, S.A. (2012). Differential relationship of DNA replication timing to different forms of human mutation and variation. Am J Hum Genet 91, 1033–1040. 10.1016/j.ajhg.2012.10.018.

33. Stamatoyannopoulos, J.A., Adzhubei, I., Thurman, R.E., Kryukov, G. V, Mirkin, S.M., and Sunyaev, S.R. (2009). Human mutation rate associated with DNA replication timing. Nat Genet 41, 393–395. 10.1038/ng.363.

34. Daigaku, Y., Davies, A.A., and Ulrich, H.D. (2010). Ubiquitin-dependent DNA damage bypass is separable from genome replication. Nature 465, 951–955. 10.1038/nature09097.

35. Lang, G.I., and Murray, A.W. (2011). Mutation rates across budding yeast chromosome VI are correlated with replication timing. Genome Biol Evol 3, 799–811. 10.1093/gbe/evr054.

36. Waters, L.S., and Walker, G.C. (2006). The critical mutagenic translesion DNA polymerase Rev1 is highly expressed during G(2)/M phase rather than S phase. Proc Natl Acad Sci U S A 103, 8971–8976. 10.1073/pnas.0510167103.

37. Haradhvala, N.J., Polak, P., Stojanov, P., Covington, K.R., Shinbrot, E., Hess, J.M., Rheinbay, E., Kim, J., Maruvka, Y.E., Braunstein, L.Z., et al. (2016). Mutational Strand Asymmetries in Cancer Genomes Reveal Mechanisms of DNA Damage and Repair. Cell 164, 538–549. 10.1016/j.cell.2015.12.050.

38. Tomkova, M., Tomek, J., Kriaucionis, S., and Schuster-Böckler, B. (2018). Mutational signature distribution varies with DNA replication timing and strand asymmetry. Genome Biol 19, 129. 10.1186/s13059-018-1509-y.

39. Letouzé, E., Shinde, J., Renault, V., Couchy, G., Blanc, J.F., Tubacher, E., Bayard, Q., Bacq, D., Meyer, V., Semhoun, J., et al. (2017). Mutational signatures reveal the dynamic interplay of risk factors and cellular processes during liver tumorigenesis. Nat Commun 8. 10.1038/s41467-017-01358-x.

40. Hu, J., Adar, S., Selby, C.P., Lieb, J.D., and Sancar, A. (2015). Genome-wide analysis of human global and transcription-coupled excision repair of UV damage at single-nucleotide resolution. Genes Dev 29, 948–960. 10.1101/gad.261271.115.

41. Suzuki, N., Ohashi, E., Kolbanovskiy, A., Geacintov, N.E., Grollman, A.P., Ohmori, H., and Shibutani, S. (2002). Translesion synthesis by human DNA polymerase kappa on a DNA template containing a single stereoisomer of dG-(+)- or dG-(-)-anti-N(2)-BPDE (7,8-dihydroxy-anti-9,10-epoxy-7,8,9,10-tetrahydrobenzo[a]pyrene). Biochemistry 41, 6100–6106. 10.1021/bi020049c.

42. Choi, J.-Y., Angel, K.C., and Guengerich, F.P. (2006). Translesion synthesis across bulky N2-alkyl guanine DNA adducts by human DNA polymerase kappa. J Biol Chem 281, 21062–21072. 10.1074/jbc.M602246200.

43. Oka, Y., Nakazawa, Y., Shimada, M., and Ogi, T. (2024). Endogenous aldehyde-induced DNA–protein crosslinks are resolved by transcription-coupled repair. Nat Cell Biol 26, 784–796. 10.1038/s41556-024-01401-2.

44. van Sluis, M., Yu, Q., van der Woude, M., Gonzalo-Hansen, C., Dealy, S.C., Janssens, R.C., Somsen, H.B., Ramadhin, A.R., Dekkers, D.H.W., Wienecke, H.L., et al. (2024). Transcription-coupled DNA–protein crosslink repair by CSB and CRL4CSA-mediated degradation. Nat Cell Biol 26, 770–783. 10.1038/s41556-024-01394-y.

45. Carnie, C.J., Acampora, A.C., Bader, A.S., Erdenebat, C., Zhao, S., Bitensky, E., van den Heuvel, D., Parnas, A., Gupta, V., D’Alessandro, G., et al. (2024). Transcription-coupled repair of DNA–protein cross-links depends on CSA and CSB. Nat Cell Biol 26, 797–810. 10.1038/s41556-024-01391-1.

46. Cheng, G., Guo, J., Wang, R., Yuan, J.-M., Balbo, S., and Hecht, S.S. (2023). Quantitation by Liquid Chromatography-Nanoelectrospray Ionization-High-Resolution Tandem Mass Spectrometry of Multiple DNA Adducts Related to Cigarette Smoking in Oral Cells in the Shanghai Cohort Study. Chem Res Toxicol 36, 305–312. 10.1021/acs.chemrestox.2c00393.

47. Jarosz, D.F., Godoy, V.G., Delaney, J.C., Essigmann, J.M., and Walker, G.C. (2006). A single amino acid governs enhanced activity of DinB DNA polymerases on damaged templates. Nature 439, 225–228. 10.1038/nature04318.

48. Yuan, B., You, C., Andersen, N., Jiang, Y., Moriya, M., O’Connor, T.R., and Wang, Y. (2011). The roles of DNA polymerases κ and ι in the error-free bypass of N2-carboxyalkyl-2’-deoxyguanosine lesions in mammalian cells. J Biol Chem 286, 17503–17511. 10.1074/jbc.M111.232835.

49. Wolfle, W.T., Johnson, R.E., Minko, I.G., Lloyd, R.S., Prakash, S., and Prakash, L. (2005). Human DNA polymerase iota promotes replication through a ring-closed minor-groove adduct that adopts a syn conformation in DNA. Mol Cell Biol 25, 8748–8754. 10.1128/MCB.25.19.8748-8754.2005.

50. Anderson, C.J., Talmane, L., Luft, J., Connelly, J., Nicholson, M.D., Verburg, J.C., Pich, O., Campbell, S., Giaisi, M., Wei, P.-C., et al. (2024). Strand-resolved mutagenicity of DNA damage and repair. Nature 630, 744–751. 10.1038/s41586-024-07490-1.

51. Stancel, J.N.K., McDaniel, L.D., Velasco, S., Richardson, J., Guo, C., and Friedberg, E.C. (2009). Polk mutant mice have a spontaneous mutator phenotype. DNA Repair (Amst) 8, 1355–1362. 10.1016/j.dnarep.2009.09.003.

52. Kokic, G., Chernev, A., Tegunov, D., Dienemann, C., Urlaub, H., and Cramer, P. (2019). Structural basis of TFIIH activation for nucleotide excision repair. Nat Commun 10, 2885. 10.1038/s41467-019-10745-5.

53. van den Heuvel, D., van der Weegen, Y., Boer, D.E.C., Ogi, T., and Luijsterburg, M.S. (2021). Transcription-Coupled DNA Repair: From Mechanism to Human Disorder. Trends Cell Biol 31, 359–371. 10.1016/j.tcb.2021.02.007.

54. Kose, C., Cao, X., Dewey, E.B., Malkoç, M., Adebali, O., Sekelsky, J., Lindsey-Boltz, L.A., and Sancar, A. (2024). Cross-species investigation into the requirement of XPA for nucleotide excision repair. Nucleic Acids Res 52, 677–689. 10.1093/nar/gkad1104.

55. Minko, I.G., Yamanaka, K., Kozekov, I.D., Kozekova, A., Indiani, C., O’Donnell, M.E., Jiang, Q., Goodman, M.F., Rizzo, C.J., and Lloyd, R.S. (2008). Replication bypass of the acrolein-mediated deoxyguanine DNA-peptide cross-links by DNA polymerases of the DinB family. Chem Res Toxicol 21, 1983–1990. 10.1021/tx800174a.

56. Inman, G.J., Wang, J., Nagano, A., Alexandrov, L.B., Purdie, K.J., Taylor, R.G., Sherwood, V., Thomson, J., Hogan, S., Spender, L.C., et al. (2018). The genomic landscape of cutaneous SCC reveals drivers and a novel azathioprine associated mutational signature. Nat Commun 9, 3667. 10.1038/s41467-018-06027-1.

57. Ganz, J., Luquette, L.J., Bizzotto, S., Miller, M.B., Zhou, Z., Bohrson, C.L., Jin, H., Tran, A. V, Viswanadham, V. V, McDonough, G., et al. (2024). Contrasting somatic mutation patterns in aging human neurons and oligodendrocytes. Cell 187, 1955–1970.e23. 10.1016/j.cell.2024.02.025.

58. Martín-Pardillos, A., Tsaalbi-Shtylik, A., Chen, S., Lazare, S., Van Os, R.P., Dethmers-Ausema, A., Fakouri, N.B., Bosshard, M., Aprigliano, R., Van Loon, B., et al. (2017). Genomic and functional integrity of the hematopoietic system requires tolerance of oxidative DNA lesions. Blood 130, 1523–1534. 10.1182/blood-2017-01-764274.

59. Mulderrig, L., Garaycoechea, J.I., Tuong, Z.K., Millington, C.L., Dingler, F.A., Ferdinand, J.R., Gaul, L., Tadross, J.A., Arends, M.J., O’Rahilly, S., et al. (2021). Aldehyde-driven transcriptional stress triggers an anorexic DNA damage response. Nature 600, 158–163. 10.1038/s41586-021-04133-7.

60. Dator, R.P., Murray, K.J., Luedtke, M.W., Jacobs, F.C., Kassie, F., Nguyen, H.D., Villalta, P.W., and Balbo, S. (2022). Identification of Formaldehyde-Induced DNA-RNA Cross-Links in the A/J Mouse Lung Tumorigenesis Model. Chem Res Toxicol 35, 2025–2036. 10.1021/acs.chemrestox.2c00206.

61. Wilson, M.R., Jiang, Y., Villalta, P.W., Stornetta, A., Boudreau, P.D., Carrá, A., Brennan, C.A., Chun, E., Ngo, L., Samson, L.D., et al. (2019). The human gut bacterial genotoxin colibactin alkylates DNA. Science 363. 10.1126/science.aar7785.

62. Guidolin, V., Li, Y., Jacobs, F.C., MacMillan, M.L., Villalta, P.W., Hecht, S.S., and Balbo, S. (2023). Characterization and quantitation of busulfan DNA adducts in the blood of patients receiving busulfan therapy. Mol Ther Oncolytics 28, 197–210. 10.1016/j.omto.2023.01.005.

63. Guidolin, V., Jacobs, F.C., MacMillan, M.L., Villalta, P.W., and Balbo, S. (2023). Liquid Chromatography-Mass Spectrometry Screening of Cyclophosphamide DNA Damage In Vitro and in Patients Undergoing Chemotherapy Treatment. Chem Res Toxicol 36, 1278–1289. 10.1021/acs.chemrestox.3c00008.

64. Schenten, D., Gerlach, V.L., Guo, C., Velasco-Miguel, S., Hladik, C.L., White, C.L., Friedberg, E.C., Rajewsky, K., and Esposito, G. (2002). DNA polymerase kappa deficiency does not affect somatic hypermutation in mice. Eur J Immunol 32, 3152–3160. 10.1002/1521-4141(200211)32:11<3152::AID-IMMU3152>3.0.CO;2-2.

65. de Vries, A., van Oostrom, C.T., Hofhuis, F.M., Dortant, P.M., Berg, R.J., de Gruijl, F.R., Wester, P.W., van Kreijl, C.F., Capel, P.J., van Steeg, H., et al. (1995). Increased susceptibility to ultraviolet-B and carcinogens of mice lacking the DNA excision repair gene XPA. Nature 377, 169–173. 10.1038/377169a0.

66. Cheo, D.L., Ruven, H.J.T., Meira, L.B., Hammer, R.E., Burns, D.K., Tappe, N.J., van Zeeland, A.A., Mullenders, L.H.F., and Friedberg, E.C. (1997). Characterization of defective nucleotide excision repair in XPC mutant mice. Mutat Res 374, 1–9. 10.1016/s0027-5107(97)00046-8.

67. Huch, M., Dorrell, C., Boj, S.F., van Es, J.H., Li, V.S.W., van de Wetering, M., Sato, T., Hamer, K., Sasaki, N., Finegold, M.J., et al. (2013). In vitro expansion of single Lgr5+ liver stem cells induced by Wnt-driven regeneration. Nature 494, 247–250. 10.1038/nature11826.

68. Broutier, L., Andersson-Rolf, A., Hindley, C.J., Boj, S.F., Clevers, H., Koo, B.-K., and Huch, M. (2016). Culture and establishment of self-renewing human and mouse adult liver and pancreas 3D organoids and their genetic manipulation. Nat Protoc 11, 1724–1743. 10.1038/nprot.2016.097.

69. Jager, M., Blokzijl, F., Sasselli, V., Boymans, S., Janssen, R., Besselink, N., Clevers, H., Van Boxtel, R., and Cuppen, E. (2018). Measuring mutation accumulation in single human adult stem cells by whole-genome sequencing of organoid cultures. Nat Protoc 13, 59–78. 10.1038/nprot.2017.111.

70. McQualter, J.L., Yuen, K., Williams, B., and Bertoncello, I. (2010). Evidence of an epithelial stem/progenitor cell hierarchy in the adult mouse lung. Proc Natl Acad Sci U S A 107, 1414–1419. 10.1073/pnas.0909207107.

71. Bartfeld, S., Bayram, T., van de Wetering, M., Huch, M., Begthel, H., Kujala, P., Vries, R., Peters, P.J., and Clevers, H. (2015). In vitro expansion of human gastric epithelial stem cells and their responses to bacterial infection. Gastroenterology 148, 126–136.e6. 10.1053/j.gastro.2014.09.042.

72. Sato, T., Vries, R.G., Snippert, H.J., van de Wetering, M., Barker, N., Stange, D.E., van Es, J.H., Abo, A., Kujala, P., Peters, P.J., et al. (2009). Single Lgr5 stem cells build crypt-villus structures in vitro without a mesenchymal niche. Nature 459, 262–265. 10.1038/nature07935.

73. Dingler, F.A., Wang, M., Mu, A., Millington, C.L., Oberbeck, N., Watcham, S., Pontel, L.B., Kamimae-Lanning, A.N., Langevin, F., Nadler, C., et al. (2020). Two Aldehyde Clearance Systems Are Essential to Prevent Lethal Formaldehyde Accumulation in Mice and Humans. Mol Cell 80, 996–1012.e9. 10.1016/j.molcel.2020.10.012.

74. Tischler, G., and Leonard, S. (2014). biobambam: tools for read pair collation based algorithms on BAM files. Source Code Biol Med 9, 13. 10.1186/1751-0473-9-13.

75. Bergstrom, E.N., Huang, M.N., Mahto, U., Barnes, M., Stratton, M.R., Rozen, S.G., and Alexandrov, L.B. (2019). SigProfilerMatrixGenerator: a tool for visualizing and exploring patterns of small mutational events. BMC Genomics 20, 685. 10.1186/s12864-019-6041-2.

76. Islam, S.M.A., Díaz-Gay, M., Wu, Y., Barnes, M., Vangara, R., Bergstrom, E.N., He, Y., Vella, M., Wang, J., Teague, J.W., et al. (2022). Uncovering novel mutational signatures by de novo extraction with SigProfilerExtractor. Cell genomics 2, None. 10.1016/j.xgen.2022.100179.

77. Díaz-Gay, M., Vangara, R., Barnes, M., Wang, X., Islam, S.M.A., Vermes, I., Duke, S., Narasimman, N.B., Yang, T., Jiang, Z., et al. (2023). Assigning mutational signatures to individual samples and individual somatic mutations with SigProfilerAssignment. Bioinformatics 39. 10.1093/bioinformatics/btad756.

78. Aloia, L., McKie, M.A., Vernaz, G., Cordero-Espinoza, L., Aleksieva, N.p, van den Ameele, J., Antonica, F., Font-Cunill, B., Raven, A., Aiese Cigliano, R., et al. (2019). Epigenetic remodelling licences adult cholangiocytes for organoid formation and liver regeneration. Nat Cell Biol 21, 1321–1333. 10.1038/s41556-019-0402-6.

79. Li, B., Qing, T., Zhu, J., Wen, Z., Yu, Y., Fukumura, R., Zheng, Y., Gondo, Y., and Shi, L. (2017). A Comprehensive Mouse Transcriptomic BodyMap across 17 Tissues by RNA-seq. Sci Rep 7, 4200. 10.1038/s41598-017-04520-z.

80. Seabold, S., and Perktold, J. (2010). Statsmodels: Econometric and Statistical Modeling with Python.

81. Sparks, J., and Walter, J.C. (2019). Extracts for Analysis of DNA Replication in a Nucleus-Free System. Cold Spring Harb Protoc 2019. 10.1101/pdb.prot097154.

82. Lebofsky, R., Takahashi, T., and Walter, J.C. (2009). DNA replication in nucleus-free Xenopus egg extracts. Methods Mol Biol 521, 229–252. 10.1007/978-1-60327-815-7_13.

83. Räschle, M., Knipscheer, P., Enoiu, M., Angelov, T., Sun, J., Griffith, J.D., Ellenberger, T.E., Schärer, O.D., and Walter, J.C. (2008). Mechanism of replication-coupled DNA interstrand crosslink repair. Cell 134, 969–980. 10.1016/j.cell.2008.08.030.

84. Hodskinson, M.R., Bolner, A., Sato, K., Kamimae-Lanning, A.N., Rooijers, K., Witte, M., Mahesh, M., Silhan, J., Petek, M., Williams, D.M., et al. (2020). Alcohol-derived DNA crosslinks are repaired by two distinct mechanisms. Nature 579, 603–608. 10.1038/s41586-020-2059-5.

85. Paiano, V., Maertens, L., Guidolin, V., Yang, J., Balbo, S., and Hecht, S.S. (2020). Quantitative Liquid Chromatography-Nanoelectrospray Ionization-High-Resolution Tandem Mass Spectrometry Analysis of Acrolein-DNA Adducts and Etheno-DNA Adducts in Oral Cells from Cigarette Smokers and Nonsmokers. Chem Res Toxicol 33, 2197–2207. 10.1021/acs.chemrestox.0c00223.

